# Medulloblastoma Forms Symbiotic Metabolic Partnerships with Macrophages to Establish Leptomeningeal Metastases

**DOI:** 10.64898/2026.07.26.740417

**Authors:** Vernon Fong, Michelle Ly, Anders W. Erickson, Namal Abeysundara, Liam Hendrikse, Randy Van Ommeren, Polina Balin, Jigyansa Mishra, Bryn Livingston, Patryk Skowron, Olga Sirbu, Ncedile Mankahla, Jiao Zhang, Cory Richman, Raul Suarez, Ning Huang, Hao Wang, Lei Qin, Tajana Douglas, Jonelle Pallotta, Esta Mak, Sachin A. Kumar, Akash K. Kaushik, Hieu Vu, Lauren Zacharias, Kelly Veerasammy, Yuki X. Chen, Oliver Ocsenas, Veronique Voisin, Farzan Taj, Daphne Koubourli, Victoria Dzieciol, Lele Xu, Madeline Harvey, Jerry J. Fan, David Przelicki, Andrea Yeh, Kaitlin Kharas, Alexandra Rasnitsyn, Evan Wang, Winnie Ong, Tyler Jubenville, Qi Yang, Xi Huang, Olivier Ayrault, Robert Wechsler-Reya, Sean E. Egan, David Largaespada, Ralph J. DeBerardinis, He Ye, Rinat Abzalimov, Lincoln Stein, David W. Ellison, Gary Bader, Calixto-Hope Lucas, Olivier Saulnier, David Shih, Jüri Reimand, Craig Daniels, Shubham Singh, Sameer Agnihotri, Jeremy N. Rich, Vijay Ramaswamy, Michael D. Taylor, Xiaochong Wu

## Abstract

Leptomeningeal metastases are the primary source of morbidity and mortality for pediatric medulloblastoma patients. Due to limited surgical sampling of metastases in patients, little is understood of the mechanisms of metastasis. Here, we identify biologically distinct quiescent small metastases (designated as micrometastases) and mitotically active larger metastases (macrometastases). Macrometastases are more metabolically active than micrometastases and contain higher levels of lipids, particularly cholesterol. Macrometastases secrete CXCL12, which attracts lipid-laden macrophages into the tumor. Lipid-laden macrophages upregulate the cholesterol transporter ABCG1, promoting the efflux of free cholesterol, which is then taken up by tumor cells via the HDL receptor SCARB1. CXCL12-driven macrophage recruitment and exogenous cholesterol are sufficient and necessary to drive progression of medulloblastoma leptomeningeal metastases in vivo. High fat diets drive metastatic progression in vivo. Dietary or pharmacological interventions targeting the CXCL12-SCARB1-cholesterol axis represent therapeutic strategies to either prevent or treat medulloblastoma leptomeningeal metastases.

## INTRODUCTION

Medulloblastoma, a pediatric embryonal cancer of the developing rhombic lip, is divided into four distinct molecular subgroups, largely based on presumed cells of origin in upper and lower rhombic lips [1–4]. Despite clinical and biological distinctions, all subgroups of medulloblastoma metastasize primarily to the leptomeninges [5–7]. Extra leptomeningeal metastases are rare outside of highly treated patients. The biological basis for leptomeningeal homing to the leptomeninges, and why some patients have metastases while others do not is largely cryptogenic. Around a third of patients harbor metastases at presentation, whereas metastases are nearly universal at recurrence for patients with Groups 3 and 4 medulloblastoma [8]. Craniospinal radiation and high dose chemotherapy as prevention or treatment of metastatic disease induce considerable morbidity and mortality for medulloblastoma patients [9]. Despite the overwhelming importance of the leptomeningeal metastases in the clinical course of medulloblastoma, the vast majority of the medulloblastoma literature focuses on the primary tumor, as surgical resection of leptomeningeal metastases is seldom clinically useful and only very rarely undertaken. Metastatic medulloblastoma cells are highly divergent in their biology from their matched primary tumor [10, 11], suggesting that targeted therapies developed using primary tumor models might not be effective in the metastatic compartment.

Risk factors for leptomeningeal dissemination include molecular subgroup (with Group 3 tumors most likely to metastasize, followed by Group 4, sonic hedgehog (Shh), and Wnt tumors) and specific tumor somatic mutations [12–14]. Therapeutic radiation to tumors may promote tumor necrosis and drive leptomeningeal dissemination [15]. The mode of leptomeningeal medulloblastoma dissemination may include shedding of cells directly into the cerebrospinal fluid followed by distal leptomeningeal adherence, or hematogenous spread of tumor cells followed by homing and vascular exit at leptomeninges [16]. Cellular populations in the leptomeninges can be recruited by the metastatic tumor cells and re-educated to provide growth factors to the metastatic tumor cells [17].

Here, we studied the life cycle of medulloblastoma metastases, revealing a metastatic hierarchy from quiescent micrometastases, a small minority of which gave rise to larger and more metabolically active macrometastases. The progression of micrometastases to macrometastases was driven by recruitment of lipid-laden macrophages through secretion of CXCL12 by metastatic cells. Recruited macrophages secreted lipids, particularly cholesterol, which are taken up by the macrometastases to drive further growth. High fat diets drove leptomeningeal dissemination in murine models of medulloblastoma, suggesting that dietary or pharmacological interventions interrupting CXCL12–lipid laden macrophage–cholesterol secretion axis could treat or prevent currently untreatable medulloblastoma metastases.

## RESULTS

### Medulloblastoma leptomeningeal metastases comprise morphologically and biologically distinct micrometastases and macrometastases

Leptomeningeal metastases are very rarely resected from human patients, as cytoreduction is not helpful, resection of thin adherent metastases from the surface of the nervous system readily causes neurological deficits, and the nature of the metastases is seldom a diagnostic dilemma [5,8]. As such, access to patient samples of metastatic medulloblastoma is highly limited. However, we collected autopsy materials from human medulloblastoma patients, as well as in genetically engineered mouse models (GEMMs) of medulloblastoma to determine the morphology of leptomeningeal metastases. We found frequent, very small metastatic deposits, which we designated as micrometastases (Figure 1, A and B). Medulloblastoma has been modeled with genetically engineered murine models with intact immune systems using cell type expression of the Sleeping Beauty transposon in conjunction with targeting of the *Ptch* tumor suppressor (*Math1-GFP/J2Q-SB11/T2Onc/Ptch*^+/−^) or overexpression of *Smoothened* (*Math1-GFP/Nestin-SB100/T2Onc2/SmoA1*^tg/tg^). Both models spontaneously develop GFP-labeled medulloblastoma with typical leptomeningeal metastases [10, 11]. After tumor development, we collected the central nervous systems of the animals and examined for the presence of metastases [17]. In parallel, we generated cohorts of immunodeficient mice bearing human patient-derived Group 3 medulloblastoma xenografts orthotopically (i.e., in the cerebellum). Tumor cell deposits in the leptomeninges were detected at the time of necropsy through either EGFP and LacZ labels in GEMMs and xenografts, respectively. Metastases were primarily dichotomized between two sizes: extremely small metastases that were only visible at high magnification (hereafter micrometastases) and larger metastases, some of which were visible with the naked eye (macrometastases) (Figure 1, C and D; see representative whole mount spine stereoscope and confocal images of D425 xenografts in Figure S1A - D). Micrometastases and macrometastases were consistently observed across species and models, whereas we seldom observed intermediate sized leptomeningeal metastases. We therefore hypothesized that the micrometastases are biologically distinct from the macrometastases, with only a small percentage of micrometastases progressing to macrometastatic size.

**Figure 1:**
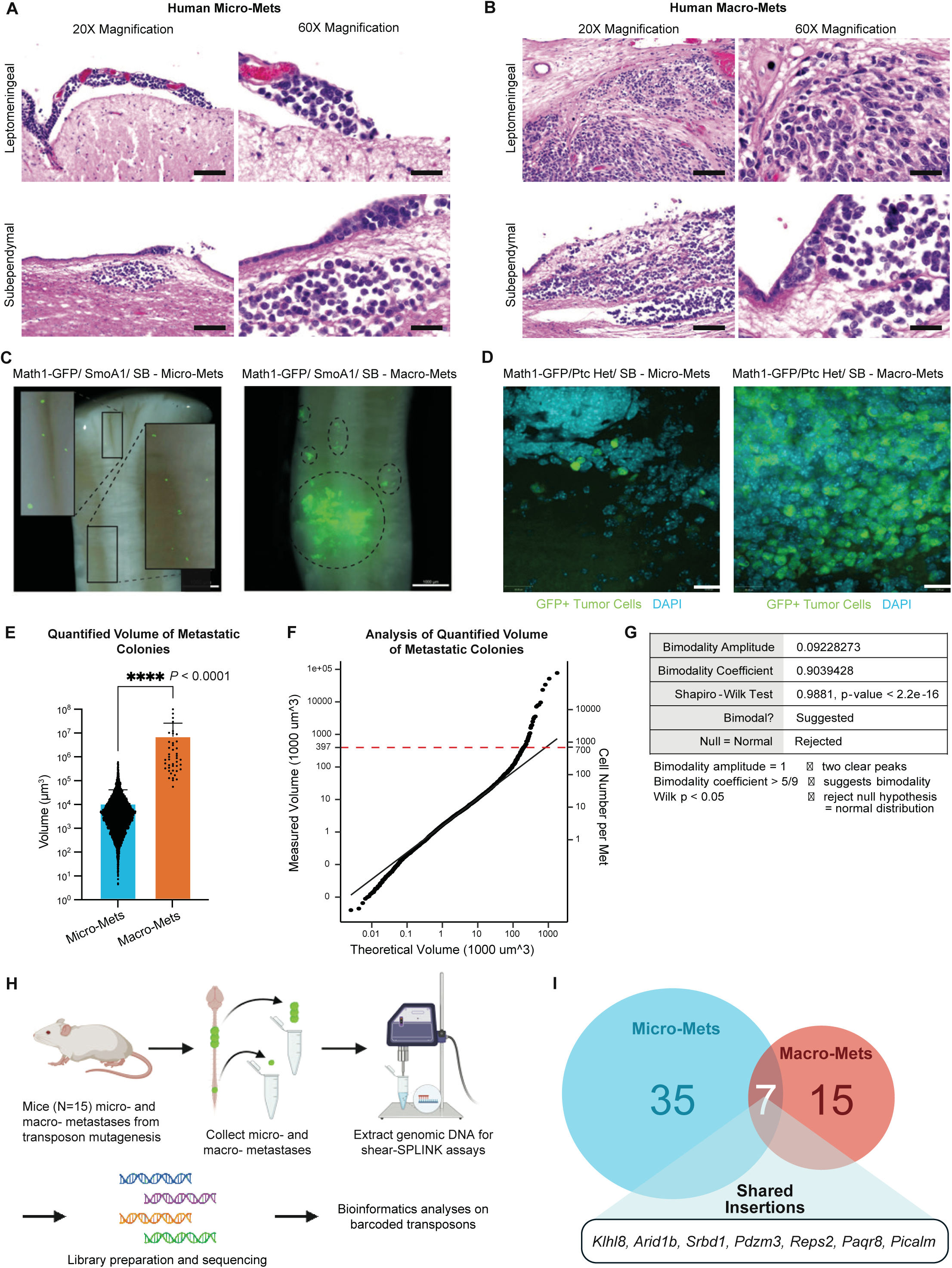
Medulloblastoma Induces Distinct Micro- and Macro-mets in the Leptomeninges. A) Hematoxylin and eosin (H&E) staining of both leptomeningeal and subependymal medulloblastoma metastases from a patient autopsy obtained from a 3-year-old male diagnosed with a posterior fossa medulloblastoma exhibiting large cell and anaplastic features. Representative 20X and 60X images of micro-mets. Each scale bar denotes 1000 µm for 20X images and 50 µm for 60X images. B) Macro-mets from the same patient. Each scale bar denotes 1000 µm for 20X images and 50 µm for 60X images. C) Fluorescence microscopy images of representative GFP-positive metastatic cells from SHH GEMM mouse models of medulloblastoma. These images illustrate both micro-mets and macro-mets adherent to the spinal cord leptomeninges in SmoA1 Sleeping Beauty mice. Macro-mets are delineated by dotted outlines. Each scale bar denotes 1000 µm. D) Confocal microscopy of whole-mounted microdissected leptomeninges from *Ptch*^+/−^ Sleeping Beauty mice, depicting both micro- and macro-mets. Each scale bar denotes 20 µm. E) Quantification of micro-met versus and macro-metastasis volumes, as measured from microdissected leptomeninges in *Ptch*^+/−^ Sleeping Beauty mice imaged using a Nikon A1R confocal microscope. Unpaired t-test, **** p-value < 0.0001. F) To delineate the difference in volume between micro-mets and macro-mets, we fitted a mixture of two Gaussian distributions on the volumetric measurements of metastases. The quantile-quantile plot showed that the first Gaussian captured the micro-mets adequately, and reveals the volumetric boundary between micro- and macro-mets. G) Statistical analysis of the bimodal distribution of micro- and macro-mets based on volume. Despite potential bimodal characteristics, the distribution appears normal due to the higher frequency of micro-mets, with a bimodality coefficient of 0.904. The Shapiro-Wilk test supports a non-normal distribution (p-value < 2.2 × 10^-16). H) Schematic representation of shear-SPLINK analysis employed to identify common and shared insertions from *Ptch*^+/−^ Sleeping Beauty mouse derived metastases. Proportions of shared insertions between primary tumors, macro-mets, and micro-mets were compared to assess similarities of clonal evolution pathways. I) Identification of specific shared genomic common insertion sites (gCISes) between micro- and macro-mets through shear-SPLINK analysis. Micro-mets contained 42 gCISes, while macro-mets harbored 22, suggesting greater clonal diversity in micro-mets compared to macro-mets.

To delineate the size threshold between micrometastases and macrometastases, we fitted a mixture of two Gaussian distributions to volumetric measurements of leptomeningeal metastases. The quantile-quantile plot showed that the first Gaussian captured the micro-metastases adequately and revealed the volumetric boundary between micro- and macro- metastases (Figure 1, E and F). The median volume of a micrometastasis was much smaller than that of a macrometastasis (2822 µm^3^ versus 396,000 µm^3^; Figure 1, E and F). We note that micrometastases are more than a hundred-fold more common than macrometastases in our model systems, suggesting that only a small minority of micrometastases progress to become macrometastases (Figure 1E).

The distribution of micro- and macrometastases suggested a bimodal distribution. We tested for characteristics of a bimodal distribution within our metastasis volumetric data. The bimodality amplitude determines the number of peaks, in which a value equivalent to 1 represents two clear, distinct peaks. Due to the large difference in the number of micrometastases compared to macrometastases, the bimodality amplitude was 0.092, indicative of one peak with a long right-sided tail (Figure 1G). However, the bimodality coefficient was 0.904, which is greater than 5/9 or 0.556, which supported a bimodal distribution (Figure 1G). Furthermore, a Shapiro-Wilk normality test result from a random sampling of 5000 volumetric data points was significant (p-value = 2.2 × 10^−16^) and rejected the null hypothesis for a normal distribution (Figure 1G). We conclude that micrometastases are morphologically distinct from macrometastases and do not likely represent a continuum of states.

We quantified the numbers of micrometastases and macrometastases by their constituent cell numbers per metastasis, surface area, and volume. The vast majority of micrometastases were single cells, with a small percentage of micrometastases comprised of clusters of fewer than 25 cells. The average number of cells was significantly different (p-value < 0.0001) between micrometastases and the macrometastases, which were 2 and 249 cells respectively. The average surface area of micrometastases was significantly smaller than macrometastases (74.01 µm^2^ versus 5690 µm^2^ (p-value < 0.0001)). The average volumes of micro- and macro-metastases were found to be 3040 µm^3^ versus 617,408 µm^3^ respectively (p-value < 0.0001). Our data are consistent with a model in which micrometastases are common with only a very small percentage progressing to form morphologically distinct macrometastases, suggesting that a biological event or switch is necessary for progression from micrometastasis to a macrometastatic state.

To test the hypothesis that micrometastases and macrometastases were genetically distinct, we interrogated our published functional genomic *in vivo* murine models of metastatic medulloblastoma driven by the Sleeping Beauty transposon system [10, 11]. We isolated primary tumors, micrometastases, and macrometastases from 15 mice with transposon-driven Shh medulloblastoma, then determined the patterns of transposon mobilization and clonal selection for each compartment (primary versus micrometastases versus macrometastases) (Figure 1H). While there was some overlap in the commonly inserted genes between micrometastases and macrometastases, each compartment contained unshared recurrent insertions (Figure 1I). Micrometastases contained a larger number of specific recurrent transposon insertions (n = 28) compared to macrometastases (n = 8), consistent with a model in which the micrometastases are more heterogeneous than the macrometastases, with only a few rare micrometastases undergoing progression to macrometastases (Figure 1I). This *in vivo* functional genomic screen supports the hypothesis that micrometastases are biologically, and perhaps genetically distinct from macrometastases, even within an individual animal.

### Micrometastases and macrometastases demonstrate distinct growth potential

Volumetric difference between micrometastases and macrometastases could arise due to either increased cell proliferation or decreased cell death (e.g., apoptosis) in the formation of macrometastases. Therefore, we administered EdU *in vivo* to *Math1-GFP/Nestin- SB100/T2Onc2/SmoA1*^tg/tg^ mice, then microdissected the leptomeninges and stained leptomeningeal sections for EdU to compare the rate of mitosis (cell cycle progression) between micrometastases and macrometastases (Figure 2, A and B). Macrometastases are composed of more mitotically active cells (Figure 2A-C). Indeed, the vast majority of micrometastases were not active in the cell cycle, consistent with these being lone, quiescent cells (Figure 2C). TUNEL (Terminal deoxynucleotidyl transferase-mediated dUTP nick-end labeling) staining of comparative micrometastases and macrometastases demonstrated a higher apoptotic index in macrometastases (Figure 2D-F). We conclude that macrometastases were larger than micrometastases, likely due to increased rate of cell cycle progression.

**Figure 2.**
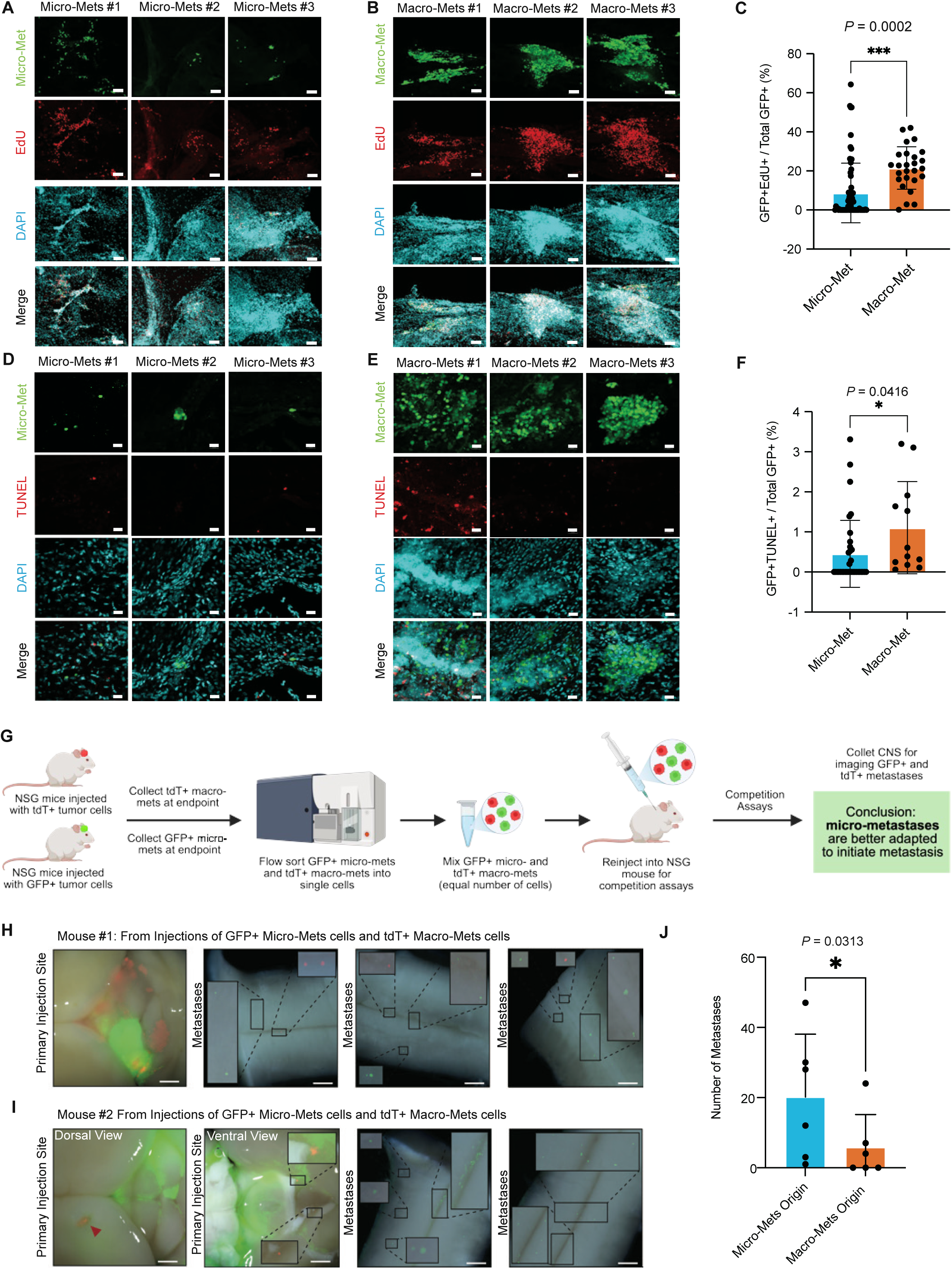
Micro-mets and Macro-mets Have Different Growth Potential. A) Confocal images of *in vivo* EdU labeled leptomeninges, demonstrating cell proliferation rates within micro-mets from SmoA1 Sleeping Beauty murine medulloblastomas. Each scale bar denotes 70 µm. B) Confocal images of *in vivo* EdU labeled leptomeninges, illustrating cell proliferation within macro-mets from SmoA1 Sleeping Beauty murine medulloblastomas. Each scale bar denotes 70 µm. C) Comparison of *in vivo* EdU incorporation in GFP-positive micro-mets versus macro-mets (N=3 mice; 65 micro-mets, 26 macro-mets). Unpaired t-test, *** P = 0.0002. D) Confocal images of TUNEL staining in micro-mets from SmoA1 Sleeping Beauty murine medulloblastomas. Each scale bar denotes 20 µm. E) Confocal images of TUNEL staining in macro-mets from SmoA1 Sleeping Beauty murine medulloblastomas. Each scale bar denotes 20 µm. F) Comparison of TUNEL staining in GFP-positive micro-mets versus macro-mets (N=3 mice; 34 micro-mets, 12 macro-mets). Unpaired t-test, * P = 0.0416. G) Experimental design for a two-color re-implantation assay to compare the ability of micro-mets versus macro-mets to found new metastases *in vivo*. Created in BioRender. Pomada Villalbi, A. (2025) https://BioRender.com/ms2fgjy H) Simultaneous orthotopic xenografting of GFP-positive micro-metastatic cells versus tdTomato- positive macro-metastatic cells in immunocompromised mice, leading to the formation of both GFP+ and tdTomato+ metastases. Scale bars: 1000 µm (first and second images); 500 µm (second, third and fourth images). I) Xenografting with equal numbers of GFP-positive micro-metastatic cells and tdTomato-positive macro-metastatic cells results in the almost exclusive formation of GFP-positive metastases, demonstrating a clear advantage for micro-mets to form nascent metastatic lesions. Scale bars: 1000 µm (first and second images); 500 µm (third and fourth images). J) Quantification of metastasis formation after simultaneous xenografting of equal numbers of GFP+ micro-metastatic versus tdTomato+ macro-metastatic cells in immunocompromised mice (N=6 mice). Wilcoxon test, * P = 0.0313.

The leptomeningeal space is a metabolic desert with very few nutrients available for growing cell populations due to the lack of capillary blood vessels [18, 19]. We hypothesized that the lone, quiescent micrometastatic tumor cells are better adapted to colonize the nutrient-poor environment of the leptomeninges. To test this hypothesis, we isolated EGFP^+ve^ tumor micrometastatic cells from immunodeficient mice xenografted with EGFP-labeled Group 3 medulloblastoma through flow cytometry. We similarly isolated macrometastatic tumor cells from mice xenografted with Group 3 medulloblastoma xenografts labelled with tdTomato (Figure 2G). We then mixed equal numbers of EGFP^+ve^ micrometastatic cells and tdTomato^+ve^ macrometastatic cells, then injected them into the lateral cerebral ventricle of secondary recipient immune deficient mice. Co-injection of equal numbers of green micrometastatic and red macrometastatic cells allowed us to test the relative efficiency of micrometastases versus macrometastases to colonize the leptomeninges and form new (secondary) micrometastases. Consistent with our hypothesis, most metastases observed in the secondary recipients were EGFP^+ve^ micrometastases (Figure 2H-J). These *in vivo* functional competition data were consistent with a model in which quiescent micrometastases were biologically distinct from macrometastases, and more capable of forming novel micrometastases on the surface of the leptomeninges.

### Macrometastases have a unique metabolic program

Medulloblastoma cells are known to exhibit metabolic programs distinct from those of healthy tissue [20]. Here, we asked whether micro-mets and macro-mets differ metabolically from one another. To address this, we performed a technique-demanding in vivo ^13^C-labeled glucose tracing assays which measures real-time glucose utilization by tracking the incorporation of ¹³C atoms from labeled glucose into downstream metabolites in living cells (Figure 3A). This approach allowed us to assess flux through glycolysis, the TCA cycle, and biosynthetic pathways directly within metastatic lesions.

**Figure 3.**
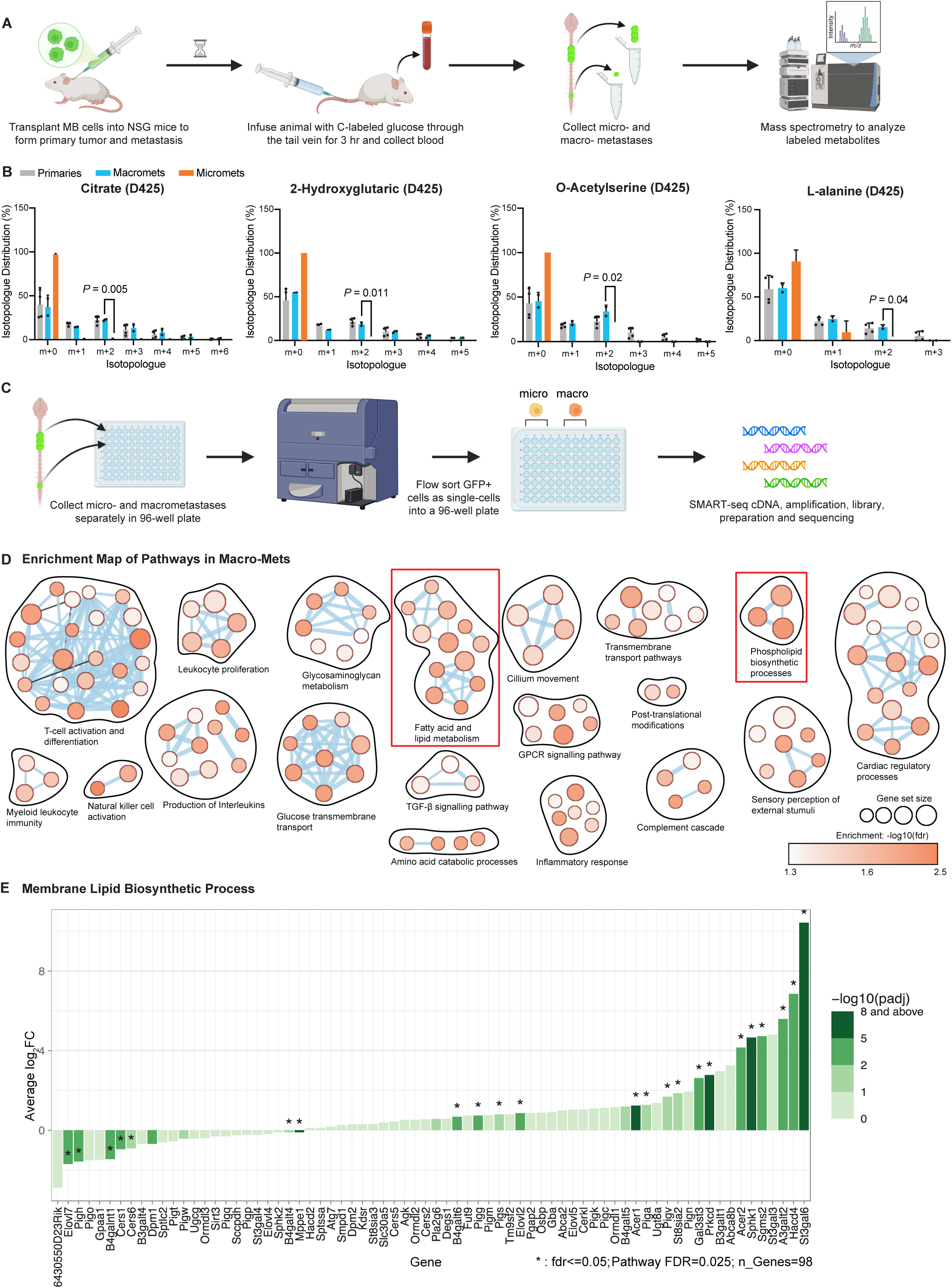
Macro-Metastasis Metabolism Differs from Micro-Metastasis *in vivo*. A) Schematic of medulloblastoma *in vivo* glucose tracing protocol using ^13^C-labeled glucose. Created in BioRender. Pomada Villalbi, A. (2025) https://BioRender.com/860vrll B) Bar plots illustrating *in vivo* glucose tracing results, with y-axis representing percentage of total metabolite pool in each isotopologue form, corrected for natural ^13^C abundance. Data are presented for primary tumors, micro-mets, and macro-mets (N=2 mice). Statistical significance was assessed using an unpaired t-test, with p-values comparing micro-mets and macro-mets. C) Protocol schematic of single-cell RNA sequencing to profile and contrast gene expression in micro- and macro-mets. D) Pathway enrichment analysis was conducted on the SMART-seq micro- and macro-mets, and an enrichment map of differentially expressed genes is presented. Notably, metabolic pathways are overrepresented in the macro-mets versus the micro-mets. E) Bar plots of critical membrane lipid biosynthesis genes contrasting macro-mets and micro- mets, including log2FC and adjusted p-values.

We found a striking metabolic divergence between micro- and macro-mets. Macrometastases contained ^13^C glucose labeling at comparable levels to the primary tumor (Figure 3B). In contrast, micrometastases displayed lower levels of ^13^C glucose integration in all interrogated metabolic pathways than macrometastases, supporting the hypothesis that micrometastases were less metabolically active than macrometastases. Micrometastases contained markedly lower ^13^C labeling of citrate compared to primary tumors and macrometastases, suggesting reduced TCA cycle activity and ongoing oxidative phosphorylation in micrometastases (Figure 3B). ^13^C labeling of L-alanine showed greater enrichment at the M+2 isotopologue in macrometastases, suggesting higher activity of the alanine synthesis pathway (Figure 3B; extended list of ^13^*C* glucose labelled metabolites shown in Figure S2A).

To determine the transcriptional regulation of metabolism, we compared single cell transcriptomes (SMART-seq) of micrometastases and macrometastases from *Math1- GFP/J2Q-SB11/T2Onc/Ptch*^+/−^ mice (Figure 3C). Pathway analysis of differentially expressed genes (DEGs) revealed remarkable convergence on metabolic pathways. Notable differential metabolic pathways included fatty acid and lipid metabolism and phospholipid synthetic processes (Figure 3D; full gene list in Supplemental Table 1). Membrane lipid biosynthesis, primarily sphingolipids, was also upregulated in macrometastases (Figure 3E; extended enrichment pathways shown in Figure S2B). Overall, micro- and macro-metastases differ in metabolism, with macrometastases being more metabolically active.

### Macrometastases contain high lipid levels

Based on the differential expression of lipid metabolism genes, we compared the lipidomes of spontaneous *Math1-GFP/J2Q-SB11/T2Onc/Ptch*^+/−^ micrometastases and macrometastases using untargeted shotgun lipidomics. Multiple lipid classes were different in abundance between micro- and macro-metastatic samples, including sterols, phosphosphingolipid, ceramides, and triacylglycerols (Figure 4, A and B; volcano and DAS plots for the untargeted lipidomics are provided in Figure S3A, B; see Supplemental Table 2 for full lipid classes). Fatty acids were found in relatively similar abundance in both metastatic compartments. To test the hypothesis that leptomeningeal micrometastases maintained their metabolism through fatty acid oxidation, we examined microdissected tumor bearing leptomeninges using matrix-assisted laser desorption ionization mass spectrometry imaging (MALDI IMS), which allowed us to determine the spatial distribution of distinct metabolites (Figure S3C, D). We detected acylcarnitines around micrometastases and hexadecanoic acid around macrometastases, with neither metabolite detected in normal, non-tumor bearing leptomeninges. Elevated levels of acylcarnitines indicate an active fatty acid oxidation in medulloblastoma micrometastases inhabiting the metabolic desert of the leptomeninges.

**Figure 4.**
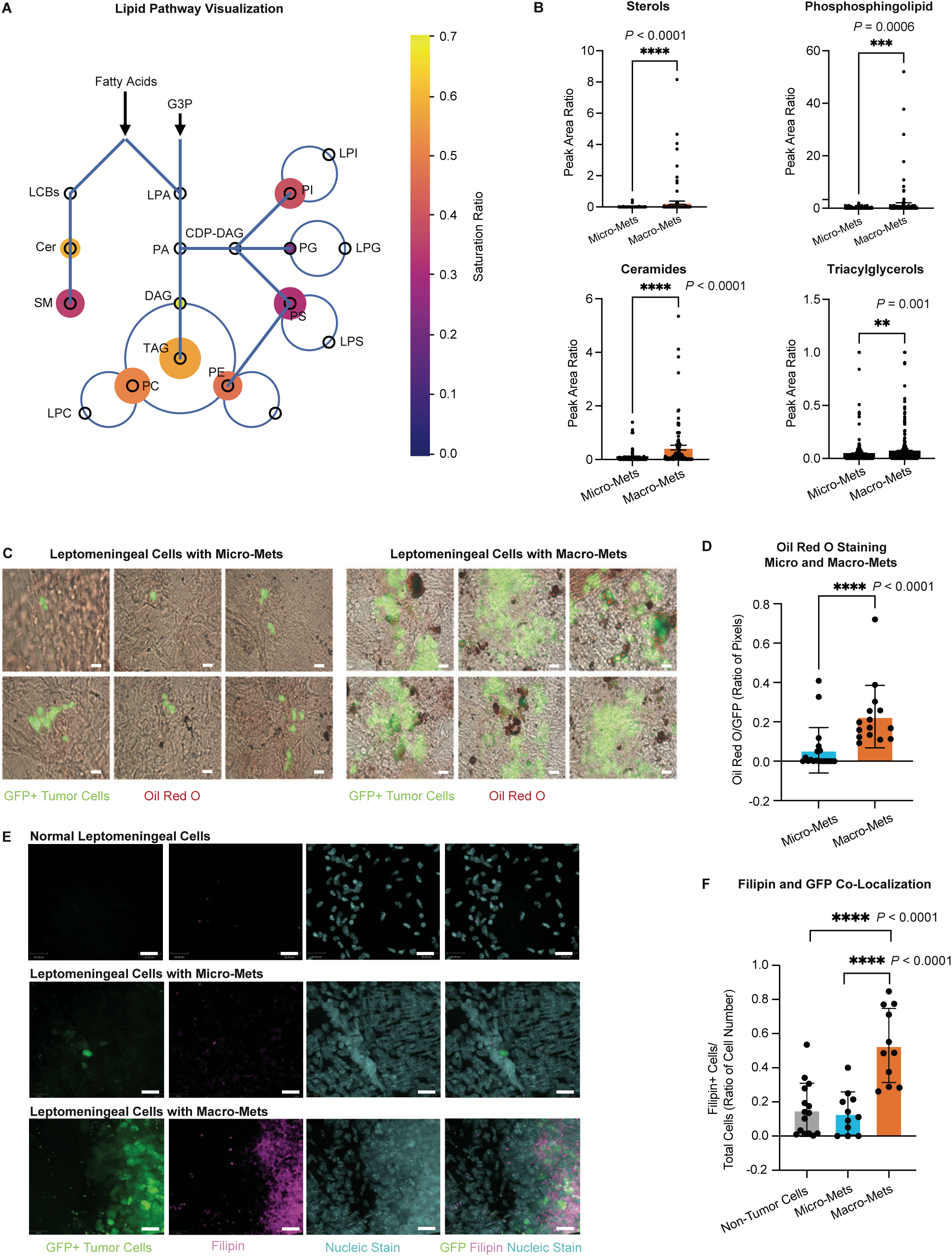
Lipid Metabolism Drives the Growth of Macro-mets. A) Pathway diagram generated using LipidCruncher [60] from untargeted shotgun lipidomics data. Nodes represent individual lipid classes. Node size represents relative abundance (larger circles denote enrichment in macro-mets compared to micro-mets). Node color indicates the saturation ratio of the fatty acids/lipids (warm colors denote higher saturation; empty nodes indicate unchanged lipid levels). Statistically significant lipid species are highlighted. Macro-mets exhibit coordinated enrichment of multiple lipid classes compared to micro-mets. B) Bar charts depicting differential abundance of key lipid classes between normal leptomeninges, micro-mets, and macro-mets (N=5 mice; Unpaired t-test, * P < 0.05, ** P < 0.01, *** P < 0.001, **** P < 0.0001). C) Oil Red O staining of neutral lipids in micro- and macro-mets from SmoA1 Sleeping Beauty spinal cord sections (N=3 mice; 114 micro-mets, 28 macro-mets). Each scale bar denotes 190 µm. D) Quantification of Oil Red O staining in micro-mets versus macro-mets (N=3 mice; 19 micro- mets, 15 macro-mets). Unpaired t-test, **** P < 0.0001. E) Filipin staining in microdissected leptomeningeal tissues from D425-GFPluc xenografts, indicating greater cholesterol accumulation in macro-mets relative to both micro-mets and normal leptomeningeal cells. Each scale bar denotes 20 µm. F) Quantification of Filipin staining, illustrating cholesterol accumulation in micro- and macro-mets compared to control leptomeningeal areas (N=3 nude mice; 15 control, 11 micro-mets, and 11 macro-mets images). Unpaired t-test, **** P < 0.0001.

We compared the availability of neutral lipids stored in lipid droplets between micro- and macro-metastases through Oil Red O (ORO) staining (Figure 4, C and D). Micrometastases had very low abundance of neutral lipids or lipid droplets as compared to macrometastases (Figure 4C). The average ORO positivity area around macrometastases was 285,086 pixels^2^ while micrometastasis was 17,489 pixels^2^. Furthermore, the detection of ORO staining for neutral lipids and lipid droplets showed that not only were neutral lipids accumulating in macrometastases tumor cells but also in the tumor microenvironment surrounding the metastasis (Figure 4C). ORO staining was then normalized to the area of tumor cells for standardized quantification (Figure 4D; Unpaired t-test, p-value < 0.0001), demonstrating markedly higher ORO staining in the macrometastases and surrounding microenvironment. Filipin3 staining for free cholesterol on leptomeninges from Group 3 D425 xenografted mice versus controls demonstrated high levels of cholesterol accumulation in tumor laden leptomeninges but not normal controls (Figure 4, E and F; additional high magnification filipin images are shown in Figure S4A). Cumulatively, our shotgun lipidomic analysis, ORO staining, and Filipin staining contrasting micro- and macrometastases support a model in which increased lipid synthesis drives the transition from micrometastasis to macrometastasis.

### Lipid metabolism is necessary for leptomeningeal metastasis formation in vivo

We have shown that macro-mets carry a higher level of lipids compared to micro-mets. To assess whether *de novo* cholesterol synthesis contributes to metastatic progression, we examined expression of key mevalonate pathway genes (*Hmgcr, Hmgcs1, Mvk, Sqle*) as well as the master regulator *Srebf2* in our SMART-seq dataset (see full DEG list in Supplemental Table 1). We did not observe enrichment of a coordinated *Srebf2* -driven transcriptional program, and expression of key mevalonate pathway genes was comparable between micro- and macro-mets, arguing against increased *de novo* cholesterol synthesis during metastatic progression.

We next tested the molecular mechanisms by which macro-mets uptake lipids during leptomeningeal metastasis. Several key mediators of lipid uptake in cells have been documented, these include, the SCARB1 (Scavenger Receptor Class B Member 1) membrane receptor crucial for selective uptake of lipids from High-Density Lipoprotein (HDL); the Low-Density Lipoprotein (LDL) Receptor for lipoproteins and chylomicron remnants; and the scavenger receptor CD36 for oxidized lipids and long-chain fatty acids [21]. LDL Receptor–Related Protein 1 (LRP1) is another endocytic receptor that binds lipoproteins and numerous ligands involved in lipid and protease metabolism [22]. LRP1 and CD36 have been reported to promote glioblastoma cell migration and invasion [23, 24]. Our SMART-seq data showed that compared to micro-mets, *Scarb1* transcript was expressed at a significantly higher level in the macro-mets from the *Math1-GFP/J2Q- SB11/T2Onc/Ptch*^+/−^ model, but not for *Ldlr, Lrp1*, and *Cd36* transcripts (Figure 5A; see violin plots of *Ldlr* and *Lrp1* in Figure S4B). Interestingly, *Cd36* transcripts were not expressed from either macro-met and micro-met samples. We performed immunofluorescence staining of SCARB1 on Group 3 medulloblastoma xenografts. The quantified results demonstrated that compared to micro-mets, SCARB1 protein accumulation was significantly higher in macro-mets (Figure 5, B and C; panels of expanded SCARB1 staining on primary tumor in Figure S4C).

**Figure 5.**
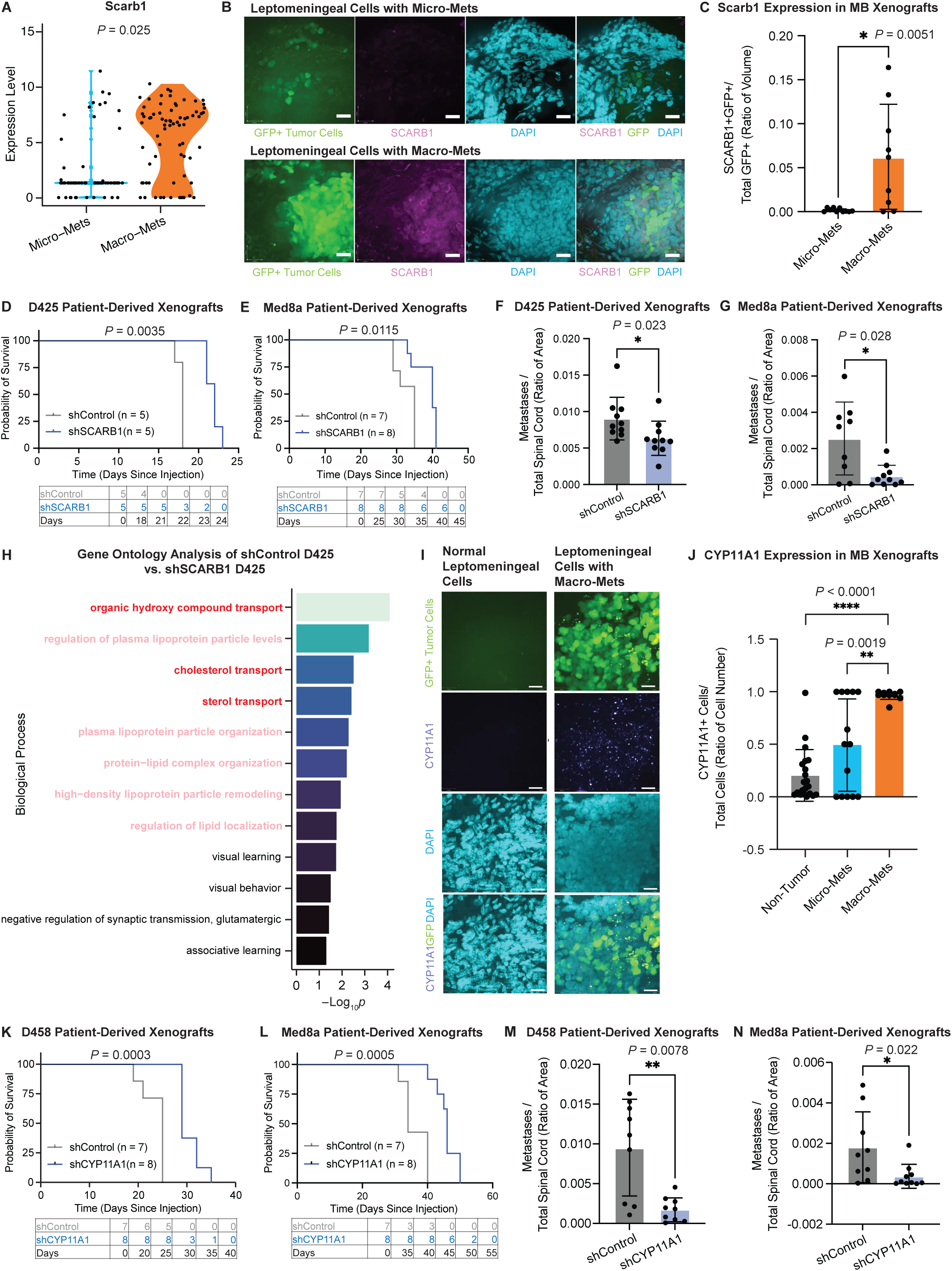
Inhibition of Lipid Metabolism Diminishes Leptomeningeal Metastases *in vivo*. A) Violin plots showing differential expression of *Scarb1* between micro- and macro-mets from Shh GEMM models, based on SMARTSEQ data (P = 0.025). B) Immunohistochemical analysis of SCARB1 accumulation in spinal meninges human Group 3 medulloblastoma xenografts demonstrating increased *SCARB1* accumulation in densely populated tumor regions. Each scale bar denotes 20 µm. C) Quantification of SCARB1 accumulation in densely versus sparsely populated tumor regions (N=3 nude mice; 10 micro-metastasis, 9 macro-metastasis images). Unpaired t-test, * P = 0.0051. D) Kaplan-Meier survival curve for shSCARB1 D425 and shControl D425 NSG xenografts, demonstrating prolonged survival of shSCARB1 D425 mice (P = 0.0035). E) Kaplan-Meier survival curve for shSCARB1 Med8a and shControl Med8a NSG xenografts, demonstrating prolonged survival of shSCARB1 Med8a mice (P = 0.0115). F) Mann-Whitney test of metastatic potential in shSCARB1 D425 versus shControl D425 NSG xenografts, indicating significantly reduced metastatic potential in shSCARB1-treated mice (N=10 shControl, 13 shSCARB1 mice; P = 0.023). G) Mann-Whitney test comparing the metastatic potential of shSCARB1 Med8a to shControl Med8a NSG xenografts demonstrates significantly lower metastatic potential in the shSCARB1 Med8a mice (N = 9 shControl mice, 10 shSCARB1 mice, P = 0.028). H) Investigation of the downstream mechanisms of increased cholesterol uptake was performed by conducting D425 xenograft experiments with and without SCARB1 shRNA. Upon reaching the endpoint (n = 3 per condition), primary tumors underwent bulk RNA-seq. Differential gene analysis and gene ontology were performed to identify differential pathway usage. Three significant pathways are highlighted in red, including organic hydroxy compound transport, cholesterol transport, and sterol transport. Five additional metabolism-related pathways are highlighted in pink. I) CYP11A1 staining was performed on spinal leptomeninges from two-month-old control nude mice versus leptomeningeal D425 patient-derived xenograft samples. Images were obtained using 63X spinning disk confocal microscopy. Each scale bar denotes 20 µm. J) A nested whisker plot displaying the percentage of CYP11A1-positive cells between micro- and macro-mets from D425 PDX metastases, versus control tumor free leptomeninges. (N = 3 nude mice, 21 control images, 14 micro-mets images, 11 macro-mets images, Unpaired t-test, **** P < 0.0001). K) Kaplan-Meier survival curve for shCYP11A1 D458 versus shControl D458 NSG xenografts, showing prolonged survival in shCYP11A1 D458 mice (P = 0.0003). L) Kaplan-Meier survival curve for shCYP11A1 Med8a versus shControl Med8a NSG xenografts, showing prolonged survival in shCYP11A1 Med8a mice (P = 0.0005). M) Mann-Whitney test comparing the metastatic potential of shCYP11A1 D458 versus shControl D458 NSG xenografts. CYP11A1 deficient medulloblastoma D425 cells have significantly lower metastatic potential as compared to controls (N = 12 shControl mice, 9 shCYP11A1 mice, P = 0.0078). N) Mann-Whitney test comparing the metastatic potential of shCYP11A1 Med8a versus shControl Med8a NSG xenografts. CYP11A1 deficient medulloblastoma Med8a cells have significantly lower metastatic potential as compared to controls (N = 10 shControl mice, 10 shCYP11A1 mice, P = 0.022).

To test the requirement of cholesterol in leptomeningeal dissemination of medulloblastoma, we knocked down *SCARB1* in Group 3 medulloblastoma cells lines D425 and MED8A, and in their respective xenograft models (Figure 5, D-G). *SCARB1* deficiency increased the survival of tumor-bearing mice and decreased metastatic burden for both D425 and MED8A, indicating that cholesterol is required for Group 3 medulloblastoma metastatic progression (Figure 5, D-G; knockdown validation by WB and representative whole mount spine images are provided in Figure S4D and Figure S5A, B).

To ascertain molecular mechanisms underlying the promotion of metastatic progression by SCARB1 and cholesterol, we performed bulk RNA sequencing on *SCARB1* knockdown primary tumors versus control D425 primary tumors at xenograft endpoint. Pathway enrichment analysis of differentially expressed genes revealed numerous metabolic pathways, many of which focused on cholesterol metabolism (Figure 5H). Unexpectedly, the most differentially expressed pathway was ‘organic hydroxy compound transport’ (Figure 5H; volcano plot from the DEGs list is in Figure S6A; see Supplemental Table 3 for full gene list), suggesting possible disruption of steroid hormone transport secondary to low intracellular cholesterol levels. *CYP11A1*, a key enzyme in steroidogenesis, was upregulated in tumors compared to normal leptomeninges and upregulated along the progression from micrometastases to macrometastases (Figure 5, I and J; CYP11A1 feature plots in scRNA data from Abeysundara et al., 2025 [17] in Figure S6C). To discern correlative versus causative effects of CYP11A1, we knocked down *CYP11A1* in both D425 and MED8A cell lines and then implanted the cells into immunocompromised hosts, resulting in longer median survival of xenografted mice *in vivo* and diminished metastatic burden (Figure 5K-N; representative whole mount spine images and knockdown validation IF are in Figure S5C, D and Figure S6B). These data support a causative driver role for cholesterol metabolism, SCARB1 expression, and possibly cholesterol dependent steroidogenesis mediated by CYP11A1 in the progression and maintenance of medulloblastoma leptomeningeal metastases.

### Medulloblastoma metastases contain lipid-laden macrophages

The source of cholesterol observed within leptomeningeal metastases and surrounding tumor cells in the leptomeningeal microenvironment as shown by ORO staining (Figure 4C) was cryptogenic, particularly in light of the limiting metabolic desert of the leptomeninges. Our prior work on medulloblastoma leptomeningeal metastases demonstrated that metastatic medulloblastoma cells can recruit and retain fibroblasts from the leptomeningeal microenvironment, resulting in the promotion of tumor growth by fibroblast secreted growth factors [17]. To determine the identity of microenvironment cells surrounding leptomeningeal medulloblastoma, we used SLP-mCherry technology [25]. Briefly, Group 3 medulloblastoma cells that expressed GFP-luciferase were transduced with SLP-mCherry virus. Transduced cells secreted the SLP-mCherry protein into their immediate microenvironment, where it was taken up by adjacent cell types (Figure 6A). D425 GFP-luc SLP-mCherry cells were xenografted into immunodeficient mice, which were allowed to come to neurologic endpoints followed by harvesting and microdissection of the leptomeninges. FACS was used to isolate cells from the leptomeninges that were mCherry^+ve^ but EGFP^−ve^, which were presumably microenvironment cells adjacent to tumor cells, then underwent single cell transcriptional profiling using SMART-seq (Figure 6B; representative immunofluorescence images for *mCherry+ve/EGFP-ve* cells in Figure S7A).

**Figure 6.**
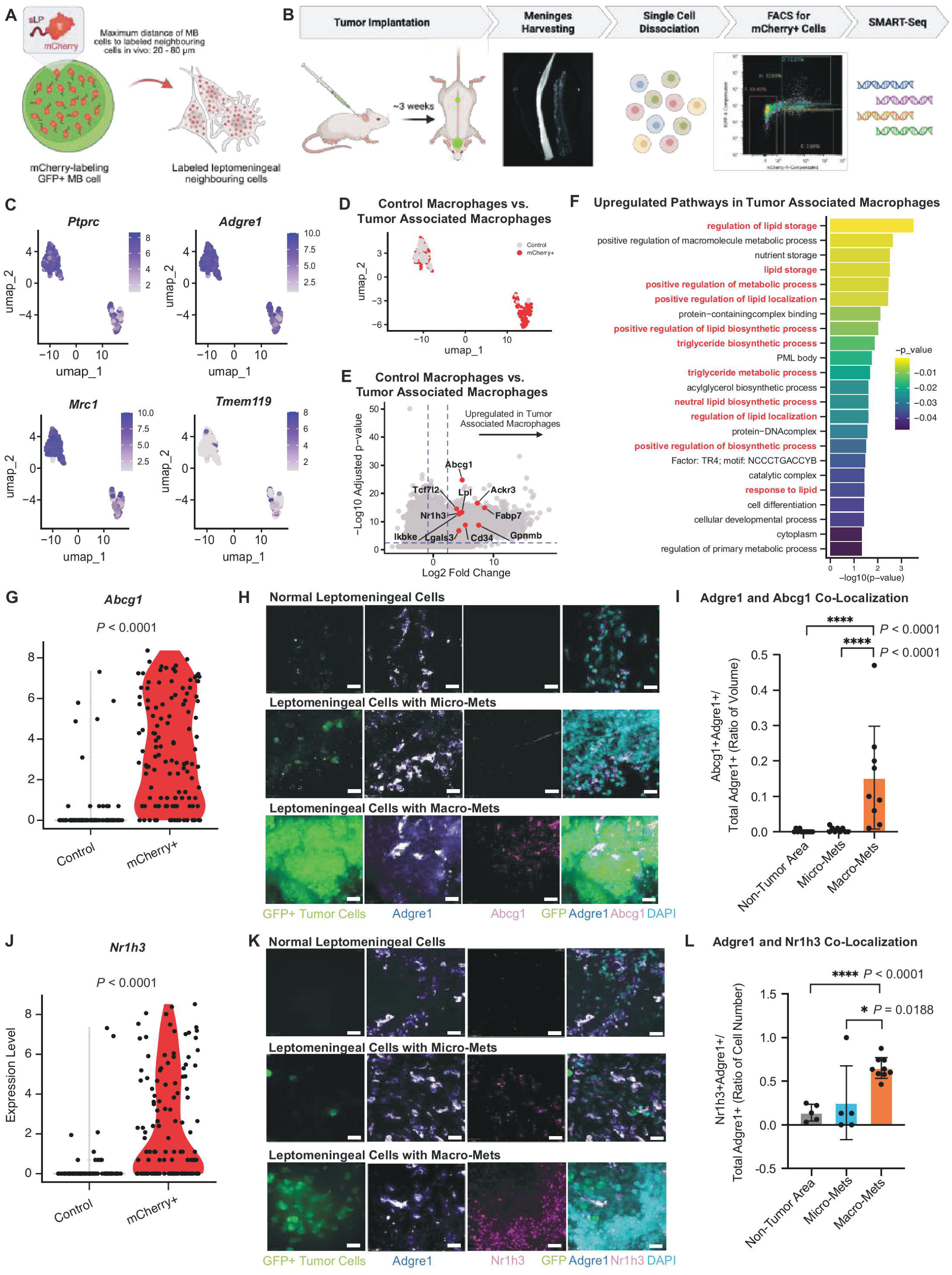
Lipid-Laden Macrophages Are Necessary to Drive Medulloblastoma Metastases. A) Schematic representation illustrating the sLP mCherry Niche methodology. EGFP^+ve^ tumor cells infected with sLP mCherry, express both GFP and mCherry. The sLP mCherry protein is subsequently secreted and taken up by directly adjacent cell populations within a 10–50-micron radius. Created in BioRender. Pomada Villalbi, A. (2025) https://BioRender.com/3hohunl B) Four mice were injected with sLP mCherry^+ve^ Med8a cells, while two control mice were injected with parental D425 lacking sLP mCherry. At endpoint, brain and spinal cord tissues from both experimental and control groups were harvested, followed by leptomeningeal dissociation with liberase. Macrophages from the control group were sorted using Adgre1 antibody, while mCherry- positive GFP-negative cells were sorted from experimental groups for subsequent single cell SMART-seq transcriptomic analysis. C) UMAP feature expression plot of SMART-seq-derived mCherry^+ve^ (TAM) and mCherry^−ve^ macrophages. The expression of *Ptprc*, *Adgre1*, and *Mrc1* are prevalent in most cells, but not *Tmem119*, indicative of a perivascular macrophage phenotype (Dimension = 7, Resolution = 0.1). D) UMAP feature expression plot showing mCherry^+ve^ (TAM) and mCherry^−ve^ macrophages as distinct heterogeneous clusters, suggesting distinct macrophage phenotypes (Dimension = 7, Resolution = 0.1). E) Differential gene analysis comparing mCherry^+ve^ and mCherry^−ve^ macrophages. Upregulated genes in tumor associated macrophages (TAM), particularly lipid transport genes, are highlighted in red font on the right side of the volcano plot. F) Gene ontology of transcriptional differences between mCherry^+ve^ macrophages versus mCherry^− ve^ controls, revealing a lipid-laden phenotypic in mCherry^+ve^ macrophages. G) Violin plots depicting *Abcg1* expression between mCherry^+ve^ macrophages versus mCherry^−ve^ macrophage controls (adjusted P < 0.0001). H) Immunofluorescence staining for Abcg1 and Adgre1 on spinal leptomeninges of D425 xenografted mice, visualized with 63X spinning disk confocal microscopy. Each scale bar denotes 20 µm. I) Nested whisker plot comparing the percentage of Abcg1^+ve^/Adgre1^+ve^ double positive cells from leptomeningeal metastases versus control leptomeninges (N=3 nude mice, 10 control leptomeningeal images, 5 micro-metastasis images, 10 macro-metastasis images, Unpaired t-test, **** P < 0.0001). J) Violin plots displaying Nr1h3 expression levels between mCherry^+ve^ macrophages versus mCherry^−ve^ macrophage controls (adjusted P < 0.0001). K) Immunofluorescence staining for Nr1h3 and Adgre1 of D425 patient-derived xenografted spinal meninges (micro-mets and macro-mets) versus control normal leptomeninges, captured by 63X spinning disk confocal microscopy. Each scale bar denotes 20 µm. L) Nested whisker plot illustrating the percentage of double positive Nr1h3^+ve^/Adgre1^+ve^ cells in leptomeningeal metastases versus normal control leptomeninges (N=3 nude mice, 5 control leptomeningeal images, 5 micro-metastasis images, 9 macro-metastasis images, Unpaired t-test, * P < 0.05, **** P < 0.0001).

The most abundant type of mCherry^+ve^ EGFP^−ve^ cell type identified from the leptomeninges were macrophages, which were transcriptionally distinct from mCherry^−ve^ macrophages that were presumably remote from the tumor cells (Figure 6C-E; feature plots of leptomeningeal resident border-associated macrophage markers showing low *CCR2* expression and enrichment of *CX3CR1, LYVE1, and MRC1* in Figure S7B). These mCherry^+ve^ macrophages showed high levels of expression of lipid pathway genes, particularly cholesterol, by both gene expression (Figure 6E; see Supplemental Table 4 for full gene list) and pathway analysis (Figure 6F). mCherry^+ve^ macrophages expressed higher levels of cholesterol efflux genes *Abcg1* (Figure 6G) and *Nr1h3* (*LXRa*) (Figure 6J) compared to mCherry^−ve^ macrophage (see feature plots in Figure S7C).

We then performed co-immunofluorescence histology analysis on the macrophage marker Adgre1 along with either Abcg1 or Nr1h3 to compare their co-expression in normal leptomeninges without metastatic tumor cells, or in the leptomeninges with micro- or macro-mets (Figure 6, H and K; panels of expanded ABCG1 staining on primary tumor in Figure S7D). Quantitative assays on the co-immunofluorescence histology images showed that the double positive ADGRE1+ve/ABCG1+ve and ADGRE1+ve/NR1H3+ve cells were heavily populated in the tumor-bearing leptomeninges, particularly in the presence of macro-mets (Figure 6, I and L; quantification and high-magnification images of *Math1- GFP/J2Q-SB11/T2Onc/Ptch*^+/−^ mice are shown in Figure S8A, B, *Abcg1* feature plots of Adgre1+ cells from Abeysundara et al., 2025 scRNA [17] in Figure S8C).

To determine whether our observed findings reflected a developmentally conserved signaling network between rhombic lip (RL) cell populations and their local microenvironment. We leveraged the publicly available human developmental dataset by Mannens *et al* [26]. Interactome analysis using CellChat and CCInx consistently uncovered a cholesterol signaling hub with microglia as significant mediators to various rhombic lip derived cell-types (ligand receptor pair predictions between microglia and cerebellar rhombic lip ventricular zone in Figure S8D; CellChat chord diagram depicting microglia as dominant cholesterol signaling senders to RL derived cell populations in Figure S9A). Collectively, the data are consistent with a model in which macrophages enriched in lipid, modulated by the metastatic microenvironment, produce and release high levels of cholesterol that are internalized by tumor cells, thereby promoting metastatic progression.

### Inhibiting lipid-laden macrophage recruitment decreases metastatic burden

The metabolic dependence of leptomeningeal medulloblastoma prompted us to examine whether tumor cells actively recruit lipid-rich macrophages into the metastatic niche. CXCL12 is a well-characterized macrophage chemoattractant [27]. Our SMART-seq data showed that macrometastases expressed higher levels of *CXCL12* compared to micrometastases, and peritumoral macrophages expressed higher levels of the CXCL12 receptor, ACKR3 (CXCR7), compared to control remote macrophages (Figure 7, A and B), consistent with directed migration along CXCL12 gradients [28]. In contrast, the expression level of *CXCR4*, the canonical alternative receptor for CXCL12, was not different between tumor-associated and control macrophages in our SMART-seq dataset. In addition, *CXCL11*, the alternative ligand for ACKR3, was not detected in the tumor cells. Collectively, these findings identify CXCL12 as the principal chemokine regulating macrophage recruitment in this system.

**Figure 7:**
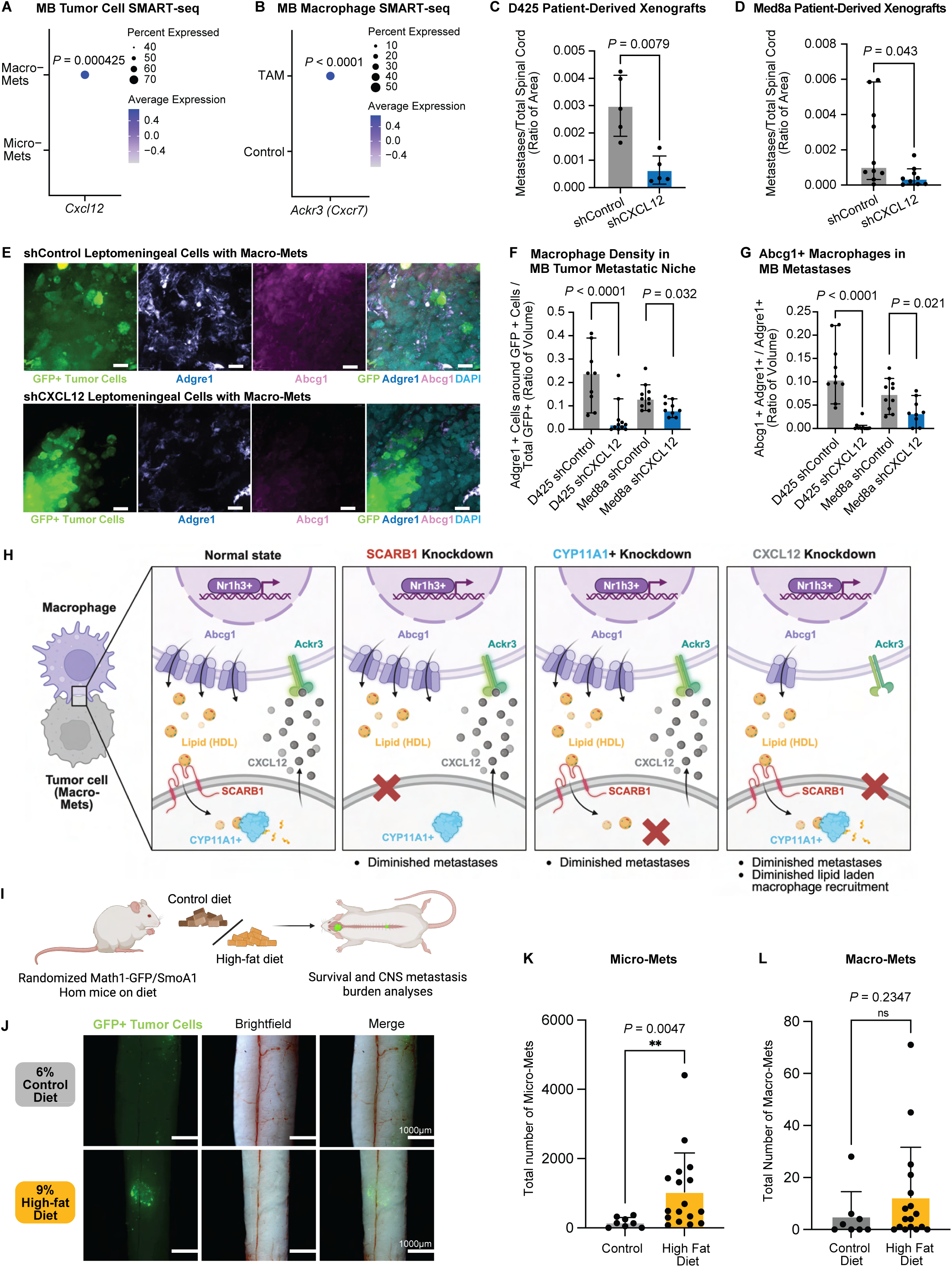
Lipids Delivered by Lipid-Laden Macrophages Are Necessary for Leptomeningeal Dissemination of Medulloblastoma. A) Dot plot from the Smart-seq dataset, showing elevated expression of *Cxcl12* in macrometastases as compared to micro metastases. B) Dot plot from the Smart-seq dataset, revealing increased expression of *Ackr3* in TAM tumor- associated macrophages compared to control macrophages. C) shCXCL12 D425 xenografted mice have a diminished metastatic burden at autopsy versus shCXCL12 D425 control xenografted mice (p = 0.0.0079). D) shCXCL12 MED8A xenografted mice have a diminished metastatic burden at autopsy versus shCXCL12 Med8a control xenografted mice (p = 0.043). E) Immunofluorescence staining for both Abcg1 and Adgre1 in spinal leptomeninges from nude mice xenografted with either shCXCL12 D425 or shControl D425 xenografts demonstrates decreased expression of both Abcg1 and Adgre1 after loss of Cxcl12. Visualized with 63X spinning disk confocal microscopy. Each scale bar denotes 20 µm. F) Adgre1 staining of microdissected leptomeningeal samples from nude mice xenografted with either shCXCL12 D425/Med8a or shControl D425/Med8a. A nested whisker plot compares the percentage of Adgre1 expression near GFP+ cells in shCXCL12 D425/Med8a versus shControl D425/Med8a samples demonstrates loss of Adgre1 expression after loss of CXCL12 (Unpaired t- test, * p < 0.05, **** p < 0.0001). G) Abcg1 and Adgre1 staining of microdissected leptomeningeal samples from nude mice xenografted with either shCXCL12 D425/Med8a or shControl D425/Med8a xenografts, with nested whisker plots comparing the percentage overlap of Abcg1 and Adgre1 (Unpaired t-test, * p < 0.05, **** p < 0.0001). H) Cartoons to illustrate possible models of therapeutic approaches to inhibit the pro-growth effect of macrophage delivered lipids on leptomeningeal metastasis growth. Created in BioRender. Pomada Villalbi, A. (2025) https://BioRender.com/ihvapuw I) Schematic diagram depicting an *in vivo* murine Phase 0 clinical trial to determine the causative role of lipids in the growth of leptomeningeal metastases. Created in BioRender. Pomada Villalbi, A. (2025) https://BioRender.com/a1yw74s J) Representative bright field-fluorescent stereoscope images of primary tumors and leptomeningeal metastases in SmoA1 Sleeping Beauty mice fed either a control or high-fat diet. Each scale bar denotes 1000 µm. K) Quantification of micro-metastases in SmoA1 Sleeping Beauty mice on control or high-fat diets (N=22 mice, Unpaired t-test, * p = 0.0047). L) Quantification of macro-metastases in SmoA1 Sleeping Beauty mice on control or high-fat diets (N=22 mice, Unpaired t-test, non-significant p = 0.2347).

To define the functional requirement of this axis, we knocked-down and performed *shCXCL12* in D425 and Med8a cells prior to xenografting (representative whole mount spine images and knockdown validation IF are in Figure S5E, F and Figure S9B). CXCL12 depletion markedly reduced leptomeningeal metastatic burden (Figure 7, C–D) and diminished macrophage infiltration (Figure 7E), including ABCG1 macrophages, within the peritumoral compartment (Figure 7, F-G; high-magnification images of the shCXCL12 D425 leptomeningeal environment in Figure S9C, D). These findings support a model in which CXCL12-mediated macrophage recruitment is essential for establishing a lipid-enriched microenvironment that enables the metabolic transition from dormant micro-mets to proliferative macro-mets (Figure 7H). Together, these data position CXCL12-dependent recruitment of lipid-laden macrophages as a necessary step in establishing a cholesterol-enriched niche that licenses the metabolic shift from dormancy to proliferative outgrowth in leptomeningeal metastasis.

### Cholesterol drives the growth of medulloblastoma metastases in vivo

To test if high systemic levels of cholesterol drive metastatic dissemination, we generated a colony of *Math1-GFP/Nestin-SB100/T2Onc2/SmoA1*^tg/tg^ mice fed either standard diet (6% fat) or high fat diet (HFD, 9% fat) immediately after weaning (Figure 7I). Collection and imaging analysis at humane endpoint for these animals showed that tumor-bearing HFD mice had greater metastatic burden in comparison to control tumor-bearing animals (Figure 7J; percent nutrient composition of control vs HFD is provided in Figure S9F). Tumor-bearing HFD mice had more micrometastases and a trend to increased burden of macrometastases (Figure 7, K and L). This *in vivo* murine preclinical trial supports a model in which high systemic levels of lipids promoted the metastatic cascade in medulloblastoma.

## DISCUSSION

Medulloblastoma remains the most common lethal pediatric brain tumor. Comprehensive molecular analyses of primary medulloblastoma have informed tumor classification, but the leptomeningeal metastases, rather than the primary tumor are usually the cause of tumor-related patient mortality. Metastatic recurrence of medulloblastoma is nearly universally fatal. No specific or targeted therapies for metastatic medulloblastoma are currently available, either upfront at diagnosis, or at recurrence. The biological divergence between tumor cells within primary medulloblastoma and from medulloblastoma metastases suggests that therapies developed based solely on study of the primary tumor are unlikely to be effective in the metastatic compartment. A reduction in the morbidity and mortality for medulloblastoma requires additional understanding and modeling of the metastatic compartment.

Medulloblastoma patients without evidence of leptomeningeal metastases at the time of presentation or after therapy may still recur with metastatic disease. The biological basis for metastatic recurrence after a clean MRI is not currently known, but we suggest that metastatic recurrences could arise from leptomeningeal micrometastases, which would be invisible on current clinical MRI imaging. Indeed, micrometastases are not mitotically active, so they are likely relatively resistant to radiation therapy as compared to the primary tumor or the cycling macrometastases.

Here, we find that medulloblastoma metastases recruit lipid-laden macrophages by secreting CXCL12 into the leptomeningeal tumor microenvironment to support tumor growth. Previous reports have shown that excessive SHH signaling in medulloblastoma promotes lipogenesis, [29] and can promote therapeutic resistance [30]. Statins have subsequently shown preclinical activity against SHH medulloblastoma [31, 32]. Group 3 and 4 medulloblastomas also show distinct lipid profiles, albeit in primary tumors [33].

Concordant with our findings, mass spectrometry identified lipid accumulation in medulloblastoma metastases [34]. Based on this background and our data, multiple therapeutic interventions could be considered based on targeting the axis between tumor cells and supporting macrophage: agents to minimize systemic lipid levels, to disrupt recruitment of lipid-laden macrophages, to block disgorgement of cholesterol into the tumor microenvironment, or to inhibit uptake of lipids by metastatic medulloblastoma cells could provide benefit to patients to either prevent or treat leptomeningeal metastases. As there are currently no targeted or specific therapies for metastatic medulloblastoma, the therapeutic possibility for novel agents is very wide. Our data support a model in which high levels of lipid in the tumor microenvironment promote cell cycle progression. An alternative translational approach suggested by this model is to provide children with a high fat diet while they are receiving radiation and/or chemotherapy, to drive previously quiescent stem cells into the cell cycle where they are vulnerable to radiation induced cell death. Further translational experiments in model systems are likely necessary to determine if strategies to diminish or elevate lipid levels are attractive strategies to test in the clinic for children with medulloblastoma.

## LIMITATIONS OF THE STUDY

A key limitation of this study is the lack of an established transgenic mouse model that faithfully recapitulates Group 3 medulloblastoma. For our *in vivo* immunofluorescence validation, we relied on xenograft models in nude mice. Although these mice support the macrophage focused questions central to our work, their limited and naïve T-cell compartment may influence tumor–immune interactions.

## ACKNOWLEDGEMENTS

V.F. is supported by The Frank Fletcher Memorial Fund, Ontario Graduate Scholarship, CIHR Frederick Banting and Charles Best Canada Graduate Scholarship (Master’s award), and the University of Toronto Fellowship (Laboratory Medicine and Pathobiology). M.L. is supported by the University of Toronto Fellowship and SickKids Research Training Competition Awards. M.D.T. is a CPRIT Scholar in Cancer Research (CPRIT - RR220051). M.D.T. is the Cyvia and Melvyn Wolff Chair of Pediatric Neuro-Oncology, Texas Children’s Cancer and Hematology Center. M.D.T. is supported by the NIH (R01NS106155, R01CA159859 and R01CA255369), NCI (2P50CA127001-16), The Pediatric Brain Tumour Foundation, The Terry Fox Research Institute, The Canadian Institutes of Health Research, The Cure Search Foundation, Matthew Larson Foundation (IronMatt), Meagan’s Walk, SWIFTY Foundation, The Victory Foundation, The Brain Tumour Charity, Genome Canada, Genome BC, Genome Quebec, the Ontario Research Fund, Worldwide Cancer Research, V-Foundation for Cancer Research, and the Ontario Institute for Cancer Research through funding provided by the Government of Ontario. M.D.T. was also supported by a Canadian Cancer Society Research Institute Impact grant, a Cancer Research UK Brain Tumour Award, and by a Stand Up To Cancer (SU2C) St. Baldrick’s Pediatric Dream Team Translational Research Grant (SU2C-AACR-DT1113) and SU2C Canada Cancer Stem Cell Dream Team Research Funding (SU2C-AACR-DT-19-15) provided by the Government of Canada through Genome Canada and the Canadian Institutes of Health Research, with supplementary support from the Ontario Institute for Cancer Research through funding provided by the Government of Ontario. Stand Up to Cancer is a program of the Entertainment Industry Foundation administered by the American Association for Cancer Research. V.R. is supported by a Canada Research Chair in Pediatric Neuro-Oncology and operating funds from the Canadian Institutes for Health Research (PJT- 488102 and PJT- 400627), Rally Foundation for Childhood Cancer Research and Kids Join the Fight, The Medulloblastoma Initiative, Canadian Cancer Society Research Institute, the C.R. Younger Foundation and Brain Canada through the Canada Brain Research Fund with the financial support of Health Canada and Alvin Segal Family Foundation Future Leader in Canadian Brain Research. Y.H. and R.A. are supported by PSC-CUNY Faculty Research Award. K. V. and Y. X. C were supported by the Sloan Foundation CSURP program. We thank Ariadna Pomada Villalbi for graphic design and figure editing. We thank The Centre for Phenogenomics’ and the Animal Research Center’s veterinary staff for their expertise and animal care. We thank the SickKids-UHN Flow Cytometry facilities for their expertise. We acknowledge K. Lau and P. Paroutis from the SickKids Imaging Facility for instrument support and expertise. We also acknowledge The Centre for Phenogenomics’ veterinary staff and animal technicians for colony management and monitoring. We thank Susan E. Archer for technical writing expertise.

## CONTRIBUTIONS

Conceptualization, V.F., M.L., N.A., S.S., S. A., J.N.R., V.R., M.D.T., and X.W.; Data Curation, A.W.E.; Formal Analysis, V.F., M.L., N.A., A.W.E., L. H., P.B., J.M., P.S.; Methodology, V.F., M.L., N.A., S.S., S.A., J.N.R., V.R., M.D.T., and X.W.; Investigation, V.F. M.L. N.A., R.V.O., P.B., J.M., B.L., O. Sirbu, N.M., S.A.K.,; Resources: V.F., M.L., A.W.E., N.A., L.H., R.V.O., P.B., J.M., B.L., P.S., O. Sirbu, N.M., J.Z., C.R., R.S., N.H., H.W., L.Q., T.D., J.P., E.M., S.A.K., A.K.K., H.V., L.Z., K.V., Y.X.C., O.O., V.V., F.T., D.K., V.D., L.X., M.H., J.J.F., D.P., A.Y., K.K., A.R., E.W., W.O., T.J., Q.Y.; Project Administration, C.D., M.D.T., and X.W.; Funding Acquisition, C.D., M.D.T., and X.W.; Supervision, X.H., O.A., R.W.R., S.E.E., D.L., R.J.D., H.Y., R.A., C.D., L.S., D.W.E., G.B., C.-H.L., O.S., D.S., J.R., S.S., S.A., J.N.R., V.R., M.D.T., X.W.; Validation, V.F., M.L.; Visualization, V. F. and M.L.; Writing – Original Draft, V.F., M.L., N.A., C.D., S.S., S.A., J.N.R., V.R., M.D.T., and X.W.; Writing – Reviewing & Editing, V.F., M.L., C.D., N.A., S.S., S.A., J.N.R., V.R., M.D.T., and X.W.; S.S., S.A., J.N.R., V.R., M.D.T., and X.W. are joint senior authors and project co-leaders.

## SUPPLEMENTAL FIGURE LEGENDS

**Figure S1**

**A**) Representative bright field-fluorescent stereoscope images of leptomeningeal metastases from D425 GFP-Luc xenografts, highlighting micro-mets. Each scale bar denotes 1000 µm.

**B**) Confocal images of leptomeningeal metastases from D425 GFP-Luc xenografts, highlighting micro-mets. Each scale bar denotes 20 µm.

**C**) Representative bright field-fluorescent stereoscope images of leptomeningeal metastases from D425 GFP-Luc xenografts, highlighting macro-mets. Each scale bar denotes 1000 µm,

**D**) Confocal images of leptomeningeal metastases from D425 GFP-Luc xenografts, highlighting macro-mets. Each scale bar denotes 20 µm.

**Figure S2**

**A**) Bar plots illustrating *in vivo* glucose tracing results, highlighting ^13^C glucose incorporation into metabolites and amino acids from the citric acid cycle in primary tumors, micro-mets, and macro-mets (N=2 mice).

**B**) Bar plots of genes representing enriched pathways in fatty acid/lipid metabolism and phospholipid biosynthetic processes, derived from differentially upregulated genes in macro-mets compared to micro-mets, with log2FC and adjusted p-values.

**Figure S3**

**A**) Differential expression of metabolites in macro- and micro-mets. Of the 774 metabolites quantified in both positive and negative ion modes, 221 metabolites with a variance >0 were retained for downstream analysis. The volcano plot illustrates the relationship between log 2 (fold change) and −log 10 (p-value), comparing macro- and micro-mets (n=5 per group) using a Wilcoxon rank-sum test. For comprehensive visualization, metabolites with non-finite values were positioned just beyond the range of extreme finite values.

**B**) DAS plot from untargeted shotgun lipidomics, illustrating the number of significant lipids within each lipid category. A value of 1 indicates all lipids within that family are significantly increased/decreased.

**C**) Schematic diagram of the protocol for MALDI-IMS of whole-mounted leptomeninges.

**D**) MALDI imaging detected various metabolites differentially expressed between micro- and macro-mets in Ptc Het Sleeping Beauty mouse leptomeninges. Small macro-mets are outlined in red, micro-mets in black. MALDI imaging revealed elevated levels of 20:0 lysophosphatidylethanolamine (20:0 LysoPE ) and hexadecenoic acid (palmitic acid) in macro-mets compared to micro-mets, and elevated levels of o-linoleoylcarnitine and 3-hydroxy-9- hexadecenoylcarnitine in micro-mets compared to macro-mets.

**Figure S4**

**A**) Filipin staining was performed on peeled meningeal samples from D425-GFPluc xenografts. Images were acquired using 63X spinning disk confocal microscopy. Stronger accumulation in macro-mets compared to micro-mets and normal leptomeningeal cells was observed. Each scale bar denotes 20 µm.

**B**) Violin plots showing differential expression of *Ldlr* and *Lrp1* between micro- and macro-mets from Shh GEMM models, based on SMARTSEQ data.

**C**) SCARB1 staining were performed on brain slices from two-month-old D425 patient-derived xenograft samples. Stronger SCARB1 accumulation were observed in tumor cells compared to control neurons. Images were acquired using 20X spinning disk confocal microscopy. Each scale bar denotes 70 µm.

**D**) Western blot analysis of D425 cells with shControl or shSCARB1 showed significantly diminished *SCARB1* protein levels in shSCARB1 compared to shControl.

**Figure S5**

**A**) Representative bright field & fluorescent (GFP) stereoscope images of whole spinal leptomeningeal metastases from D425 shControl GFP-Luc xenografts. Each scale bar denotes 1000 µm.

**B**) Representative bright field & fluorescent (GFP) stereoscope images of whole spinal leptomeningeal metastases from D425 shSCARB1 GFP-Luc xenografts. Each scale bar denotes 1000 µm.

**C**) Representative bright field & fluorescent (GFP) stereoscope images of whole spinal leptomeningeal metastases from D458 shControl GFP-Luc xenografts. Each scale bar denotes 1000 µm.

**D**) Representative bright field & fluorescent (GFP) stereoscope images of whole spinal leptomeningeal metastases from D458 shCYP11A1 GFP-Luc xenografts. Each scale bar denotes 1000 µm.

**E**) Representative bright field & fluorescent (GFP) stereoscope images whole spinal leptomeningeal metastases from D425 shControl GFP-Luc xenografts. Each scale bar denotes 1000 µm.

**D**) Representative bright field & fluorescent (GFP) stereoscope images of whole spinal leptomeningeal metastases from D425 shCXCL12 GFP-Luc xenografts. Each scale bar denotes 1000 µm.

**Figure S6**

**A**) Volcano plot depicting results of bulk RNA-sequencing–based differential gene expression analysis comparing shControl and shSCARB1 D425 cells.

**B**) Quantification of CYP11A1 accumulation in D458 primary tumors comparing shControl and shCYP11A1 groups (n = 10 images per group). Statistical significance was determined by unpaired two-tailed t-test (**** P < 0.0001).

**C**) 10X single-cell RNA-seq dataset from Abeysundara et al. 2025^17^. Feature plots show CYP11A1 is predominantly expressed in tumor cells from the *Math1-GFP/active SB/Ptch+/− (PtcSB)* tissues, with minimal presence in surrounding microenvironmental cells and from the *Math1-GFP/active SB/Ptch+/+ (PtcCon*) tissues.

**Figure S7**

**A**) Immunofluorescence was performed on spinal slices of mice xenografted with patient-derived medulloblastoma cells transduced with SLP*-*mCherry virus. Tumor cells expressed both GFP and mCherry signals, whereas tumor microenvironment niche cells expressed only the mCherry signal. Each scale bar denotes 30 µm.

**B**) Feature plots of leptomeningeal resident border-associated macrophages markers showing low CCR2 expression and enrichment of CX3CR1, LYVE1, and MRC1 in both mCherry+ and mCherry- cells.

**C**) Feature plots of the top two lipid storage functions were generated, comparing mCherry+ and control macrophages. *Abcg1* and *Nr1h3* expression were the focus of this study, as both were consistently upregulated across all mCherry+ batches.

**D**) Abcg1 staining were performed on brain slices from two-month-old D425 patient-derived xenograft samples. Stronger Abcg1 accumulation were observed in control neurons compared to tumor cells. Images were acquired using 20X spinning disk confocal microscopy. Each scale bar denotes 70 µm.

**Figure S8**

**A**) Immunofluorescence staining for Abcg1 and Adgre1 on spinal meninges from *PtcSB* and *PtcCon* samples, visualized with 63X spinning disk confocal microscopy. Each scale bar denotes 20 µm.

**B**) Nested whisker plot comparing the percentage overlap of Abcg1 and Adgre1 in *PtcSB* versus *PtcCon* leptomeningeal areas (N=3, 9 control leptomeningeal images, 10 macro-metastasis images, Unpaired t-test, ** P < 0.01, **** P < 0.0001).

**C**) 10X single-cell RNA-seq dataset from Abeysundara et al. 2025^17^. Feature plots show Abcg1 expression is upregulated in immune cells of *PtcSB* compared to immune cells of *PtcCon*, **** P < 0.0001.

**D**) Semi-supervised Ligand receptor pair predictions between microglia and cerebellar rhombic lip ventricular zone (RL-VZ). Cholesterol related ligands are some of the most significant predicted interactions between these two populations. Node colours represent mean normalized gene expression for each cell type.

**Figure S9**

**A**) CellChat Chord diagram depicting microglia as dominant cholesterol signaling senders to RL derived cell populations in developing cerebellum.

**B**) Quantification of CXCL12 expression in D425 primary tumors comparing shControl and shCXCL12 groups (n = 9 images per group). Statistical significance was determined by unpaired two-tailed t-test (** P < 0.01).

**C**) *Abcg1* and *Adgre1* staining were performed on spinal meninges from two-month-old nude shControl D425 xenografts. Images were obtained using 63X spinning disk confocal microscopy. Each scale bar denotes 20 µm

**D**) *Abcg1* and *Adgre1* staining were performed on spinal meninges from two-month-old nude shCXCL12 D425 xenografts. Images were obtained using 63X spinning disk confocal microscopy. Each scale bar denotes 20 µm.

**E**) Table displaying the percent nutrient compositions of control and high-fat diet.

## STAR METHODS

### Key Resources Table

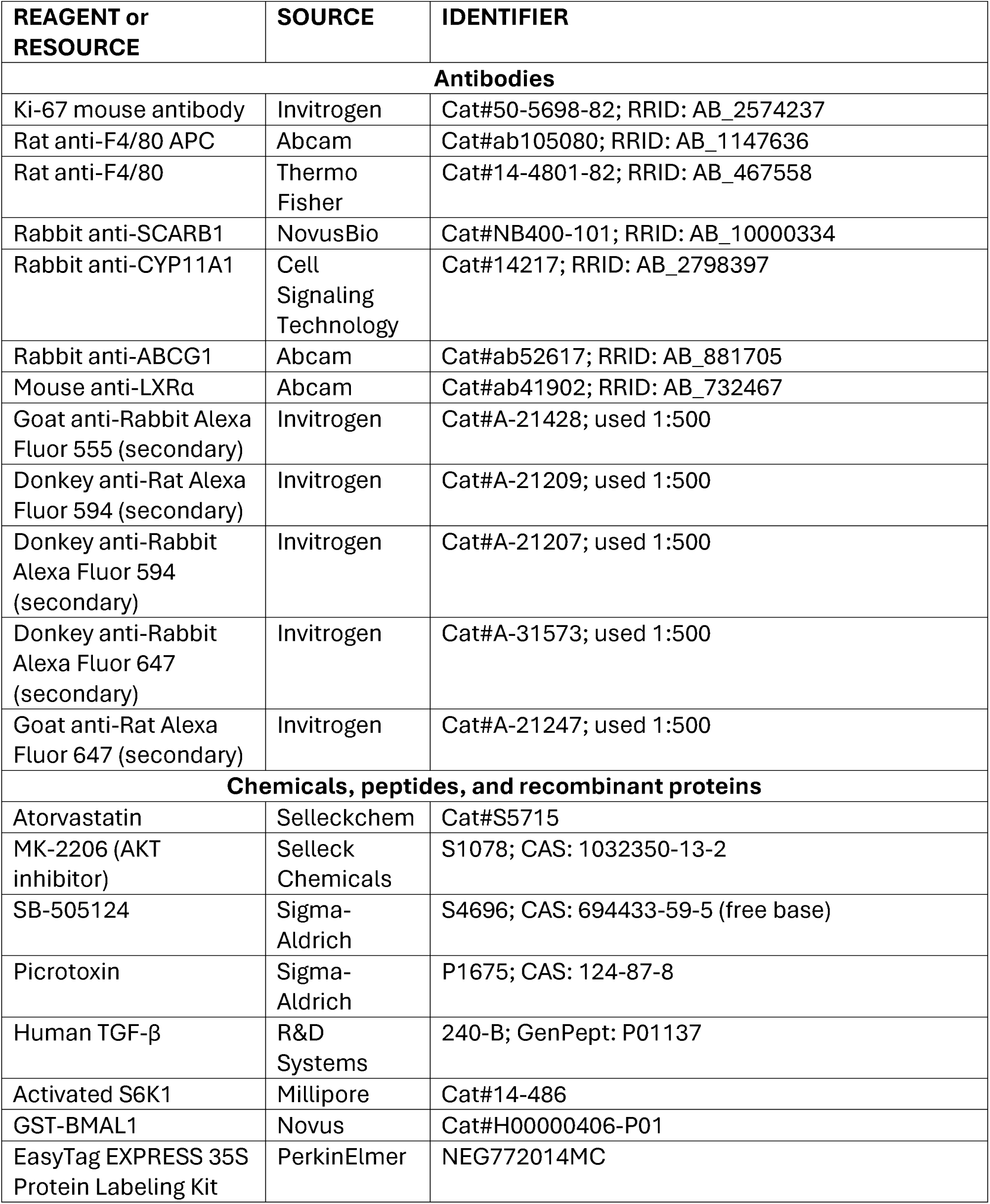
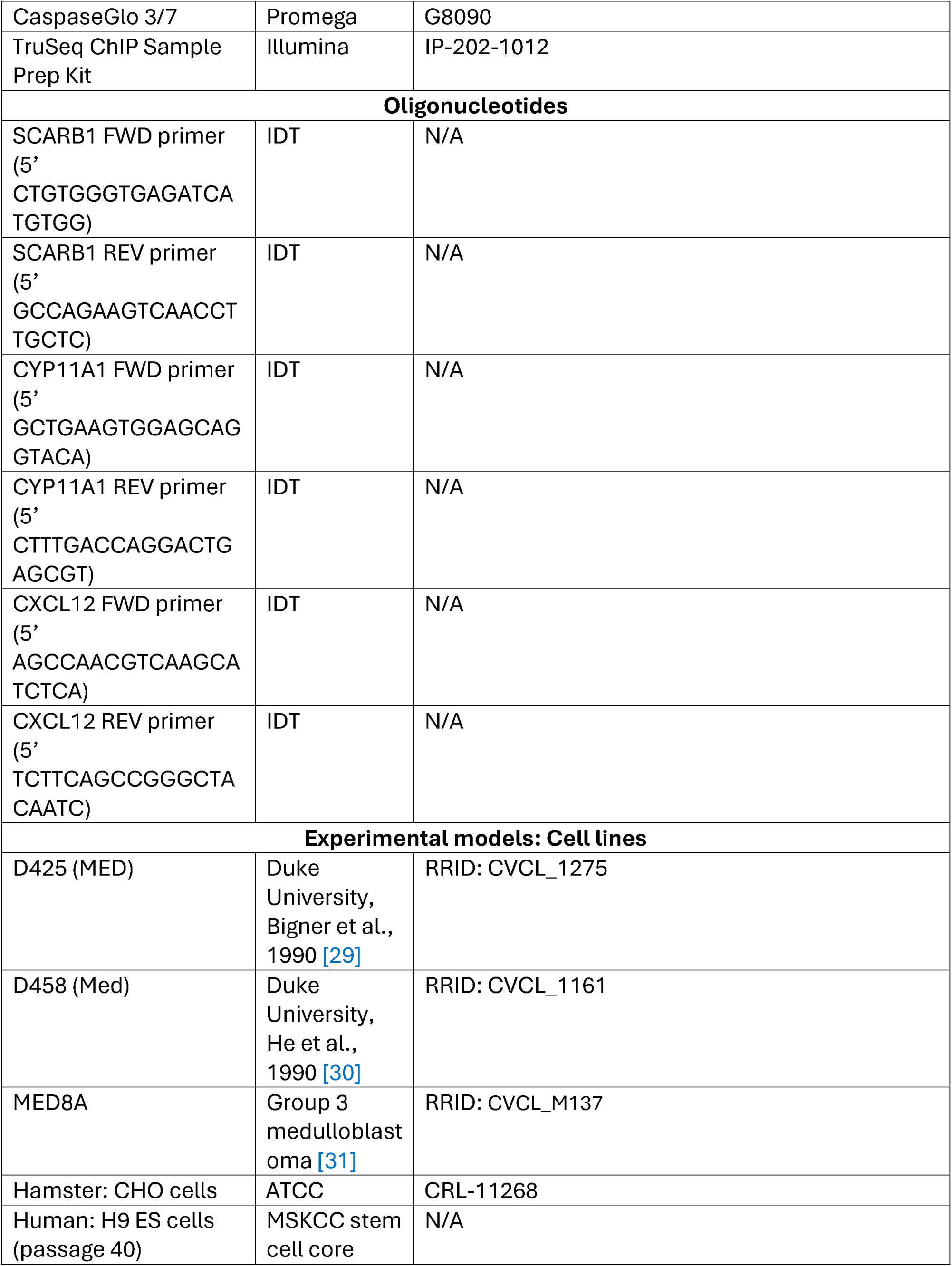
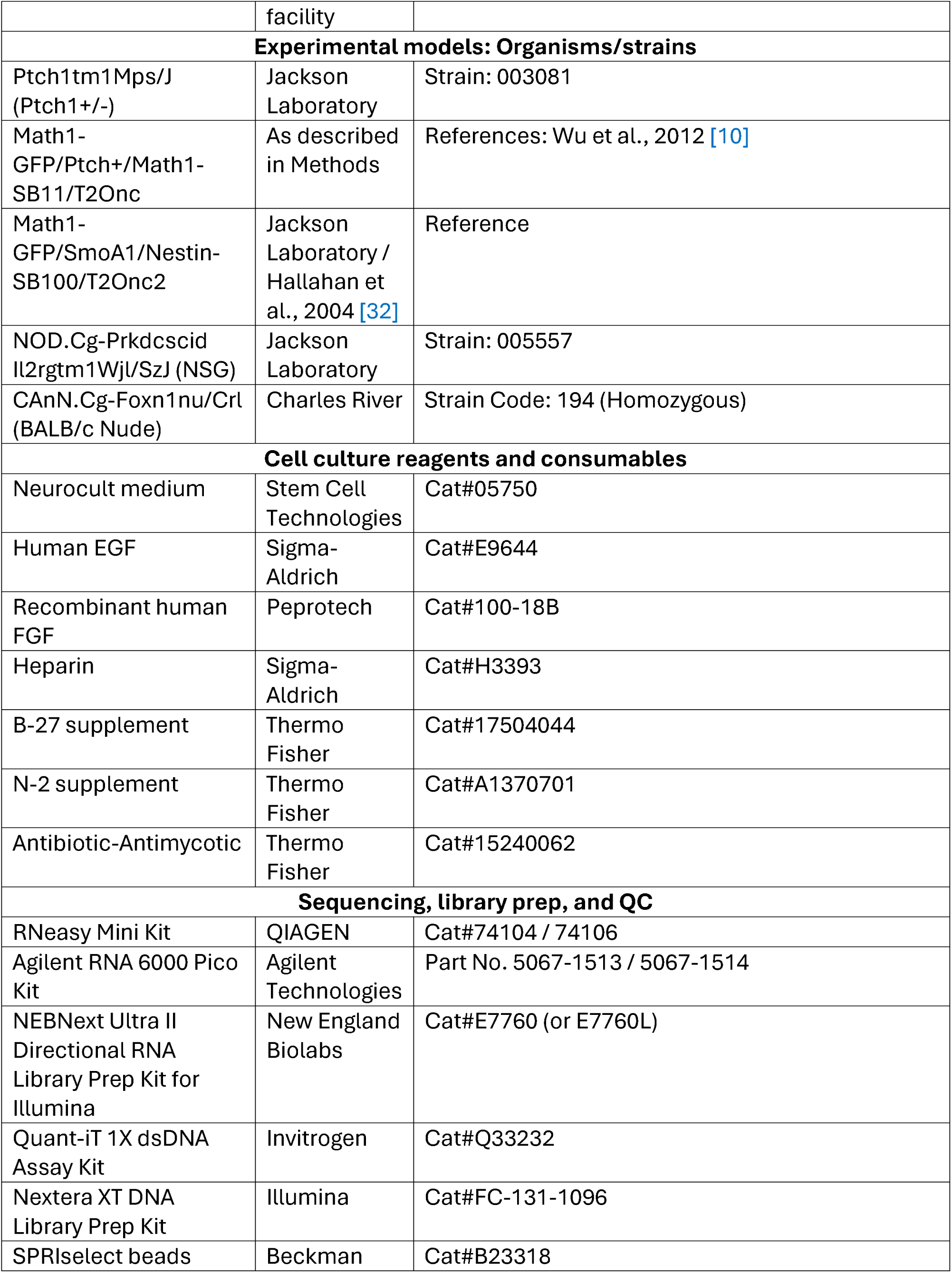
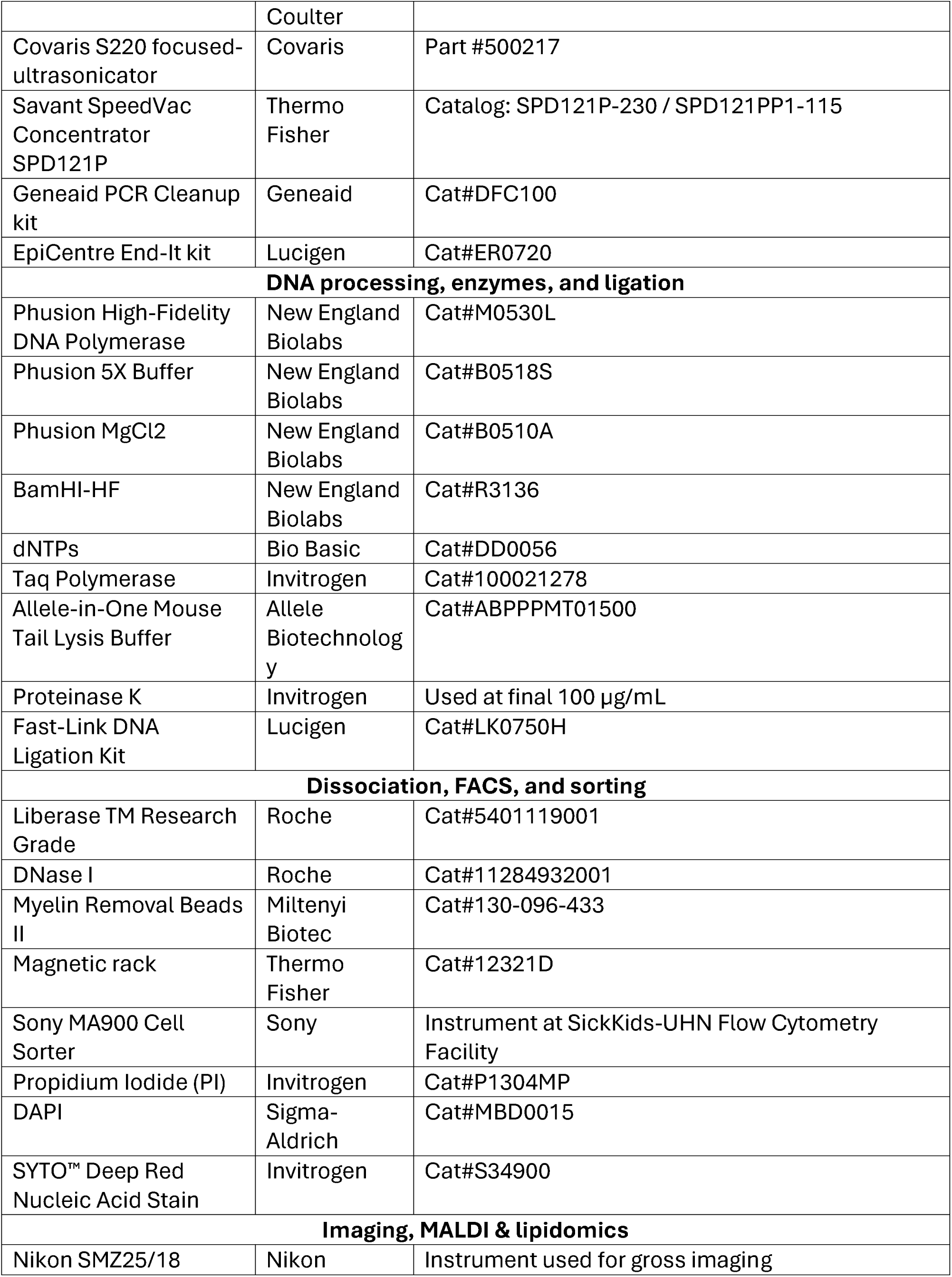
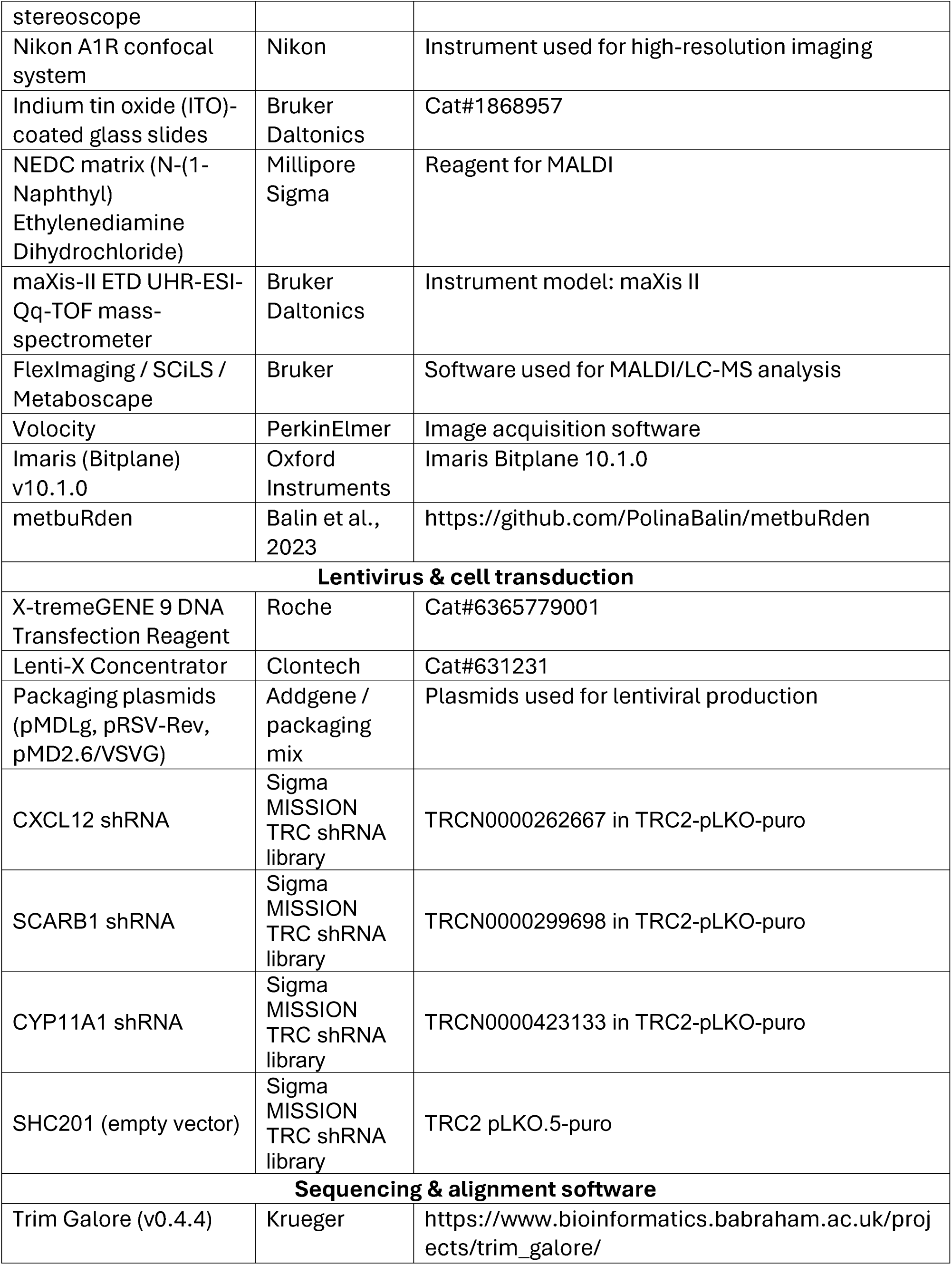
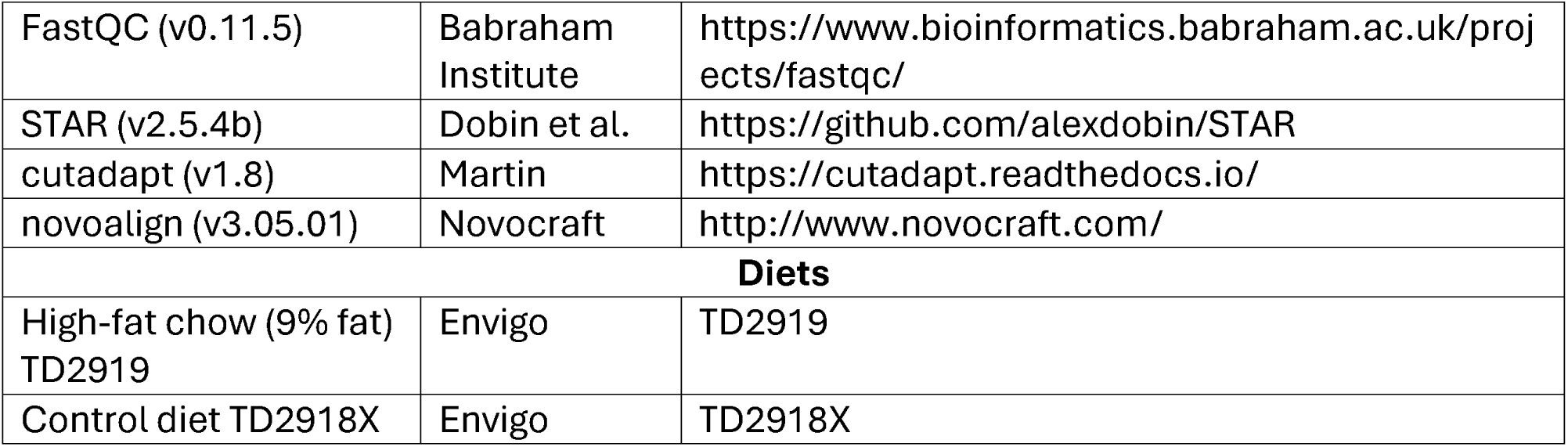

### Materials and Methods

#### Mice

Ptch+/−/Math1-SB11/T2Onc were generated from crossing *Ptch1* heterozygous (Ptch1 tm1Mps /J, #003081 Jackson Laboratories) with *T2Onc* (TgTn(sb-T2/Onc)76Dla) and Math1- SB11 mice [10]. These mice were crossed with Math1-GFP mice to generate quadruple transgenic mice. *Math1-GFP/Ptch+/Math1-SB11/T2Onc* mice were previously generated as described [16][10]. *Nestin-SB100/T2Onc2* mice were crossed with *Math1-GFP/SmoA1* to generate the quadruple transgenic mouse line *Math1-GFP/SmoA1/Nestin- SB100/T2Onc2* with high MB penetrance and metastatic burden. NOD scid gamma (NSG) mice were purchased from Jackson Laboratories and bred at the Toronto Centre for Phenogenomics. Both females and males were used in mouse experiments. BALB/c Nude mice (CAnN.Cg-Foxn1nu/Crl, Strain Code: 19*4* Homozygous) were purchased from Charles River. All animals bred at the Toronto Centre for Phenogenomics (Toronto, ON) were housed according to the guidelines from the Canadian Animal Care Committee. Animals bred at Charles River follows were housed according to the guidelines by the Canadian Council on Animal Care (CCAC) and accredited by Association for Assessment and Accreditation of Laboratory Animal Care (AAALAC).

#### Genotyping

Samples of ear notches from mice were kept frozen at -20°C until ready for lysing and genotyping. Samples were lysed in 25 µL of Allele-in-One Mouse Tail Lysis Buffer (Allele Biotechnology, #ABPPPMT01500) overnight at 56°C. Enzymatic activity was inactivated within 10 min, incubation at 95°C and samples were centrifuged briefly. Polymerase chain reactions (PCRs) were set up with the following master mix (2.5 µL 10X PCR Buffer (Invitrogen, #52724), 1 µL 50 mM MgCl2 (Invitrogen, #52723), 0.5 µL 10mM dNTP (Bio Basic, DD0056), 0.5 µL F primer, 0.5 µL R primer, 2 µL loading dye, 0.2 µL Taq polymerase (Invitrogen, #100021278, 15.8 µL ddH2O (Wisent, #809-115-CL) and 2 µL DNA), primers, and protocols according to the queried genes.

#### Cell Lines

D425 (6-year-old male medulloblastoma tumor; *RRID: CVCL_1275*), D458 (7-year-old female medulloblastoma tumor; RRID: CVCL_1161), and MED8A (Group 3 medulloblastoma tumor; RRID: *CVCL_M137*) were cultured in Neurocult medium (Stem Cell Technologies, #05750) supplemented with 20 ng/mL human epidermal growth factor (EGF) (Sigma-Aldrich, #E9644), 10 ng/mL recombinant human fibroblast growth factor (FGF) (Peprotech, #100-18B), heparin (Sigma-Aldrich, #H3393), B-27 (Thermo Fisher, #17504044), N-2 (Thermo Fisher, #A1370701), and antibiotic-antimycotic (Thermo Fisher, #15240062). Cell lines were authenticated by short tandem repeat (STR) profiling (Geneprint, Promega) and periodically tested for mycoplasma contamination.

#### Intracranial Injections of MB Tumor Cells

Medulloblastoma cell lines were xenografted by stereotactic injection into 6-8 weeks old NSG or Nude males and female mice using the following stereotactic coordinates for cerebellar injections: 2 mm posterior to lambda, 1 mm lateral and 2 mm deep. 6-8 weeks old NSG or Nude mice were also injected with tumor cells into the lateral ventricles to test for metastatic growth on the spinal leptomeninges using the following stereotactic coordinates: 3 mm posterior to bregma, 1.5 mm lateral and 2 mm deep. Tumors were allowed to develop until a humane endpoint as determined by the animal facility based on symptoms such as weight loss, tilted gait, domed head, and ataxia.

#### Imaging with the Nikon SMZ25 System

Brains and spinal cords are kept on ice until imaging analysis with the Nikon SMZ25/18 stereoscope system. Tissues are imaged in PBS to prevent drying. Both dorsal and ventral surfaces of the brains/spinal cords and the microdissected peripheral nerves were imaged. Brightfield images are taken with an exposure time of 200 msec. For transgenic animals with brighter GFP fluorescent signals, the exposure time of the primary tumor is 1 second and the spinal cord metastases is 5 seconds. For NSG animals injected with MB cell lines, the GFP or tdTomato fluorescent signal is dim. The exposure time of the primary tumor is 1 second and the spinal cord metastases is 5 seconds for optimal detection.

#### H&E Staining of Primary Tumor and Metastasis

The brain and spinal cord of animals were collected as described above. The brain with the primary tumor was placed into 10% formalin for a minimum of 24 hr. and then washed with PBS prior to submission to the pathology lab for H&E staining at the University Health Network. The spinal cord was subjected to imaging on the Nikon SMZ25 system as described and areas with micro- and macro-metastases were identified and isolated for fixation with 10% formalin. Again, fixation was 24hr. at minimum and samples were then washed with PBS and submitted to pathology for H&E staining.

#### Volumetric Boundary between Micro- and Macro-Metastasis

Different distributions (Gaussian, gamma, and Weibull) were fitted to the volumetric measurements by numerical optimization of the maximum likelihood. The fit of each distribution to the data were determined using quantile-quantile plots, and the Gaussian distribution was determined to have adequate fit to the volumetric measurements on a log scale. Subsequently, the fit of one Gaussian or a mixture of two Gaussian distributions to the data was assessed by the log likelihood ratio test, and the two-Gaussian mixture model was selected. The cut-point was set at the volumetric value for which the assignment probabilities for both mixture components are equal.

#### Statistical Tests for Bimodality of MB Metastasis

The assessment of bimodality for MB metastasis was carried out using volumetric data of micro- and macro-metastases that were collected from Z-stack imaging of the leptomeninges. Using R Studio, an R script using the Modes package was used to test the characteristics of a bimodal distribution which included the bimodality amplitude, the bimodality coefficient, and the bimodality ratio. The volumetric data was also analyzed for kurtosis and skewness. Finally, a Shapiro-Wilk Test to randomly sample 5000 samples from the volumetric data was completed to test for a normal distribution.

#### Shear-SPLINK Experiment

Spinal meningeal regions lacking grossly visible macrometastases were dissected, and areas containing GFP-positive micrometastatic foci were identified by fluorescence microscopy, excised as tissue fragments, and enzymatically dissociated for downstream processing and fluorescence-activated cell sorting. Individual micrometastatic lesions were not manually isolated. Micro- and macro-metastatic cell were then collected into Eppendorf tubes containing 1 mL PBS. The samples were centrifuged to form a pellet and lysed with Proteinase K. Lysis was completed by resuspending the samples in 100 uL of lysis buffer (10mM Tris-Cl and 0.1M EDTA with pH 8.0, 0.5% SDS) and incubating at 95°C for 10 min. Then, 2.5µL Proteinase K (final concentration 100µg/ml) was added and the samples were incubated at 56°C overnight. The Proteinase K was inactivated by heat at 95°C for 10 min.

Prior to shearing, the samples were lyophilized with a Savant SpeedVac Concentrator (Thermo Fisher, #SPD121P). The condenser and concentrator were turned on and set to 45°C heat, 1.5 hr. heat time, 1.5 hr. run time, and 0.1 vac setting. The vacuum pump was then turned on and the samples were loaded. After 1.5 hr. run time, all dehydrated samples were resuspended in TE buffer for downstream processing.

Due to the small tissue input from micro- and macro-metastases, all of the extracted DNA was submitted for mechanical shearing. Lyophilized samples were rehydrated with 50 µL of ddH2O and transferred to Covaris snap cap tubes for shearing to the desired length of 300 bp using the Covaris S220 sonicator at The Center for Applied Genomics (TCAG). Sheared samples were stored at -20°C.

Sheared DNA required end repair via the EpiCentre End-It kit (Lucigen, #ER0720). In PCR 8- strip tubes, 20 µL of sheared DNA was combined with 0.5 µL ddH2O, 3 µL End-It 10X Buffer, 3 µL 2.5 mM dNTP Mix, 3 µL 10mM ATP, and 0.5 µL End-It Enzyme Mix, then incubated at room temperature for 45 min prior to transfer to a thermocycler for 10 min. at 70°C. The samples were cooled before proceeding to ligation steps.

The adaptors must be prepared for ligation. For each sample reaction, 0.82 µL of 100 µM Linker+ (5’ GTATATACGACTCACTATAGGGCTCCGCTTAAGGGAC) and 0.82 µL of 100 µM Linker– (5’ Phos-GTCCCTTAAGCGGAG-C3 spacer) primers were combined and placed in the thermocycler to boil at 95°C for 5 min. and slowly cooled to room temperature. After the adaptors were prepared, ligation was completed with the EpiCentre Fast-Link DNA Ligation kit (Lucigen, #LK0750H) in which 1.64 µL of adaptors was added with 1.75 µL 10 mM ATP, 0.5 µL Fast-Link 10X Ligation Buffer, and 1.11 µL Fast-Link Ligase to each sample. Samples were placed in a thermocycler on the following program: 25°C for 15 min., 70°C for 15 min. and then cooled to 4°C before proceeding to digestion.

Shear-SPLINK samples were digested with 7.5 µL of molecular grade ddH2O (Wisent, #809-115-CL), 5 µL 10X BSA (Sigma-Aldrich, #A9647-100G), 1.5 µL NEB Buffer 4 (New England Biolabs, #B7004S), and 1 µL BamHI-HF (New England Biolabs, #R3136) overnight at 37°C.

Two primary PCRs were set up per sample, one for the IRL forward primer (5’ AAATTTGTGGAGTAGTTGAAAAACGA) and another for the IRR forward primer (5’ GGATTAAATGTCAGGAATTGTGAAAA). In each reaction, the following were combined: 17.25 µL digested DNA, 5 µL 5X Phusion Buffer (New England Biolabs, #B0518S), 0.75 µL 10 mM MgCl2 (New England Biolabs, #B0510A), 0.5 µL 10 mM dNTPs (Bio Basic, #DD0056), 0.5 µL 10 µM IRL or IRR primer, 0.5 µL Linker A-1 (5’ GTAATACGACTCACTATAGGGC) primer, and 0.5 µL Phusion Taq (New England Biolabs, #M0530L). The PCR protocol on the thermocycler was set up to 98°C for 30 sec. and then 25 cycles of 98°C for 20 sec., 55°C for 30 sec., and 72°C for 1 min., then 72°C for 2 min. and hold at 4°C.

To set up the secondary PCR reactions, diluted 3 µL of the primary PCR reactions in 147 µL ddH2O, and froze the rest at -20°C. Then vortexed the dilution to properly mix and incubated at room temperature for 30 min. A secondary PCR reaction was prepared for each primary PCR using 2 µL of 1:50 diluted primary PCR product, 1 µL 2.5 µM primer 1 / transposon barcode, 16.25 µL ddH2O (Wisent, #809-115-CL), 5 µL 5X Buffer (New England Biolabs, #B0518S), 0.5 µL 10mM dNTPs, 0.125 µL 10 µM primer 2 / Linker A2 (5’ CAAGCAGAAGACGGCATACGAGATCGGTCTCGGCATTCCTGCTGAACCGCTC

TTCCGATCTTAGGGCTCCGCTTAAGGGAC), and 0.5 µL Phusion Taq polymerase. The thermocycler conditions for the secondary PCR reactions were 98°C for 3 min., 10 cycles of 95°C for 30 sec., 49°C for 30 sec., and 72°C for 2 min., then 25 cycles of 95°C for 30 sec., 53.3°C for 1 min., and 72°C for 2 min., followed by 72°C for 10 min. and hold at 4°C. Once completed, 5 µL of each secondary PCR reaction was analyzed on a 1.5% agarose gel to find enrichment of a smear from 300-400 bp. A larger smear is acceptable and primer dimers are found at 150 bp. Once confirmed that both IRR and IRL reactions worked, samples were prepared for sequencing submission.

All the samples were pooled together and purified with the Geneaid PCR Cleanup kit (Geneaid, #DFC100). The final volume was 150-200 µL, with a final concentration of approximately 20-25 ng/µL for sequencing submission to TCAG. Paired end sequencing was completed on the NovaSeq SP flow cell v1.5 platform s 2×100 bp and with 15% PhiX spike-in.

#### Shear-SPLINK Insertions Analysis

Adaptors were trimmed with cutadapt (v1.8) with parameters *‘-m 5 --no-indels --discard-untrimmed -g R1_5prime=^NNNNNNNNTGTATGTAAACTTCCGACTTCAACTG’* from Read 1 (R1) of each sample. Only the reads that began with TA were kept for downstream analysis since the SB transposon inserts into a TA dinucleotide. Read 1 reads were then paired with their respective Read 2 reads and aligned with novoalign (v3.05.01) and the mm9 mouse genome assembly using *‘-r ALL 1 -R 0 -c 8 -o SAM’* parameters. The aligned files were then converted into BAM files for downstream analysis.

Each detected insertion was annotated using refFlat tables obtained from the UCSC genome database. The following details were extracted and recorded: tumor ID, gene name, the specific region of the gene affected, the predicted impact of the insertion on gene expression, the read count at the insertion site, and the orientation of the transposon relative to the gene. Insertions that did not occur within annotated genes were excluded from further annotation.

Clonality was estimated using the shear-SPLINK’s unique ligation point (ULP) score [39]. Each BAM file was converted into a bed file using samtools (v1.9) and then the length of each fragment was calculated by extracting the start coordinates of Read 1 and end coordinates of Read 2. From this, the unique fragment lengths such as ULP counts were calculated and appended to the list of insertions. A ULP score was calculated by dividing the insertion ULP count by the highest in each library. A score of 1 represented the most clonal insertion in a given library.

Insertion locations within 5 base pairs were stitched together as a cluster in the same library as they likely represented the same insertion. The insertion with the highest number of mapped reads was assumed to a true insertion location. The IRL and IRR libraries were merged and insertions that were detected in both libraries were used to proceed with filtering and gCIS analysis.

An insertion was filtered out if it was found in less than 3 biological libraries, had an ULP count of 1, had an ULP score less than 0.5, and was found on the donor chromosome (SB76 = chromosome 1 and SB68 = chromosome 15). If the insertion was in more than one merged library, the only insertion that was kept was the one with the highest read count. This served to ensure that no cross-contamination occurred between libraries pooled together.

For the gCIS analysis of shear-SPLINK data, a detailed explanation of this method can be found in Brett et al., 2011 [40]. This method calculated the probability of transposon insertions for each gene and compared the expected number of insertions to the observed number of insertions using a Chi-squared distribution to produce the results of a gCIS analysis. The p-values were adjusted with stringent Bonferroni group-wise correction and those that are < 0.05 are considered significant.

#### Proportions of Shared Insertions Analysis

The evolutionary trajectory of the primary tumor, micro-metastasis, and macro-metastasis was revealed through measures of similarity. The proportion of shared insertions between the three different malignant cell populations were calculated and then scored against each other. A higher score indicated a greater proportion of insertions shared between two populations. Significance was assessed with the Paired Wilcox test and p-values < 0.05 were significant.

#### Peeling the Leptomeninges

The procedure for extracting leptomeninges was adapted from Abeysundara et al. (2025) [17]. The leptomeninges of the spinal cord can be easily removed by peeling after imaging. In a petri dish filled with PBS, the nerve endings of the spinal cord are cut for ease of downstream processing and to avoid contamination of nerve tissue. Depending on the location of the spinal metastases, the leptomeninges and spinal cord are cut along the ventral or dorsal line with spring scissors (Fine Science Tools #91500-09) from the cervical to the sacral end. Fine forceps (Fine Science Tools, Dumont #5, #11254-20) were used to peel and strip the leptomeninges away from the spinal cord tissues from the cut opening. Peeled leptomeninges are placed into 1 mL PBS in Eppendorf tubes on ice until ready for downstream applications.

#### Dissociation of Leptomeninges

Peeled leptomeninges can be dissociated into single-cell suspensions with Liberase (Roche, #5401119001). Collected in PBS, the peeled leptomeninges were centrifuged at 300 rpm for 10 min. The leptomeninges were then incubated with 30 µL 13U/ml Liberase, 31.5 µL 2000U/1 ml DNase I (Roche, #11284932001) and 938.5 µL of PBS for 1 hr. at 37°C on a rocker and inverted every 15 min. After 1 hr. incubation, the leptomeninges were centrifuged at 300 rpm for 10 min. and the supernatants were discarded. Samples underwent myelin removal, 120 uL myelin removal beads (Miltenyi Biotec, #130-096-433) in 1 mL media were added to the dissociated leptomeninges and then incubated at 4°C for 15 min. on a rotator. After incubation, the samples were placed on a magnetic rack (Thermo Fisher, #12321D) for 3 min. and the supernatants were collected and centrifuged. The cell pellets were washed with PBS and then plated in cell media for downstream applications.

#### Micro- and Macro-Metastasis Re-Injection

Immunocompromised NSG mice were injected with 5, 000 GFP+ D425 cells or 5, 000 tdTomato+ D425 cells into the lateral ventricle as indicated above and allowed to progress towards human endpoint, in which then the spinal cord was collected. GFP+ and tdTomato+ micro- and macro-metastases were identified through the imaging analysis on the Nikon SMZ25 stereoscope and collected separately into 1 mL PBS, then placed on ice. The metastatic tissues were then dissociated with Liberase as described above, stained with 1: 1000 propidium iodide (Invitrogen, #P1304MP), and flow sorted from Sony MA900 cell sorter equipped with 488 nm, 405 nm, 561 nm lasers and corresponding bypass filters (525/30, 617/30, 450/50) into equal numbers of GFP+ micro-/ tdTomato+ macro- metastases in one tube and tdTomato+ micro- / GFP+ macro-metastases in a second tube. The sorted cells were immediately re-injected into secondary NSG animals such that one NSG received equal numbers of GFP+ micro-/ tdTomato+ macro-metastatic cells and another NSG animal received equal numbers of tdTomato+ micro- / GFP+ macro- metastatic cells in their lateral ventricles. The CNS of animals were collected at endpoint, in which then the spinal cord was imaged and analyzed for the presence of GFP+ or tdTomato+ metastases. Quantification of the number of GFP+ and tdTomato+ metastases were completed in ImageJ.

A similar experiment but only with GFP+ tumor cells was also completed. Immunocompromised animals were injected with 5, 000 GFP+ D425 cells and allowed to progress towards endpoint, in which then the GFP+ micro- and macro-metastases were collected and dissociated with Liberase. GFP+ micro- and macro-metastatic cells were then stained with 1: 1000 propidium iodide (Invitrogen, #P1304MP) and flow sorted into equal numbers but separated Eppendorf tubes. The sorted cells were immediately re-injected into the lateral ventricles of secondary NSG mice, such that one animal received GFP+ micro-metastatic cells and another received an equal number of GFP+ macro- metastatic cells. At the endpoint, the animals were sacrificed, the CNS was collected, and the spinal cord was analyzed for the presence and quantification of GFP+ metastases.

#### Bioluminescence imaging

Xenografted mice undergoing bioluminescent imaging were anesthetized with 4% isoflurane in an induction chamber and administered intraperitoneal (i.p.) injections of IVISBrite D-luciferin (150 mg/kg; PerkinElmer, Cat no. 122799). Imaging was performed five minutes post-injection using the Xenogen Spectrum (IVIS-Spectrum) imaging system. Bioluminescent signals were analyzed with Living Image Software v4.6 (PerkinElmer).

#### Lentiviral production and transduction

The following lentiviral plasmids were designed using VectorBuilder as follows: pLV[Exp]- CMV>TurboGFP(ns):T2A:Luc2(ns):F2A:Neo (GFP-Luc); pHIV-Luc-ZsGreen (Luc-ZsGreen) was a gift from Bryan Welm (Addgene plasmid #39196). Lentiviral plasmids were obtained through the Sigma MISSION TRC shRNA library with the following constructs: CXCL12 shRNA (TRCN0000262667) in the TRC2-pLKO-puro vector, SCARB1 shRNA (TRCN0000299698) in the TRC2-pLKO-puro vector, CYP11A1 shRNA (TRCN0000423133) in the TRC2-pLKO-puro vector, and the SHC201 TRC2 pLKO.5-puro empty vector control plasmid DNA. To generate lentiviral particles, 8 × 10⁶ HEK293T cells were seeded into a 15 cm adherent plate with 30 ml of DMEM supplemented with 10% FBS and incubated for 24 hours. Cells were co-transfected with 10 µg of the lentiviral plasmid, 7.5 µg pMDLg, 7.5 µg pRSV-Rev, and 5 µg pMD2.6/VSVG using the X-tremeGene 9 DNA Transfection Reagent (Roche, Cat no. 6365779001). Supernatants containing lentiviral particles were collected 72 hours post-transfection, concentrated using the Lenti-X concentrator (Clontech, Cat no. 631231), and incubated overnight at 4°C. The concentrated particles were pelleted by centrifugation at 1500 × g for 45 minutes, resuspended in 1/100th of the original volume with 1X PBS, and stored at -80°C. Medulloblastoma cell lines (D425, D458, MED8A) were transduced with lentiviral particles for 24 hours. Following transduction, cells were sorted based on GFP, mCherry positivity using viability dyes (PI or DAPI) on the Sony MA900 at the SickKids-UHN Flow Cytometry Facility.

#### Protocol Quantitative real time PCR

Total RNA was extracted using the RNeasy Mini Kit (QIAGEN, Germany; catalog numbers 74104 and 74106). Complementary DNA (cDNA) was synthesized following the manufacturer’s instructions using the SuperScript III First-Strand Synthesis System (ThermoFisher Scientific, USA; catalog number 11752). For each quantitative real-time PCR reaction, 5 ng of total cDNA was used. PCR amplification, data acquisition, and analysis were conducted on a QuantStudio™ 3 Real-Time PCR System (ThermoFisher Scientific, USA). GAPDH served as the internal control for normalization of gene expression.

#### sLP-mCherry Labelling system

A soluble peptide (SP)3 and a modified TAT peptide were cloned upstream of the mCherry cDNA under the control of a mouse phosphoglycerate kinase (PGK) promoter, as described previously [21]. The sLP-Cherry sequence was inserted into a pRRL lentiviral backbone to generate the expression construct. Lentiviral particles were produced in HEK293T cells using a standard third-generation packaging system. D425 GFP+ medulloblastoma cells were transduced with sLP-Cherry lentivirus at a multiplicity of infection (MOI) optimized for stable expression. Cherry+GFP+ cells were then isolated using fluorescence-activated cell sorting (FACS) for downstream analyses.

#### Fluorescence-activated cell sorting

##### Slp-mCherry+/GFP- cells

Brain tumors and microdissected spinal meninges were digested in 1 ml of 1X PBS containing 0.41 U/ml Liberase TM Research Grade (Roche, Cat no. 5401119001), 60 U/ml DNase I (Roche, Cat no. 11284932001), and 1X Collagenase/Dispase (Roche, Cat no. 10269638001) for 1 hour at 37°C on an orbital shaker. Following enzymatic dissociation, the samples were centrifuged at 300 × g for 10 minutes, and the supernatant was discarded. Brain tumor pellets were resuspended in fluorescence-activated cell sorting (FACS) buffer (1% BSA, 1 mM EDTA in 1X PBS) and filtered through a 40 µm strainer, while meningeal pellets were resuspended in 1 ml of FACS buffer. For myelin removal, meningeal suspensions were incubated with 120 µl of Myelin Removal Beads II (Miltenyi Biotec, Cat no. 130-096-433) at 4°C for 15 minutes with gentle rocking. The samples were then placed in a magnetic separator for 3 minutes, after which the supernatant was transferred to 1.5 ml tubes and centrifuged at 300 × g for 5 minutes. samples were washed once with FACS buffer and resuspended in FACS buffer DAPI (Sigma-Aldrich Cat no. MBD0015; 1:3000) to label dead cells. Cell sorting was performed using a Sony MA900 cell sorter equipped with 488 nm, 405 nm, and 561 nm lasers and corresponding bypass filters (525/30, 617/30, 450/50) at the SickKids-UHN Flow Cytometry Facility. Cells were identified as mCherry+/GFP⁻/+ cells, as outlined in Figure 6.

##### APC+/GFP⁻/PI- cells

For staining, following two rinses, meningeal samples were blocked in 100 µl of 0.5 mg/ml Rat anti-CD16/CD32 antibody (Thermo Fisher, Cat no. 14-0161-82) prepared in FACS buffer at 4°C for 10 minutes. Meningeal samples and the Anti-Rat Ig, κ/Negative Control Compensation Particle Set (BD Biosciences, Cat no. 552844) were incubated with APC-conjugated anti-mouse F4/80 antibody (Abcam, Cat no. ab105080) in FACS buffer for 30 minutes at 4°C. Antibody dilutions were 1:100 for compensation beads and 1:300 for the meningeal samples. After staining, samples were washed once with FACS buffer and resuspended in FACS buffer containing propidium iodide (PI, BD Biosciences, Cat no. 556463; 1:1000) to label dead cells. Cell sorting was performed using a Sony MA900 cell sorter equipped with 488 nm, 638 nm, and 561 nm lasers and corresponding bypass filters (525/30, 617/30, 660/20) at the SickKids-UHN Flow Cytometry Facility. F4/80⁺ macrophages were identified as APC+/GFP⁻/PI+ cells, as outlined in Figure 6.

##### Micro-Metastases and Macro-Metastases cells

Micro - and macro - metastatic tumor cells were collected from the spinal cords of *Math1- GFP/J2Q-SB11/T2Onc/Ptc Het* during imaging on the Nikon SMZ25/18 system. A macro - metastasis can be visualized easily under the green, fluorescent laser without use of the microscope. Spinal meningeal regions lacking grossly visible macrometastases were dissected, and areas containing GFP-positive micrometastatic were mechanically dissociated in 100 uL of enzyme - free media (Sigma - Aldrich, #S - 014 - B) in 96 - well plates prior to transfer into polypropylene FACS tubes. Cells were stained with propidium iodide (Invitrogen, #P1304MP) to differentiate between live and dead cell.

#### Single-Cell RNA Sequencing with SMART-Seq

Single-cell RNA sequencing was completed for micro-metastases, micro-metastases, and APC+/GFP-/PI- cells according to the manufacturer’s instructions for SMART-seq (Takara Bio USA, #634437). A master mix of FACS dispensing solution containing 10X lysis buffer, RNase inhibitor, ERCC spike-in, the 3’ SMART-seq CDS Primer II A and water was prepared for each well of a 96-well plate for cell collection. Fluorescence-activated cell sorting samples were sorted as single cells and collected into individual wells of the plate. The plate was centrifuged for 30 sec. to ensure all cells are submerged in the solution, and then were either processed immediately or stored at -80°C after flash-freezing. Each plate containing sorted primary tumor, micro-, and macro-metastatic cells were lysed during a 3 min. incubation at 72°C and then processed for a one-step first strand cDNA synthesis and amplification. The master mix was composed of nuclease-free water, one-step buffer, SMART-seq HT oligonucleotide, RNase inhibitor, SeqAmp DNA polymerase and SMARTScribe reverse transcriptase. The PCR program was set to 42°C for 90 min., 95°C for 1 min., and then 18 cycles of 98°C for 10 sec, 65°C for 30 sec., and 68°C for 3 min., before finishing with 72°C for 10 min and 4°C on hold. After cDNA synthesis and amplification, the cDNA was purified with SPRIselect beads (Beckman Coulter, #B23318). The cDNA was incubated with 25µL of SPRIselect beads for 8 min. at room temperature, then washed twice with 200 µL of 80% ethanol for 30 sec. and placed on the 96-well magnetic stand (Alpaqua, #A001322). The cDNA-bead pellet was briefly dried, then rehydrated with 17 µL of elution buffer for 2 min. and collected into a low-adhesion 96-well plate for quantification library preparation.

Quantification of the amplified cDNA was completed with the Quant-iT 1X dsDNA assay kit (Invitrogen, #Q33232), according to the manufacturer’s instructions. The cDNA of each individual cell was barcoded with the Nextera XT DNA Library Prep kit (Illumina, #FC-131- 1096) as instructed by the user manual. The cDNA was tagmented, which is a process that fragmented and tagged the DNA with adapter sequences, and then amplified with a limited cycle PCR that added the unique i7 and i5 adapters required for barcoding each sample. The barcoded libraries were purified with SPRIselect beads following the same protocol as previously completed and then stored at -80°C until ready for sequencing. The Micro/Macro-Metastases libraries were submitted to the Ontario Institute of Cancer Research (OICR) for sequencing on the HiSeq2500 system on Rapid Run mode with a read length of 2 x 125 bp. Each batch of slp-mCherry+/GFP- cells and slp-Cherry-/GFP- cells were sequenced on the NovaSeq SP flowcell with 325 - 400 million reads (Paired End 150)

##### Alignment of SMART-Seq Raw Reads

Adapter sequence was trimmed using Trim Galore (v0.4.4) with the parameters ‘-- stringency 10 --clip_R1 13 --clip_R2 13 --length 36’ and the trimmed reads were subjected to quality control using FastQC (v0.11.5) with default parameters. Trimmed reads were then aligned to a custom reference based on the GRCm38 (mm10) reference genome with ERCC spike-in sequence appended. Alignment was performed using STAR (v2.5.4b) with the parameters ‘--outFilterMultimapNmax 20 --alignSJoverhangMin 8 --alignMatesGapMax 200000 --alignIntronMax 200000 --chimSegmentReadGapMax parameter 3 --alignSJDBoverhangMin 10 --alignSJstitchMismatchNmax 5 -1 5 5 --outSAMmultNmax 20 --chimSegmentMin 12 --chimJunctionOverhangMin 12 --twopassMode Basic’.

##### Quality Control and Normalization of SMART-Seq Data

Computational analysis was performed with R (v4.0.3). Quality control was performed using the R package scater [41]. (v1.0.4). Briefly, low-quality cells were identified on the basis of outlier (defined using 3 median absolute deviations) counts and mitochondrial gene content. Genes expressed in less than 10 cells were removed in addition to pseudogenes. The remaining high-quality cells were normalized using the R package scran [42] (v1.16.0). Briefly, size factors were calculated using the function *computeSpikeFactors* based on the ERCC spike-in counts followed by the *normalize* function.

##### Clustering Analysis and Visualization of SMART-Seq Data

Downstream clustering and visualizations were performed using the R package Seurat [43] (v3.1.0). scRNA-seq data was processed with the Seurat (v4.0.0) pipeline with Harmony (v1.0) batch effect correction. Briefly, the top 3000 highly variable genes were calculated, PCA analysis was performed to determine statistically significant PCs which were then used to perform uniform manifold approximation and projection (UMAP) dimensionality reduction. Clusters of transcriptionally similar cells were determined using Seurat’s shared-nearest neighbor algorithm. Differential expression analysis was performed with Seurat (v4.0.0) and pathway enrichment with gProfier2(v0.2.1).

##### Pathway enrichment analysis

Pathway enrichment analysis was performed using the ActivePathways method [44] using the top 100 p-values of genes identified as differentially expressed in slp-mCherry+ and slp-mCherry-macrophages. Pathway enrichment analysis was performed using the ActivePathways method [44] using the complete set of p-values of all genes identified as differentially expressed in micro- and macro-metastases, as described above. Mouse gene sets were retrieved from two biological databases, Gene Ontology (GO) [45] and Reactome [46] by downloading them in the GMT (Gene Matrix Transformed) format from the g:Profiler web server (version e111_eg58_p18_f463989d, downloaded January 25, 2024) [47]. For a stringent high-confidence analysis, we used a background set of 11,318 mouse genes that were both included in the differential gene expression analysis and had at least one annotation in GO or Reactome. Gene sets were size filtered to include only those containing 50 to 250 genes to reduce multiple testing and focus on a set of more specific and interpretable pathways and processes. Enriched pathways were selected using Benjamini-Hochberg false discovery rate (FDR) as the multiple-testing correction method. Otherwise, default parameters were used in ActivePathways. Significantly enriched pathways were selected (FDR < 0.05) and subsequently visualized as an enrichment map in the Cytoscape software according to standard protocols [48, 49]. For each detected pathway or process, associated genes from differential gene expression data were visualised to display their log2-transformed fold-change values and significance values of differential expression.

#### Bulk RNA sequencing

shControl D425 and shSCARB1 D425 cells were collected from their respective xenograft primary tumors (GFP +ve) at the endpoint. RNeasy Mini Kit (QIAGEN, Germany, cat nos. 74104 and 74106) that allows high quality RNA extraction from samples with low cell numbers (<10,000 cells), were performed on the samples, and RNA quality for each sample was assessed using the Agilent RNA 6000 Pico Kit (Agilent Technologies, USA). RNA was amplified with the NEB Ultra II Directional polyA mRNA prep and analysed at the Centre for Applied Genomics and the Hospital for Sick Children. The sequencing was performed on biological triplicates for each condition generating approximately 30 million 150 paired end reads.

#### Slingshot Analysis

Pseudotime trajectory analysis was performed using the function *slingshot* from the R package slingshot [50] (v1.6.1) without specifying a root cluster in order to unbiasedly determine the direction of cellular evolution.

#### Mutation Calling and Phylogenetic Analysis

Mutation calling was performed on an individual cell basis using GATK’s Best Practices for RNAseq [51]. Briefly, STAR aligned BAM files were cleaned using GATK’s (v4.0.1.2) *MarkDuplicates* function to sort and remove duplicate reads followed by the function *SplitNCigarReads*. Variant calling was performed using the function HaplotypeCaller with parameters ‘*--dont-trim-active-regions true --dont-use-soft-clipped-bases true --standard-min-confidence-threshold-for-calling 20.0*’. The resulting variants were filtered using the function VariantFiltration with the parameters ‘*-filter-name FS -filter \“FS > 60.0\” -filter-name QD -filter \“QD < 2.0\” -filter-name MQ -filter \“MQ < 40.0\” -filter-name ReadPosRankSum -filter \“ReadPosRankSum < -8.0\“*’ and annotated using ANNOVAR [52] (v2018.04.17). As our objective was to infer the degree of genetic relationship between metastatic cells arising from a given primary tumour, we further filtered the variant calls to retain only variants detected in at least 2 cells and with a minimum variant allele frequency of 0.05. Finally, to construct a variant-based phylogeny we utilized the R package DENDRO [53] (v0.2.2).

#### Untargeted Shotgun Lipidomics

*Math1-GFP/SmoA1/Nestin-SB100/T2Onc2* mice were sacrificed with CO2 at a humane endpoint and the CNS was collected as described above. Using the Nikon SMZ25 imaging system, micro- and macro-metastases were identified and isolated with fine forceps (Fine Science Tools, Dumont #5, #11254-20) and placed into separate Eppendorf tubes filled with 1 mL PBS. A small piece of the primary tumor was also collected into an Eppendorf tube and placed on ice. The samples were briefly centrifuged to pellet the tumor cells and the PBS was removed prior to being flash frozen in a dry ice and ethanol bath. The samples were then stored at -80°C until ready for shipment to the Cayman Chemical Company (Michigan, US) for untargeted shotgun lipidomics mass spectrometry and analysis.

The lipidomics analysis on breast cancer cell line MDA-MB-23145 was completed similarly. The breast cancer cell line was graciously provided by Dr. Adrienne Boire’s laboratory (Memorial Sloan Kettering Cancer Centre, New York, US). Immunocompromised NSG mice were injected with 50,000 MDA-MB-231 cells into the fourth mammary fat pad. At the endpoint, the mice were sacrificed and the CNS was collected. Samples of micro-and macro-metastases were processed as above and frozen at -80 °C until ready for shipment. These samples were analyzed with untargeted shotgun lipidomics at Creative Proteomics (New York, US).

#### Immunofluorescence (IF)

The Immunofluorescence protocol was obtained and performed as described from Abeysundara et al., 2025 [17]. Mice were intracardially perfused with 30 ml of 1X PBS prior to immunofluorescence (IF) experiments. Primary tumor and spinal cord tissues were fixed in 4% paraformaldehyde at 4°C for 24 hours, followed by five washes in 1X PBS. The samples were sequentially immersed in 15% sucrose in 1X PBS for at least 24 hours at 4°C and then in 30% sucrose in 1X PBS for an additional 24 hours at 4°C. Afterward, tissues were embedded in OCT compound on dry ice, and the blocks were stored at -80°C until sectioning. Cryosections were prepared at a thickness of 16 µm, and slides were stored at -80°C until use for IF procedures. For IF staining, slides were air-dried at room temperature for 10 minutes, and sample borders were outlined using the ImmEdge Hydrophobic Barrier PAP pen (Vector Laboratories, Cat no. H-4000). Subsequent staining steps were conducted within an immunostaining moisture chamber. Tissue sections were washed three times (5 minutes each) in PBSp (1X PBS containing 0.1% saponin; Sigma-Aldrich, Cat no. 47036-50G-F) and blocked for at least 1 hour in a solution of 10% goat or donkey serum with 0.1% saponin in 1X PBS. Primary antibodies were diluted in the blocking solution and applied to the samples, which were incubated overnight at 4°C. The following day, tissues were washed six times (5 minutes each) with PBSp. Secondary antibody staining was performed at room temperature for 1 hour using the same blocking solution, followed by six additional washes (5 minutes each) in PBSp. Nuclear counterstaining was achieved by incubating the sections in DAPI (1:1000; Sigma-Aldrich, Cat no. MBD0015) for 5 minutes. Finally, samples were rinsed three times in 1X PBS and mounted using Dako mounting medium (Agilent, Cat no. S302380-2).

For whole mount staining procedure, meninges were isolated from spinal cords through microdissection and fixed in 4% paraformaldehyde for 20 minutes. Following fixation, the tissues were rinsed thoroughly with 1X PBS at least five times and stored at 4°C. Prior to staining, samples were rinsed once in 1X PBS and blocked for at least 1 hour at room temperature in a solution containing 5% BSA and 0.1% saponin in 1X PBS, with gentle rocking. Primary antibody staining was carried out by incubating the meninges overnight at 4°C in 1 ml of the respective antibody solution. Afterward, the samples were washed extensively in PBBSp (1X PBS supplemented with 1% BSA and 0.1% saponin) for a minimum of 1 hour. Secondary antibodies were then applied for 2 hours at room temperature, using 1 ml of the secondary antibody solution. The tissues were subsequently washed again in PBBSp for at least 1 hour and incubated with DAPI (1:1000; Sigma-Aldrich, Cat no. MBD0015) for 5 minutes to counterstain nuclei. Finally, the meninges were rinsed three times in 1X PBS and mounted using Dako mounting medium (Agilent, Cat no. S302380-2).

##### Ki67 Staining

Proliferation by Ki-67 staining was completed following the protocol for immunofluorescence, as described above with a few changes due to the use of the Ki-67 antibody. The process of fixation, blocking, and primary antibody incubation were completed in the same manner. The Ki-67 mouse antibody (Invitrogen, #50-5698-82) is conjugated to Alexa Fluor 660 and hence, when used at 1:1000 dilution, no secondary antibody was required. Following the overnight primary antibody incubation, the samples were washed and stained with DAPI as usual, then imaged on the Nikon A1R system. Quantification for Ki-67 staining was completed in ImageJ through the analysis of overlapping GFP+ and Ki67+ areas. For interrogating the microenvironment, the quantification focused on the Ki67+ / GFP-populations and area overlap.

##### *In vivo* Labeling of EdU

*In vivo* labeling with EdU for assessing proliferation of micro- and macro-metastases were completed according to the kit’s user manual from the manufacturer (Sigma-Aldrich, # BCK594-IV-IM-S). GEMMs that reached a human endpoint were injected intraperitoneally with 100 mg / kg of body mass of 10 mM EdU approximately 24 hr. prior to sacrifice. Animals were sacrificed with CO2 and the CNS was collected as described. The leptomeninges were peeled and then fixed with 1 mL 4% PFA (Fisher Scientific, #50-980- 487) for 20 min. at room temperature. After fixation, the samples were washed twice with 1 mL of 3% BSA (Sigma-Aldrich, #A9647-100G) in PBS and permeabilized with 1 mL of 0.5% Triton X-100 (Sigma-Aldrich, #T8787-100ML) in PBS for 20 min. at room temperature. As the samples were incubating, the reaction cocktail for EdU detection was prepared in the following order: 379 µL deionized water, 50 µL 10X Reaction Buffer, 20 µL Catalyst Solution, 1 µL 10 mM 6-FAM-Azide, and 50 µL 10X Buffer Additive. Once permeabilization was completed, the tissues were washed twice with 1 mL 3% BSA in PBS and 500 µL of the reaction cocktail was added. The samples were protected from light and incubated at room temperature for 30 min. After this incubation, the reaction cocktail was removed and the samples were washed 3 times with 1 mL 3% BSA in PBS. The samples were then subjected to immunofluorescent staining as described above for GFP detection and DAPI nuclear staining. The slides were imaged on the Nikon A1R system.

Quantification for EdU detection was completed in ImageJ. The area of overlap between GFP+ and EdU+ staining informed micro- and macro-metastatic tumor cells that were undergoing proliferation.

##### Apoptosis Staining with TMR Red

Peeled leptomeninges harbouring micro- and macro-metastases were fixed with 4% PFA (Fisher Scientific, #50-980-487) for 20 mins at room temperature on a rocker. Staining for TUNEL as a marker of apoptosis was completed according to the manufacturer’s manual for the In Situ Cell Death Detection Kit, TMR Red (Roche, #12156792910) following the labeling protocol for tissues adherent cells, cell smears, cytospin preparations, and tissues.

Prior to staining, the fixed leptomeninges were washed 3 times with PBS for 30 min. on the rocker and then treated with 1 mL permeabilization solution (0.1% Triton-X-100 (Sigma-Aldrich, #T8787-100ML) and 0.1% sodium citrate (Bio Basic Canada, #CB0035), freshly prepared) on ice for 10 min. The samples were then rinsed with PBS and 50 µL of the TUNEL reaction mixture (50 µL enzyme solution with 450 µL label solution) was added. Since the leptomeningeal tissue is free floating in the Eppendorf tube, the 50 µL volume should be sufficient to cover the tissue. The samples were then covered with aluminum foil and placed inside a 37°C incubator for 1 hr.

Once staining was completed, the leptomeninges were rinsed thrice with PBS, then counterstained with DAPI for 5 min. at room temperature and washed 3 times with PBS. The samples were mounted onto glass slides with Dako Mounting Medium (Agilent Technologies, #S3023) and stored at 4°C in the dark until imaged on the Nikon A1R confocal microscope system. Quantification for apoptosis was completed in ImageJ by analysing the area of GFP+ and TUNEL+ staining that overlapped in micro- and macro-metastases.

##### Oil Red O Staining

The Oil Red O (ORO) protocol was adapted from Mehlem et al., 2013. for staining neutral lipid droplets in the leptomeninges. ORO was purchased from Millipore Sigma (Sigma-Aldrich, #O0625-100G) and an ORO stock solution was made by adding 2.5 g ORO to 400 mL of 99% v/v 2-propanol (Sigma-Aldrich, #190764-4L). The solution was mixed by magnetic stirring for 2 hr. and stored at room temperature. An ORO working solution was made by adding 1.5 parts of ORO stock solution to one part of ddH_2_O (Wisent, #809-115-CL). The solution was then placed in 4°C for 10 min. to thicken and filtered through a 45 µm filter to remove precipitates. The working solution was used within 6 hr. of preparation.

The leptomeninges harbouring micro- and macro-metastases were fixed as free-floating tissue in 1 mL of 4% PFA (Fisher Scientific, #50-980-487) for 20 min. on a rocker at room temperature. The tissue was washed 3 times with PBS and stored at 4°C in PBS until ready for ORO staining.

Tissues were brought to room temperature prior to ORO staining. 1 mL of ORO working solution was added to each tube containing metastatic leptomeninges and incubated at room temperature for 10 min. on a rocker for equal distribution of ORO. After 10 min. incubation, samples were washed with water 3 times for 10 min. each before mounting onto a slide with the Dako fluorescence mounting medium (Agilent Technologies, #S3023). The slides were imaged on the Nikon TE-2000 Epifluorescence Microscope to capture both brightfield staining of ORO and fluorescence from micro- and macro-metastases.

Captured images that overlay ORO staining with GFP+ micro- and macro-metastases were quantitatively analyzed in ImageJ, similarly to Mehlem et al., 2013 [54], for the pixel area of each ORO stain associated with nearby micro- and macro-metastases. For GFP pixel area quantification, we used the Volocity software and selected the ‘Measure’ function. Using the commands ‘Find, Measure, and Analyze Objects,’ we specifically selected the GFP object for evaluation. We then used the output, ‘sum of all pixel count,’ to determine the GFP-positive pixel area.

##### Filipin Staining

Filipin complex (Sigma, Cat no. F-9765) was prepared as a 25 mg/ml stock solution in DMSO and diluted to a working concentration of 0.05 mg/ml in PBS supplemented with 10% FBS. Meningeal samples were first rinsed thoroughly with 1X PBS at least five times and stored at 4°C until processing. Before staining, the tissues were rinsed once in 1X PBS and incubated with 1.5 mg/ml glycine in PBS for 10 minutes at room temperature to neutralize residual paraformaldehyde. Samples were then blocked in a solution containing 5% BSA and 0.1% saponin in 1X PBS for a minimum of 1 hour at room temperature with gentle rocking. Filipin staining was performed by incubating the meninges with 1 ml of the working solution for 2 hours at room temperature. Following staining, the tissues were washed three times with PBS and imaged in PBS using fluorescence microscopy equipped with a UV filter set (excitation: 340–380 nm, dichroic mirror: 40 nm, emission: 430 nm long-pass filter). For filipin staining tissues, SYTO deep red nucleic acid stain (1:2000) were used instead of DAPI.

#### Microscopy and Image analysis

##### Imaging with the Confocal microscopy

Whole mounted leptomeninges that were stained with antibodies for immunofluorescence were prepared for confocal imaging on either the Quorum Leica DMi8 spinning disk or Nikon A1R system. Quorum Leica DMi8 spinning disk confocal microscope is equipped with a Photometric Prime 95B camera and a 63× objective lens. Image acquisition was performed using Volocity software. All imaging were carried out at the SickKids Imaging Facility, and data were subsequently analyzed using Imaris Bitplane 10.1.0, as detailed below.

Nikon A1R system was set for resonant scan imaging and DAPI-GREEN-RED was selected. To capture a large image of the whole leptomeninges, ND Acquisition with ‘Z-Series’ and ‘Large Image’ were used. After adjusting the laser power and gain for detection of DAPI, green, and red fluorescence. The typical range for the Z-stack range of the meninges is 50-100 µm, but every tissue and its mount varied. Next, the area of the leptomeninges was determined in the ‘Large Image’ tab. The typical range for the peeled leptomeninges was approximately 12 x 40. Stitching was completed with 10% overlap between each image. To reduce noise, each image reported the average of 4 scans. Each slide with whole mounted leptomeninges would take approximately 3-5 hrs. to image and process. Deconvolution was completed by the software after the image was taken and exported for downstream analysis in Imaris (Oxford Instruments).

##### Proportion of Adgre1+ Nr1h3 cells

Using Imaris, the number of DAPI, GFP, and LXRa cells were quantified using the “Spots” function and background subtraction with the following parameters for DAPI (Estimated diameter of 10µm; Quality threshold above 0.101), GFP (Estimated diameter of 15µm; Quality threshold above 2.42), and LXRa (Estimated diameter of 3µm; Quality above 4.30). The F4/80 volume was quantified using the “Surface” function with smoothing enabled, surface grain size of 0.260µm, absolute intensity, and Enabled Automatic Threshold = true. The number of DAPI spots close to the F4/80 surface (F4/80+ cells) was quantified using the “Find Spots Close to Surface” tool with a threshold set to 5µm. The number of LXRa spots that collocated with the DAPI spots close to F4/80 (LXRa+ F4/80+ cells) was quantified using the “Colocalize Spots” tool using a threshold of 5µm. The proportion of LXRa+ F4/80+ cells was determined by dividing the number of LXRa+ F4/80+ cells by the total number F4/80+ cells and multiplying by 100.

##### Filipin+ cells/Total cells

Using Imaris, the number of SYTO and GFP cells were quantified using the “Spots” function and background subtraction with the following parameters for SYTO (Estimated diameter of 5µm; Quality threshold above 1.85) and GFP (Estimated diameter of 5µm; Quality threshold above 63.5). The Filipin volume was quantified using the “Surface” function with smoothing enabled, surface grain size 0.100µm, absolute intensity, and Threshold value of 394.41. The number of GFP- SYTO+ cells (non-tumor cells) was quantified by subtracting the number of GFP spots from the total number of SYTO spots. The number of GFP spots and SYTO spots close to Filipin (Filipin+ GFP+ cells and Filipin+ SYTO+ cells, respectively) was quantified using the “Find Spots Close to Surface” tool with a threshold set to 5µm. The number of non-tumour cells close to Filipin (Filipin+ non-tumor cells) was calculated by subtracting the number of Filipin+ GFP+ cells from the total number of Filipin+ SYTO+ cells. The number of Filipin+ (GFP or non-tumor) cells was divided by the total number of GFP or non-tumor cells, respectively.

##### SCARB1+ GFP+/Total GFP Volume

Using Imaris, the volume of GFP and SCARB1 was quantified using the “Surface” function with smoothing enabled and the following parameters for GFP (Surface grain size of 0.260µm; Absolute intensity; Enabled Automatic Threshold = true) and SCARB1 (Surface grain size of 0.260µm; Absolute intensity; Threshold of x1.63). The SCARB1+ GFP+ co- localization volume was quantified using the “Surface-Surface coloc” tool with no smoothing, which was divided by the total GFP volume.

##### Adgre, GFP, and Abcg1 co-localization analysis

Using Imaris, the number of GFP cells was quantified using the “Spots” function with background subtraction and the following parameters: Estimated diameter of 15µm and Quality threshold above 0.402. The F4/80 and ABCG1 volume was quantified using the “Surface” function with smoothing enabled, absolute intensity and the following parameters for F4/80 (Surface grain size 0.260µm, Threshold value of 309.923) and for ABCG1 (Surface grain size 0.260µm, Threshold value of 217.794). The number of GFP spots close to F4/80 was quantified using the “Find Spots Close to Surface” tool with threshold set to 10 µm and divided by the total number of GFP spots (GFP cells close to F4/80 normalized to GFP cells). The F4/80+ ABCG1+ co-localization volume was quantified using the “Surface-Surface coloc” tool with no smoothing, which was divided by the total F4/80 volume.

##### Cyp11a1+ cells/Total cells

Using Imaris, the number of DAPI and GFP cells were quantified using the “Spots” function and background subtraction with the following parameters for DAPI (Estimated diameter of 5µm; Quality threshold above 0.5) and GFP (Estimated diameter of 5µm; Quality threshold above 13). The Cyp11a1 volume was quantified using the “Surface” function with smoothing enabled, surface grain size 0.100µm, absolute intensity, and Threshold value of 50. The number of GFP- DAPI+ cells (non-tumor cells) was quantified by subtracting the number of GFP spots from the total number of DAPI spots. The number of GFP spots and DAPI spots close to Cyp11a1 (Cyp11a1+ GFP+ cells and Cyp11a1+ DAPI+ cells, respectively) was quantified using the “Find Spots Close to Surface” tool with a threshold set to 5µm. The number of non-tumour cells close to Cyp11a1 (Cyp11a1+ non-tumor cells) was calculated by subtracting the number of Cyp11a1+ GFP+ cells from the total number of Cyp11a1+ DAPI+ cells. The number of Cyp11a1+ (GFP or non-tumor) cells was divided by the total number of GFP or non-tumor cells, respectively.

##### Quantification of Cell Numbers, Area, and Volume

Large scan, Z-stack images of peeled leptomeninges were imported into Imaris (Oxford Instruments) and consequently converted into the IMS format. The images were opened in 3D view and the visualization of DAPI, GFP tumor cells, and staining of interest were adjusted using ‘Display Adjustment’. For measuring the volumes of micro- and macro- metastases, metastases on the leptomeninges were created into ‘Surfaces’ within the green channel. The ‘Surface Detail’ (ranged from 2.5-5.0 µm) and ‘Threshold’ (ranged from 200-400) were adjusted to best reflect the images. The volumes and other information regarding the metastases were exported from ‘Statistics’. For measuring the number of cells in micro- and macro-metastases, the surfaces of metastases were separated into micro- and macro-metastases based on their volume (>100, 000 µm3) and ‘Spots’ were created to identify DAPI stained nuclei, with a set diameter of 5 µm, which was measured in ‘Slice’ View. Then, using the Imaris function for ‘Spots Close to Surface’ and a threshold of 0 µm, the number of DAPI ‘Spots’ were counted in micro- and macro-metastases. Area was also measured in ImageJ by outlining the micro- and macro-metastases in each image.

#### Quantification of the metastatic burden

Quantification of metastatic burden in spinal cord images was performed using a custom analysis pipeline available on GitHub (Balin et al., 2023). The metbuRden program is designed to measure metastasis area from microscopy images and can be found at https://github.com/PolinaBalin/metbuRden. Images were acquired with Nikon SMZ25/18 system. Green channel tiff images were rasterized using brick function of the raster package (v_3.6-20). Green pixel intensity values were then isolated. Density function was run on the green pixel intensity using bandwidth of 0.5.

Spine area was isolated by subtracting the background fluorescence. Local minima and maxima were calculated using turnpoints function in pastecs package (v_1.3.21). First pit value was recorded to subset the pixels above it. The number of pixels above the pit value was denoted as the spinal area. Only pixels above the first pit value were used for metastatic burden analysis to ensure that only metastatic areas on the spine were captured.

To check whether there are any outliers in the spinal area selection, geometric mean and geometric standard deviation of the spinal area were calculated across all mice of the given experiment using EnvStats package (v_3.0.0). Threshold for outliers was calculated by multiplying geometric mean by geometric standard deviation and by a factor of 1.25. Spinal images where spinal area exceeded the calculated threshold were removed from the analysis.

Next, we examined whether there was contrast variation across images. If there was, images were divided into high versus low contrast images. Threshold for low contrast images was determined by calculating 0.25 quantile of the standard deviation of the green pixel intensity across all mice of a given experiment. Images with standard deviation of the green pixel intensity below the threshold were considered low contrast, while the ones above the threshold were considered high contrast.

To calculate the metastatic signal threshold per image, mean of the green pixel intensity per image was added to the standard deviation of the green pixel intensity per image multiplied by an index. The multiplier index was determined manually per experiment. If images were divided into low versus high contrast, different multiplier indices were selected. Additionally, if some images showed highly extensive metastatic coverage, the multiplier index for these images was capped at a threshold. Capping threshold was calculated by adding 0.75 quantile of the mean green pixel intensity per image to interquartile range of the mean green pixel intensity per image multiplied by 1.5. The number of pixels above the selected thresholds was recorded, indicating metastatic area.

A threshold for metastatic area outliers was determined by summing up the mean of metastatic areas per experimental arm and standard deviation of metastatic areas per experimental arm multiplied by 3. Images where metastatic area exceeded the threshold were flagged and examined. In cases when an artificial glare on a spine was captured instead of a metastatic lesion, the image was removed from the analysis.

Spinal area and metastatic area values were then summed up per mouse, a fraction of metastatic area divided by spinal area was then calculated. Statistics was performed using Mann-Whitney U test.

#### In Vivo [^13^C] Glucose Tracing

Immunocompromised NSG mice were injected with D425 MB cells as described above. At endpoint, mice were anesthetized with isoflurane and tail vein access was achieved with a 23-gauge butterfly needle (BD, #367297). ^13^C -labeled glucose (Cambridge Isotope Laboratories, #CLM-1396) was infused as a bolus of 0.4125 mg / g of body mass in 125 µL saline for over a minute, followed by a 3-hr. continuous infusion of 0.008 mg / g of body mass per minute (150 µL per minute). Blood samples were collected prior to bolus infusion, and then at the 30 min., 60 min., 120 min., and 180 min. marks. At the end of the infusion period, cardiac perfusion with PBS was performed and the mouse was euthanized. The primary tumor was immediately dissected and flash frozen in a dry ice and ethanol bath. The spinal cord was rapidly dissected with care to keep the leptomeninges intact and cooled on a bed of ice. Micro- and macro-metastases were identified by fluorescent microscopy, dissected, and flash frozen with the dry ice and ethanol bath.

To prepare samples for mass spectrometry, samples were suspended in 1 mL of 80: 20 v/v methanol and water solution and disrupted with a homogenizer. The homogenate was then vortexed and 200 µL was transferred into a tube with 800 µL of ice cold 80: 20 v/v methanol and water solution. This mixture was vortexed rigorously prior to centrifugation at 20,000 g for 15 min. in a refrigerated tube. The samples were then dehydrated without heat using a SpeedVac and stored at -80°C before shipment to the DeBerardinis Lab for mass spectrometry and analysis (University of Texas Southwestern Medical Center, US).

#### MALDI Imaging

To minimize the salt content that may interfere with MALDI TOF MS, non-perfused leptomeninges from *Math1-GFP/Ptch+/Math1-SB11/T2Onc* mice were preferred for this application. After imaging with the Nikon SMZ25/18 to identify micro- and macro-metastases, the leptomeninges were peeled from the spinal cord. Extra care was taken to maintain a specific orientation so that metastatic tumor cells will be near the surface when transferred onto the pre-cooled conductive side of indium tin oxide (ITO)-coated glass slides (Bruker Daltonics, #1868957) and flash frozen on dry ice. Samples were then desiccated for 45 min. and stored at -80°C until ready for shipment to the City University of New York for MALDI TOF MS imaging analysis.

The paired experimental mouse brains were collected and rapidly frozen for 5 minutes using an aluminum tray suspended over liquid nitrogen. Once frozen, the tissue was cryosectioned into 10 μm thick slices at −20°C, with both the specimen head and cryostat chamber maintained at this temperature. The resulting sections were carefully placed onto the pre-chilled conductive surface of ITO-coated glass slides. Slides were desiccated under vacuum at room temperature for 45 minutes. After drying, these slides were stored at -80°C until ready for shipment to the City University of New York for MALDI TOF MS analysis.

The MALDI TOF MS imaging was carried out in the MALDI MS Imaging Joint Facility at the Advanced Science Research City of City University of New York. High purity grade N (1- Naphthyl) Ethylenediamine Dihydrochloride (NEDC), and Phosphorus (red) were purchased from Millipore Sigma-Aldrich (USA). Optima UHPLC/MS-grade methanol, isopropanol and water were purchased from Fisher Scientific (USA).

NEDC matrix solution of 10 mg/mL in isopropanol/water (70/30, v/v) was deposited at a flow rate of 0.05 ml/min and a nozzle temperature of 80°C for 30 cycles with no drying between each cycle. A spray velocity of 1300 mm/min, track spacing of 2 mm, N2 gas pressure of 10 psi, flow rate of 3 L/min and nozzle height of 40 mm were used.

MALDI mass spectra were acquired in negative ion mode for NEDC by MALDI TOF MS, Autoflex (Bruker Daltonics). MS spectra were 1) externally calibrated using red phosphorus as the standard for all experiments; 2) internally calibrated for higher accuracy based on the previously identified metabolites. The laser spot diameters were focused to “small” modulated beam profile for 20 μm raster width for leptomeninges and “medium” beam with 60 μm raster for brains. For each array position, imaging data were acquired by summing 500 laser shots at a repetition rate of 500 Hz. To prevent ion peak broadening, all measurements were conducted using the lowest possible laser power that still allowed for the collection of spectra with an adequate signal-to-noise ratio. Mass spectra were recorded across an m/z range of 40 to 1200, with a low mass cutoff set at 40 Da. Data acquisition and initial processing were performed using FlexImaging version 3.0, followed by further analysis in SCiLS (version 2015b). Ion images were generated using root-mean-square (RMS) normalization and a bin width of ±0.10 Da. Spectral interpretation was carried out manually, and analyte identification was supported by comparison with results from LC-MS experiments.

Following the experiment, the MALDI slides were promptly retrieved and rinsed with 95% ethanol. Standard hematoxylin and eosin (H&E) staining was then performed. The stained slides were imaged using a Leica Aperio CS2 slide scanner, and the resulting H&E images were used as the anatomical reference for mass spectra imaging (Figure S4A).

For LC-MS/MS experiments, cold methanol or cold methanol and isopropanol method was used to extract metabolites. Briefly, brain tissues were snap frozen in liquid nitrogen and approximate 30 mg of tissue was homogenized in 1mL cold methanol/water (80/20, v/v) or cold methanol/isopropanol/water (3:3:2 ratio) pre-cooled at -20°C. Following 10 min of gentle sonication in Bioruptor (30 s on, 30 s off, 10 cycles) at 4°C, samples were centrifuged for 10 min at 10,000 g at 4°C. and 20 μl solution was subjected to LC-MS/MS experiments. Comprehensive metabolic profiling was conducted on all tissue samples using ZIC-HILIC chromatography, employing a solvent system composed of acetonitrile, water, and 7 mM ammonium acetate. High-resolution mass spectrometry was performed using a maXis-II-ETD UHR-ESI-Qq-TOF instrument (Bruker Daltonics Inc.), coupled with a Dionex Ultimate-3000 liquid chromatography system. Each LC-MS/MS run was carried out in duplicate. Data analysis was performed using MetaboScape and XCMS Online platforms [55], which reference the METLIN and Human Metabolome databases. Metabolite identification was based on both high-accuracy mass measurements (within 5 ppm) and MS/MS fragmentation spectra. The presence of metabolites was compared between micro- and macro-metastases, as well as normal leptomeninges, in each and across individual animals.

#### High-Fat Diets

*Math1-GFP/Ptch+ and Math1-GFP/SmoA1/Nestin-SB100/T2Onc2* animals were genotyped and fed either a high-fat chow (9% fat) (TD2919, Envigo) or control diet (TD2918X, Envigo, Figure S8D) after weaning until their humane endpoint, in which then the central nervous system was collected and analyzed through imaging with the Nikon SMZ12/18 system and quantification with ImageJ of metastases. At the endpoint, the central nervous system was collected and analyzed for changes in metastatic burden through imaging with the Nikon SMZ12/18 system and quantification with ImageJ.

#### Cell-Cell communication Inference

To generate a single cell RNA-seq reference dataset for ligand receptor interaction in developing human fetal cerebellum, we utilized a published human cerebellar development dataset^26^ and subsetted rhombic lip derived glutamatergic lineage populations. We used semi-supervised and unsupervised opensource cell interaction R packages CCInx (v.0.5.2) and CellChat (v.2.1.2) respectively, which integrate comprehensive signaling molecule interaction databases that incorporate ligand-receptor complexes, soluble agonist-antagonist and coreceptor elements to construct cell-cell communication networks [56, 57]. To identify significant signaling sources, targets and mediators, we generated a centrality heatmap which computes centrality metrics from graph theory as used in social network analysis, described previously [58, 59].

#### Statistics and Reproducibility

Statistical approaches and sample sizes for each experiment are indicated in the corresponding figure legends. Unless noted otherwise, experiments were performed with at least three independent replicates for quantitative analysis. For immunofluorescence (IF) experiments, a minimum of three biological samples were quantified, with multiple measurements taken across each tissue. No data points were excluded from any analyses. We assumed normal data distributions but did not perform formal normality testing. Sample sizes were not determined by formal statistical power calculations; however, they are comparable to those used in prior studies^17^. Humane endpoints for animal studies were defined by an investigator blinded to group assignments, although data collection and subsequent analysis were conducted unblinded. For IF and GFP-burden quantifications, image acquisition and analysis parameters were held constant between control and experimental groups to minimize technical bias. Representative IF images were selected based on average values. P value was generated using two-tailed unpaired t-test with Welch’s correction, center line represents mean, and error bars represent standard deviation. For all tests that includes three groups, the statistical analysis was performed as two separate pairwise comparisons using two-tailed unpaired t-tests with Welch’s correction, rather than as a single three-group comparison. Statistical analyses were performed using GraphPad Prism 8. Survival curves were plotted in GraphPad Prism 8 and compared using the Log-rank (Mantel–Cox) test.

## Code Availability

No custom code was used to generate or process the data.

## Availability of materials

Please contact Michael D. Taylor or Xiaochong Wu

## Genotyping Primers and Protocols Table

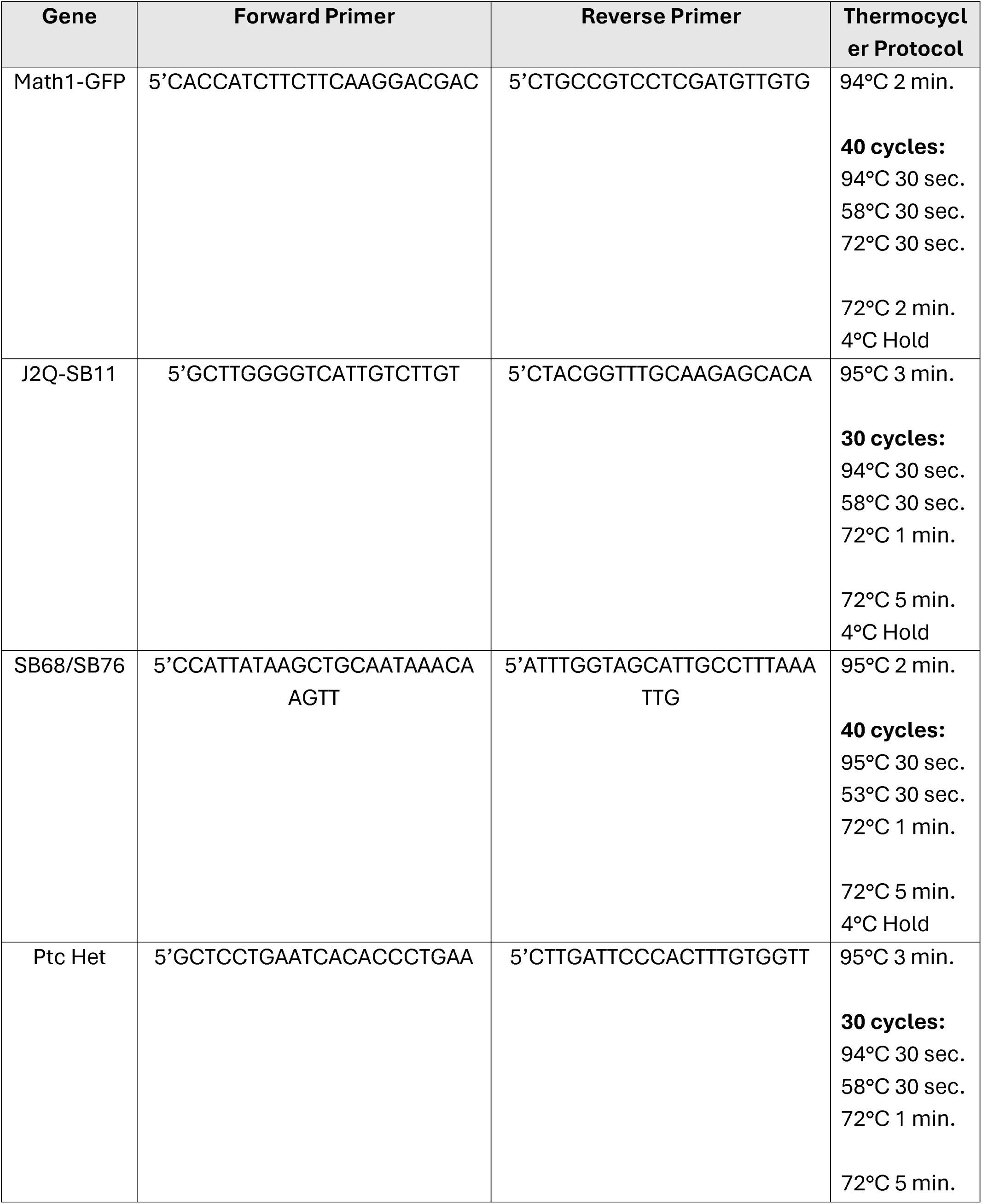
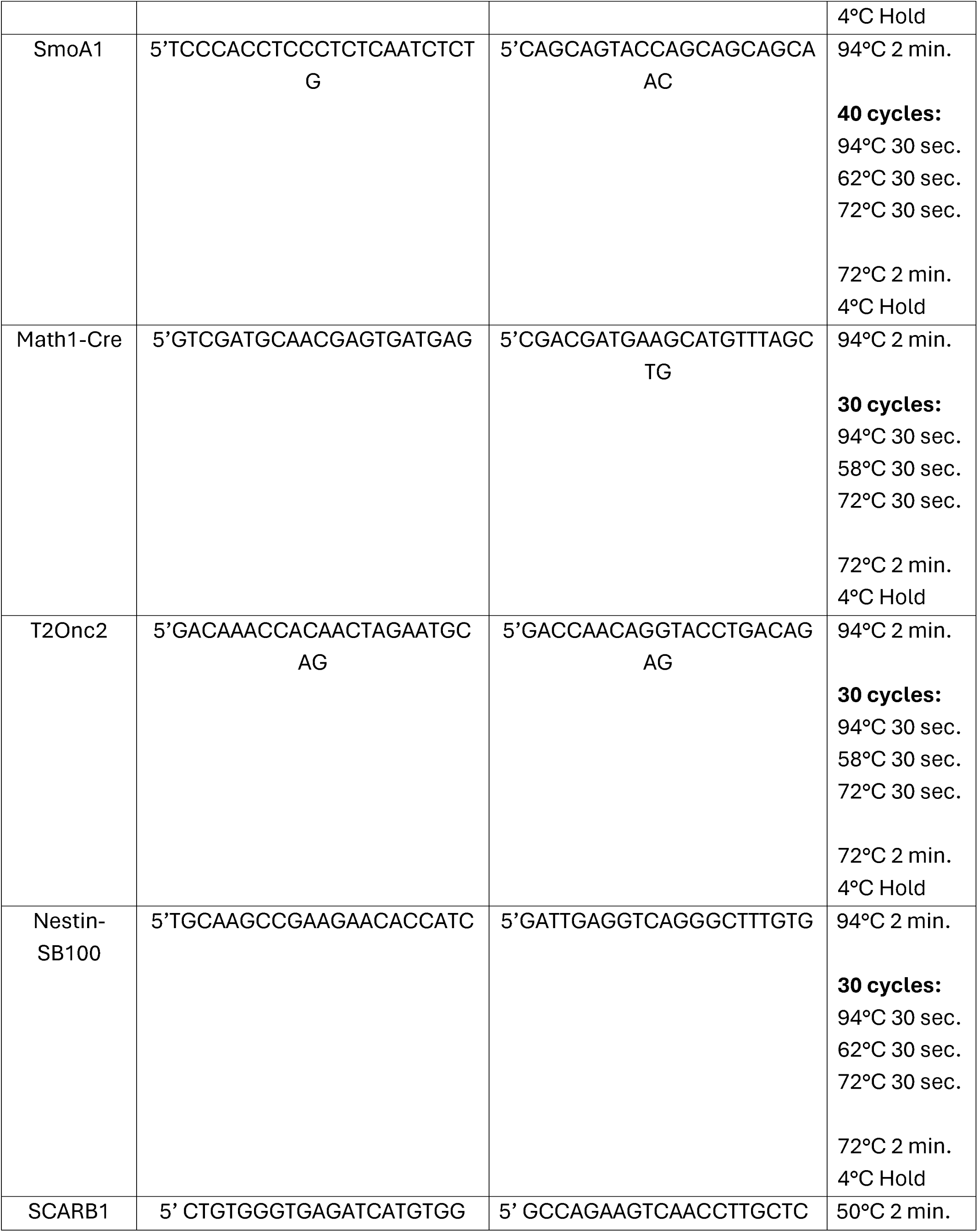
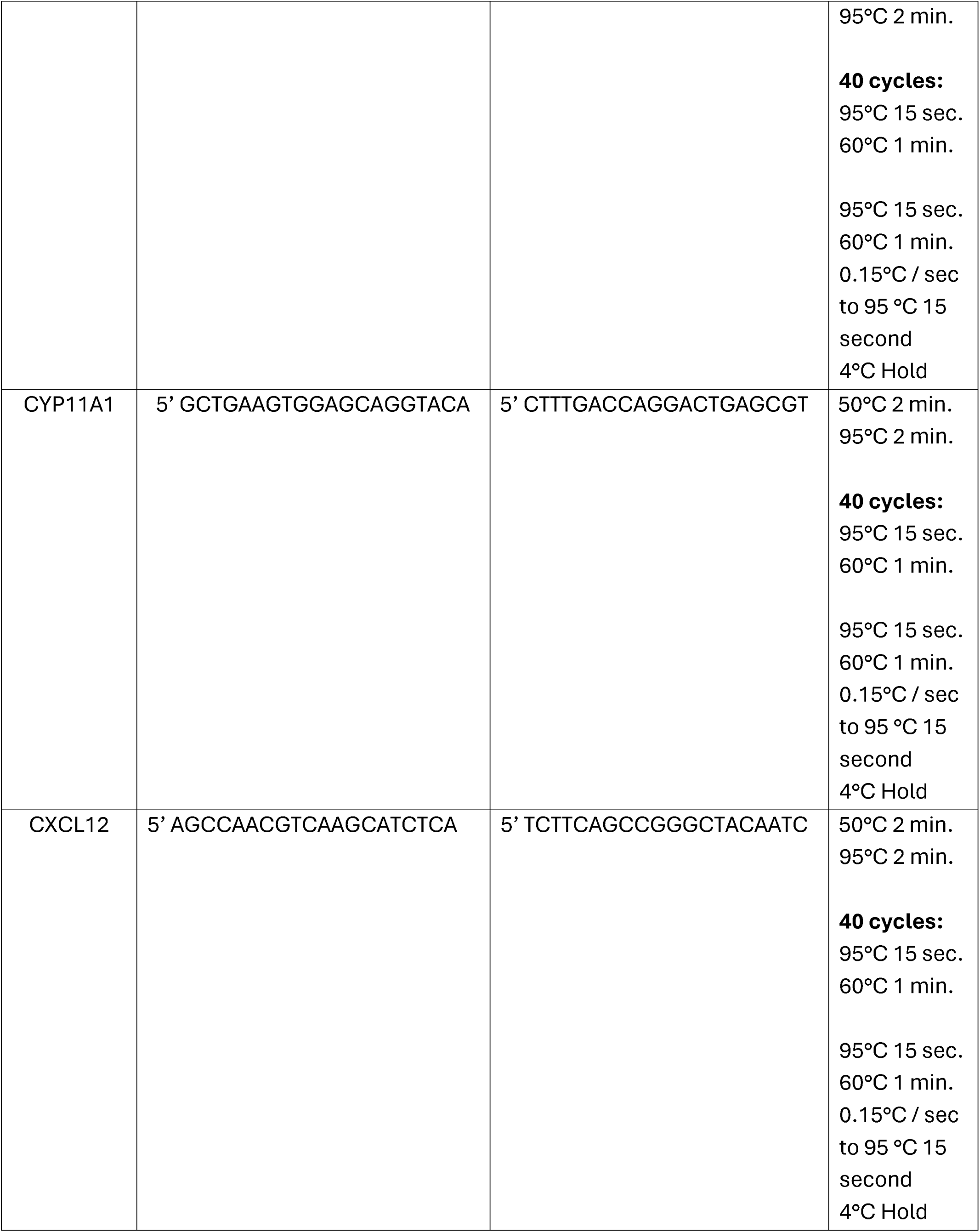

## SUPPLEMENTAL INFORMATIONS

**Supplemental Table 1.** Differentially expressed genes (SMART-seq) between leptomeningeal micrometastases and macrometastases, with the p_val, avg_log2FC, p_val_adj, and neg_log_p listed as columns. Related to Figure 3.

**Supplemental Table 2.** Lipidomic dataset for leptomeningeal micrometastases and macrometastases, with molecular species, lipid class, lipid subclass, features (m/z@rt), and peak area ratio listed as columns. Related to Figure 4.

**Supplemental Table 3.** Differentially expressed genes (Bulk-seq) between shControl D425 and shSCARB1 D425 primary tumors, with the log2FoldChange, pvalue, and padj listed as columns. Related to Figure 5.

**Supplemental Table 4.** Differentially expressed genes (SMART-seq) between SLP*-*mCherry + macrophages and SLP*-*mCherry - Macrophages, with the p_val, avg_log2FC, p_val_adj, and neg_log_p listed as columns. Related to Figure 6.

