## Supplementary figures and images for "Medulloblastoma Forms Symbiotic Metabolic Partnerships with Macrophages to Establish Leptomeningeal Metastases"

### Supplemental Figure 1

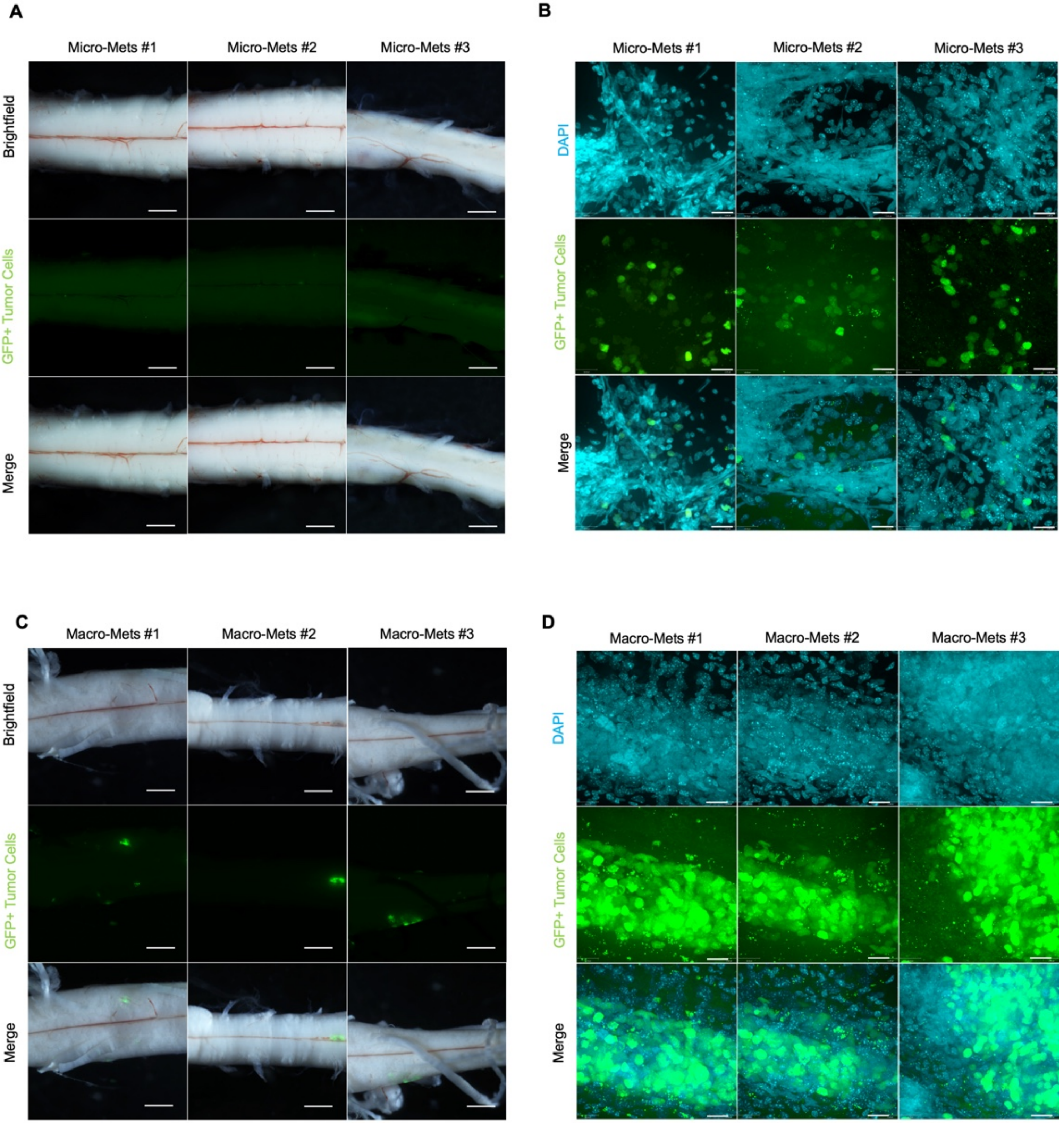

### Supplemental Figure 3

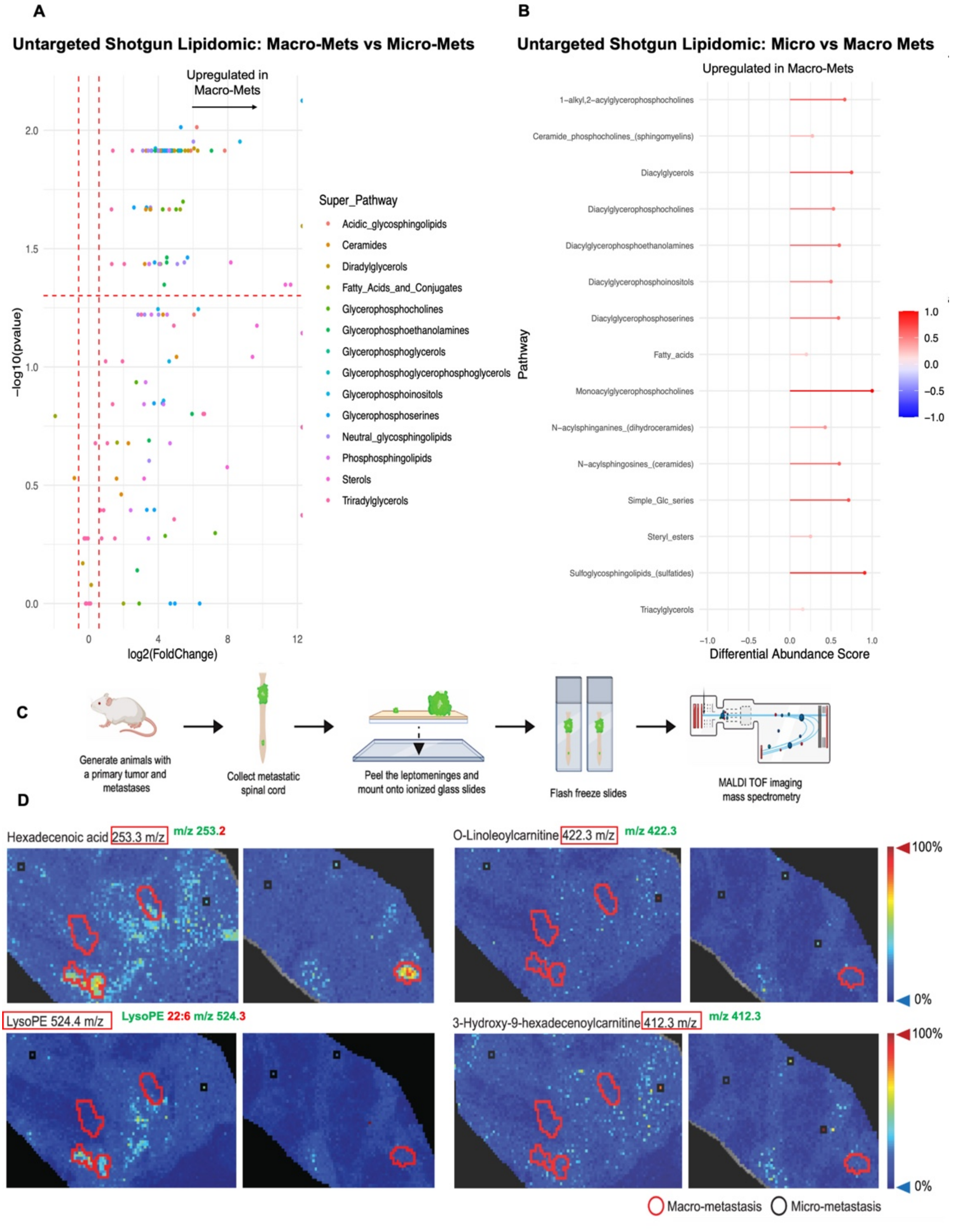

### Supplemental Figure 4

**Figure S4**

**A** Spinal Meningeal Filipin Staining

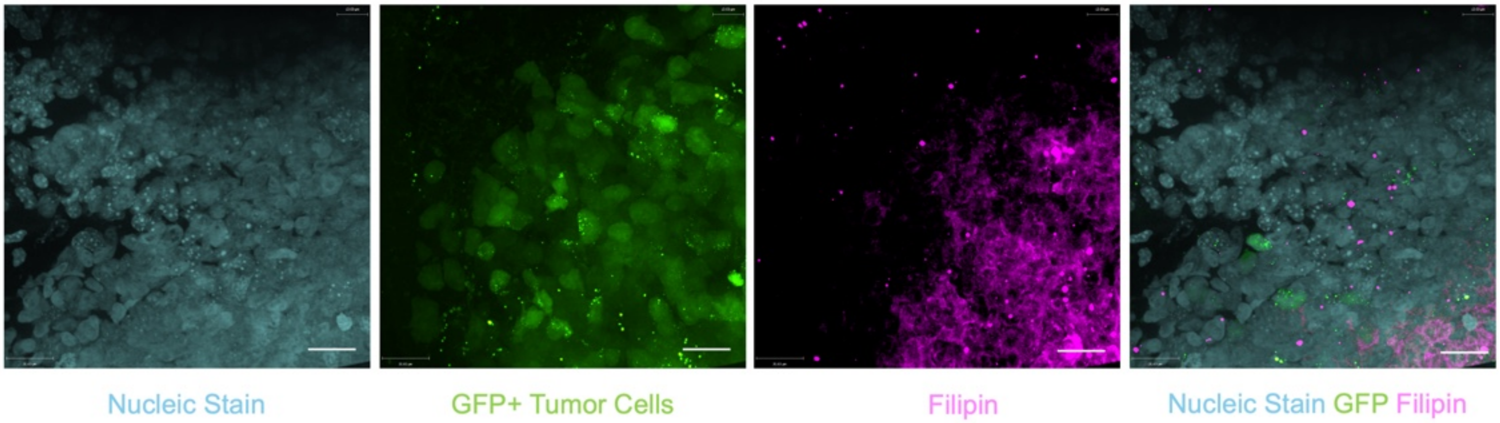


**B**

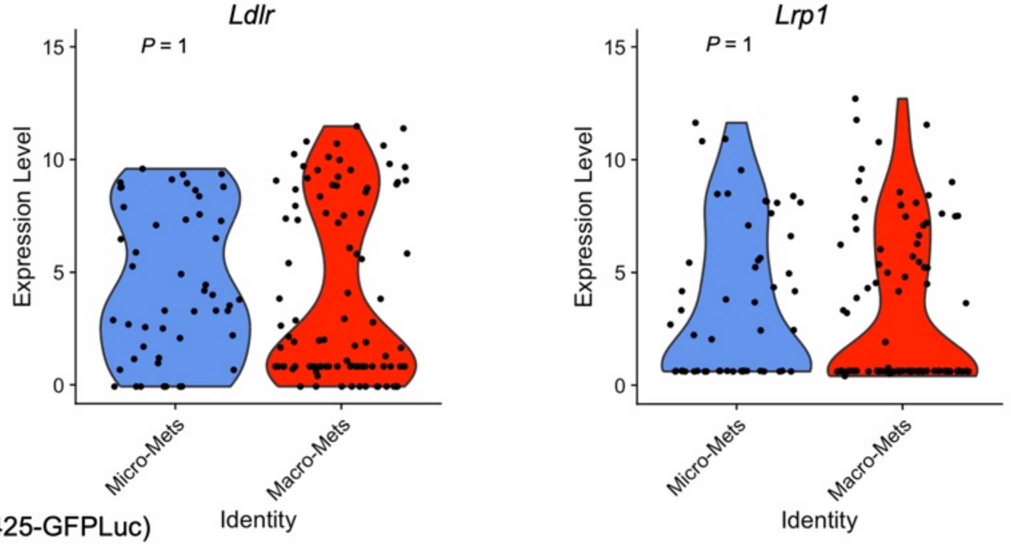


**C**

Primary Tumor (D425-GFPLuc)

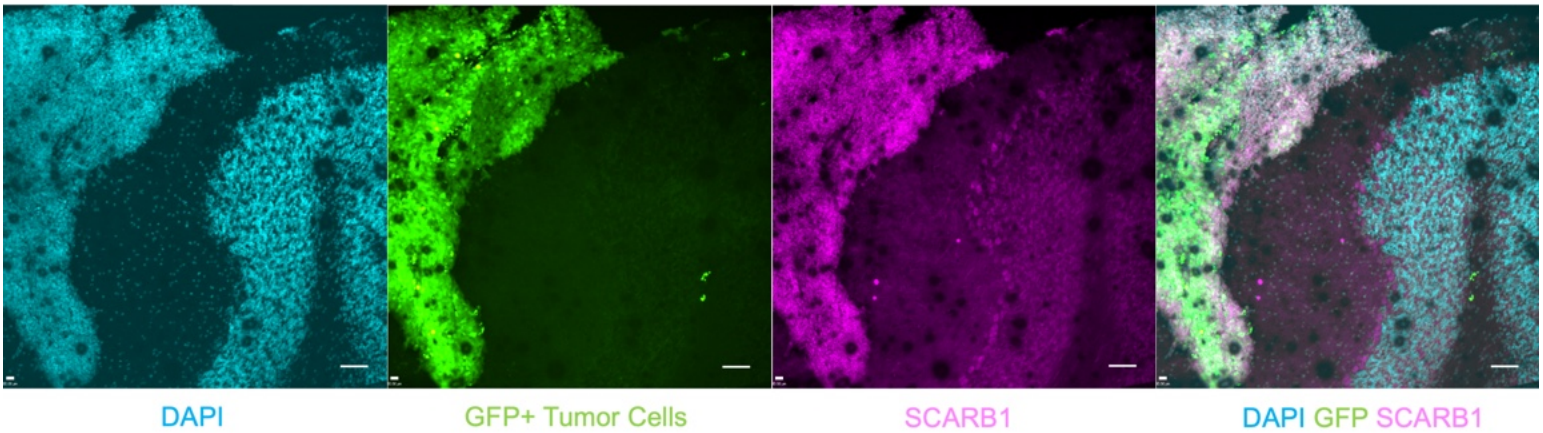


**D**

shSCARB1 Western Blot

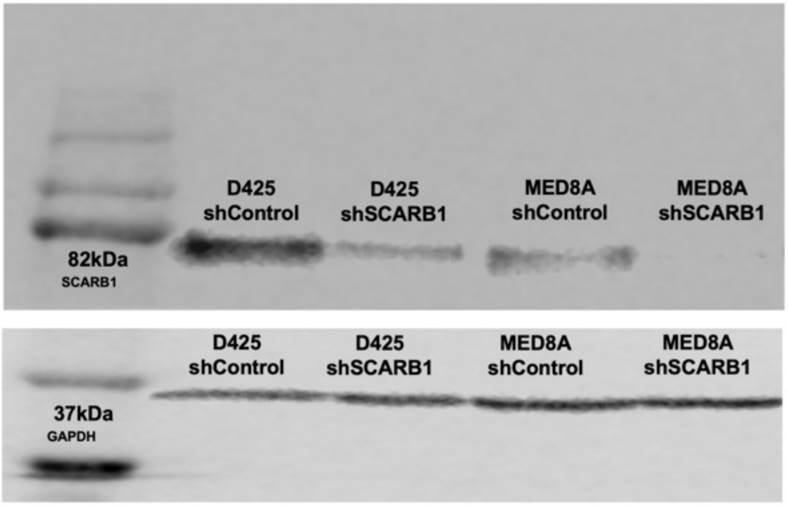

### Supplemental Figure 5

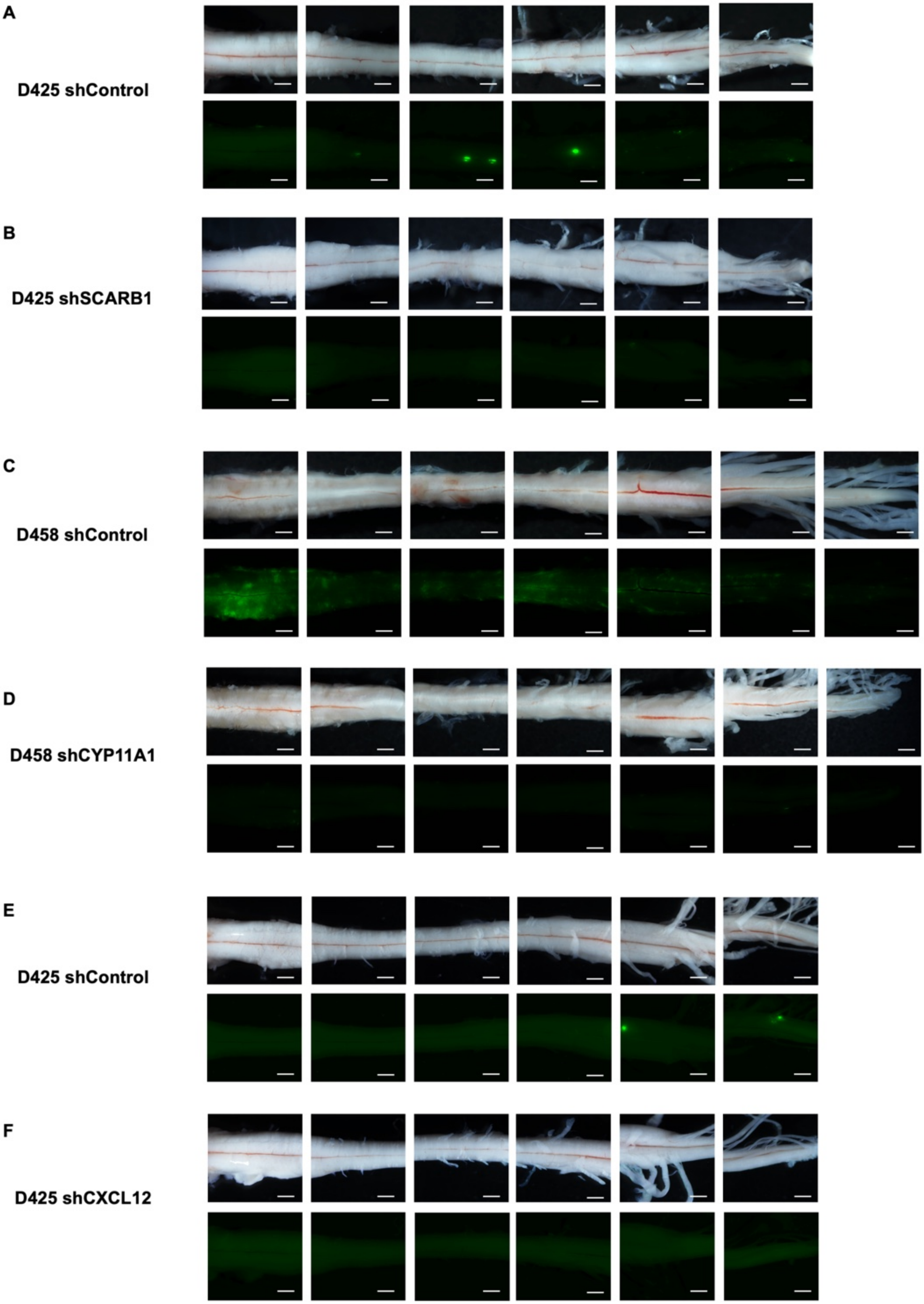

### Supplemental Figure 6

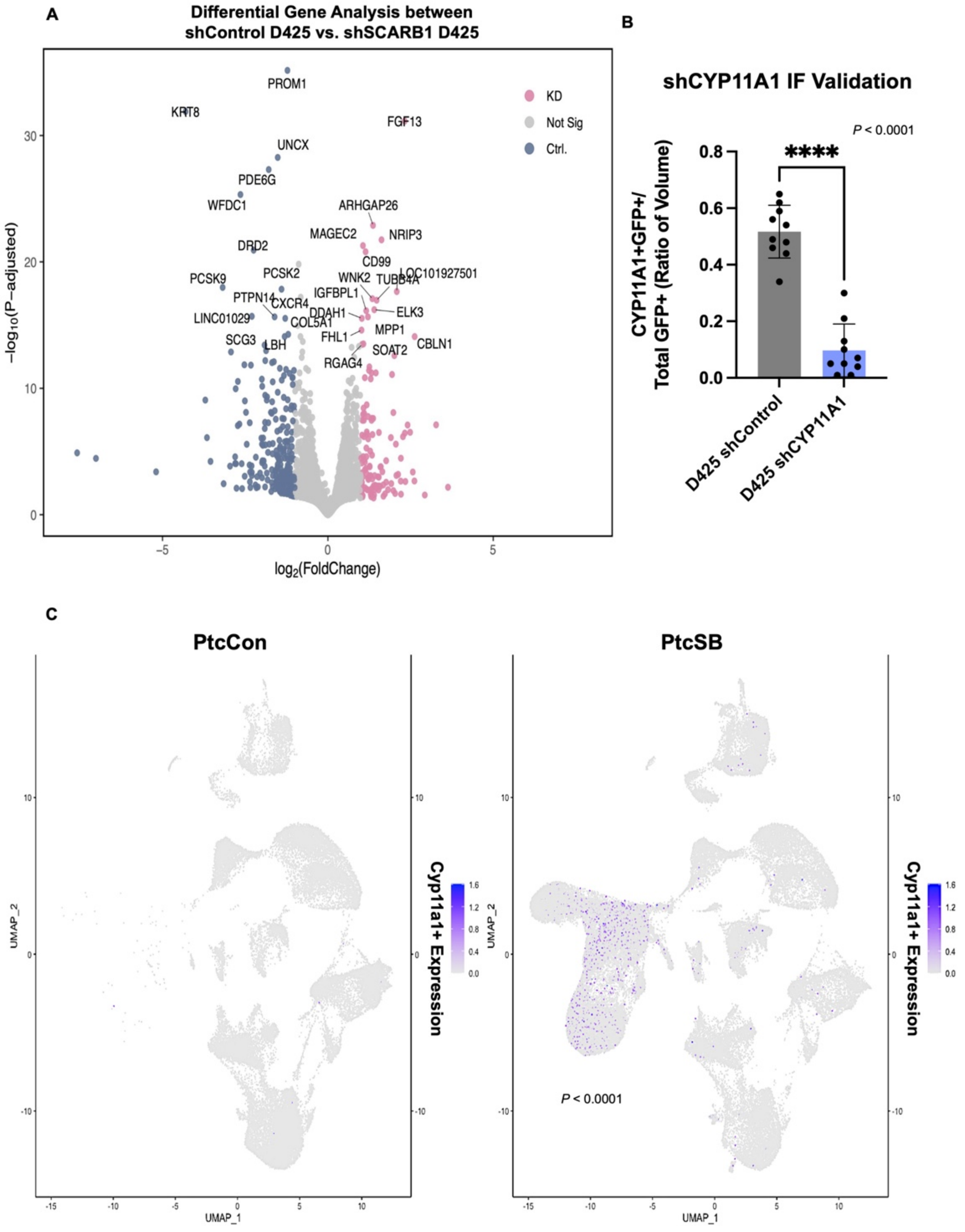

### Supplemental Figure 8

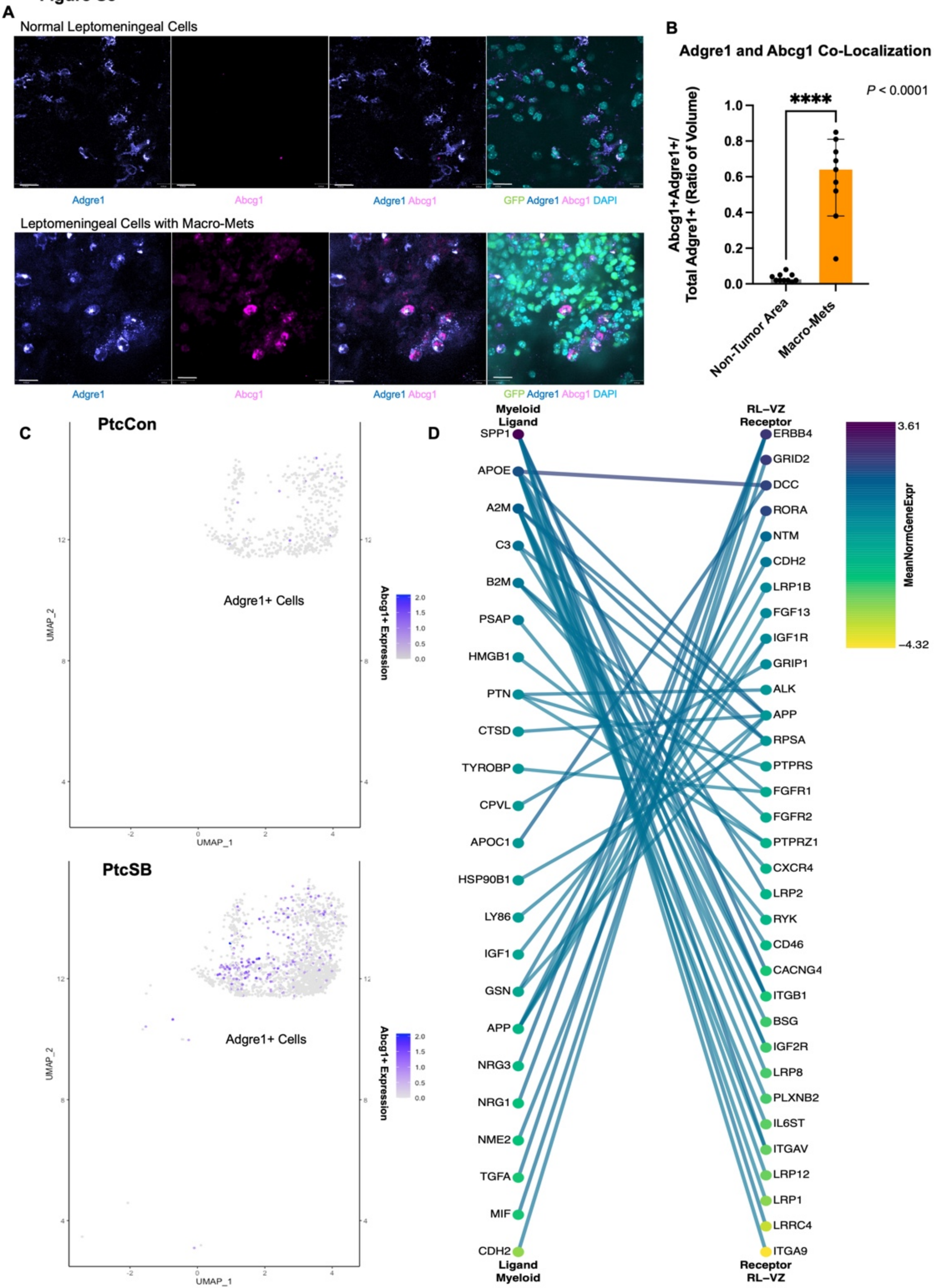
