## Supplemental Figure 2 for "Medulloblastoma Forms Symbiotic Metabolic Partnerships with Macrophages to Establish Leptomeningeal Metastases"

### Figure S2

#### A In vivo Glucose Tracing Results

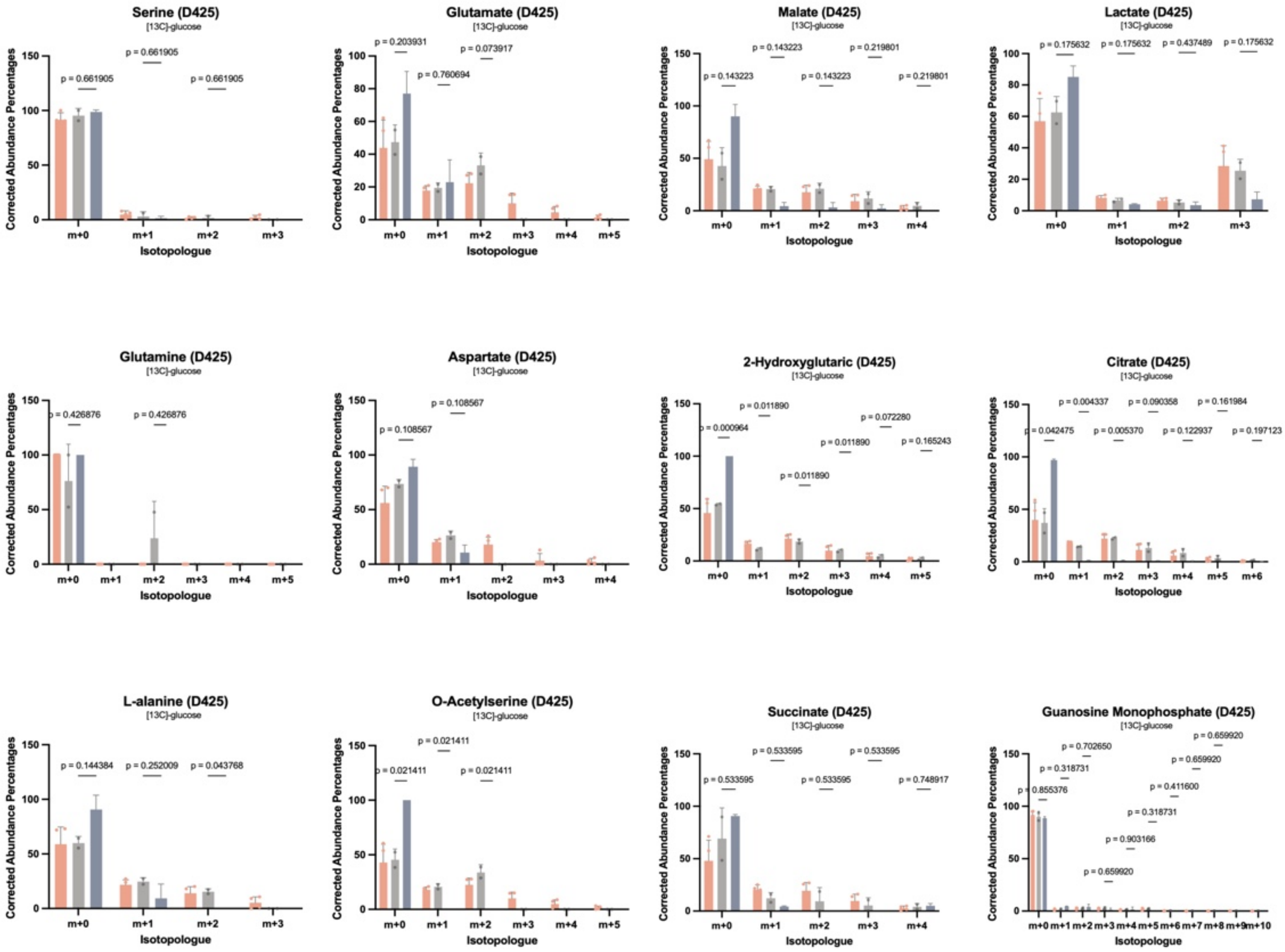


#### B Enriched pathways in fatty acid/lipid metabolism from Micro/Macro Mets SMART SEQ

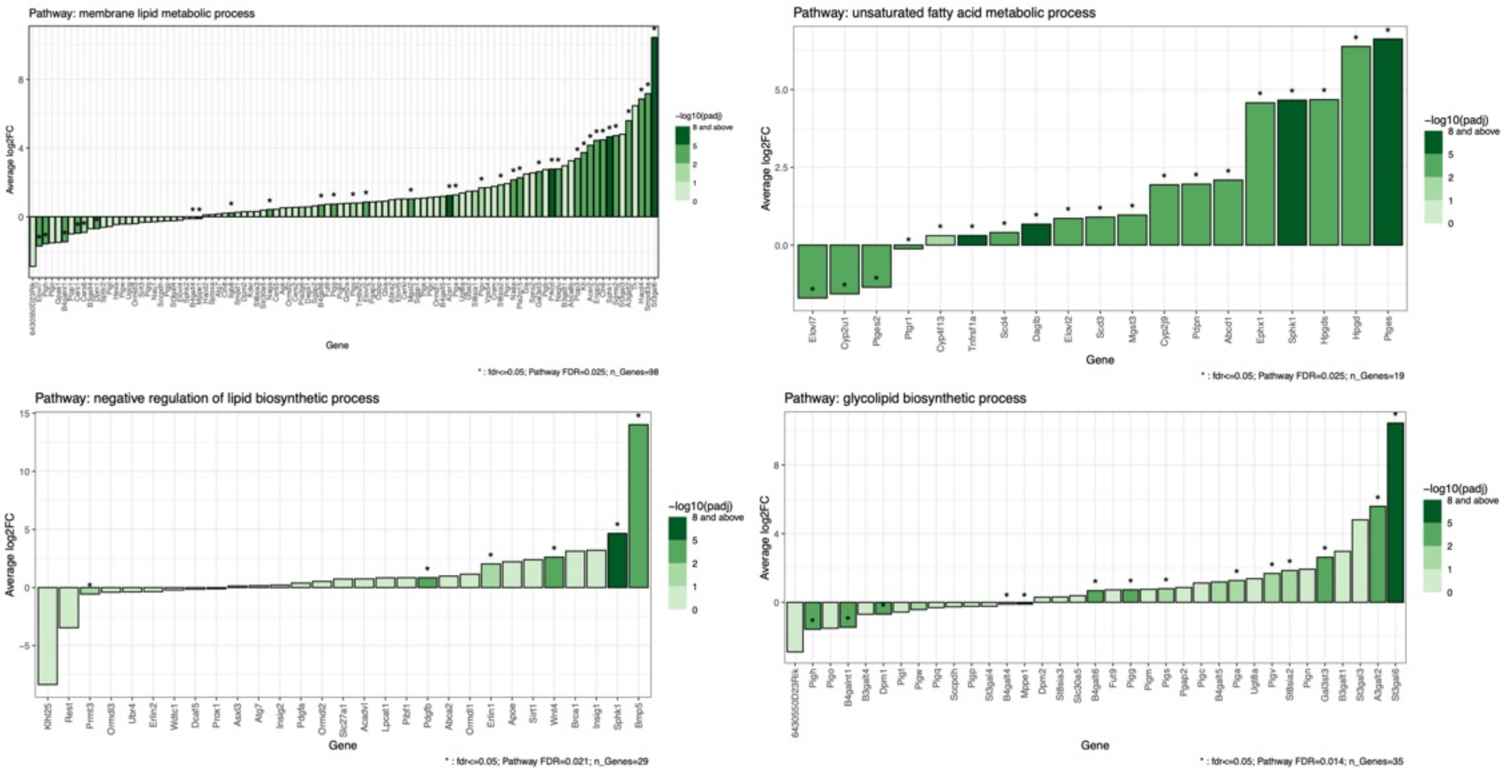
