## Supplemental Figure 7 for "Medulloblastoma Forms Symbiotic Metabolic Partnerships with Macrophages to Establish Leptomeningeal Metastases"

Figure S7

A

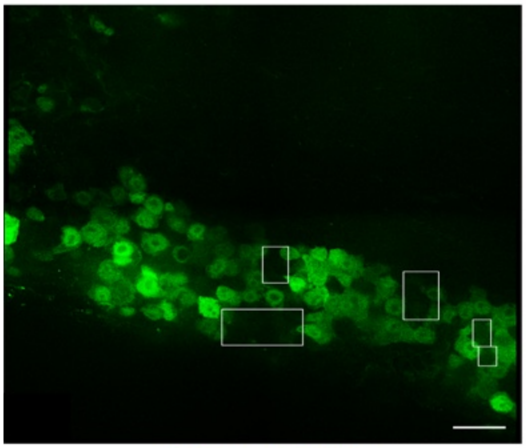

GFP+ Tumor Cells

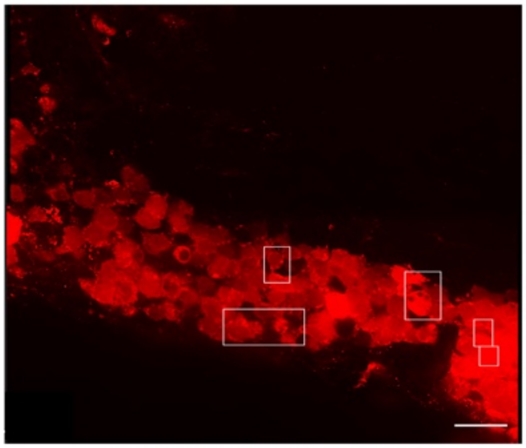

sLP-mCherry Expression

B

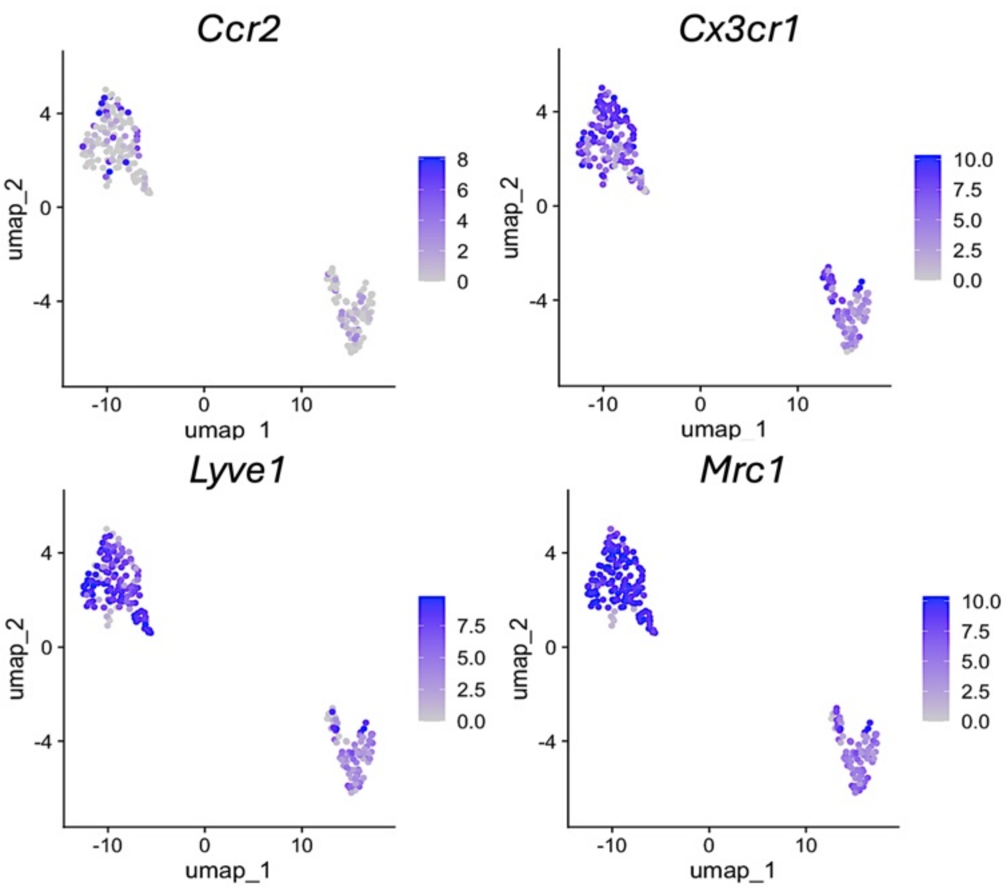


C

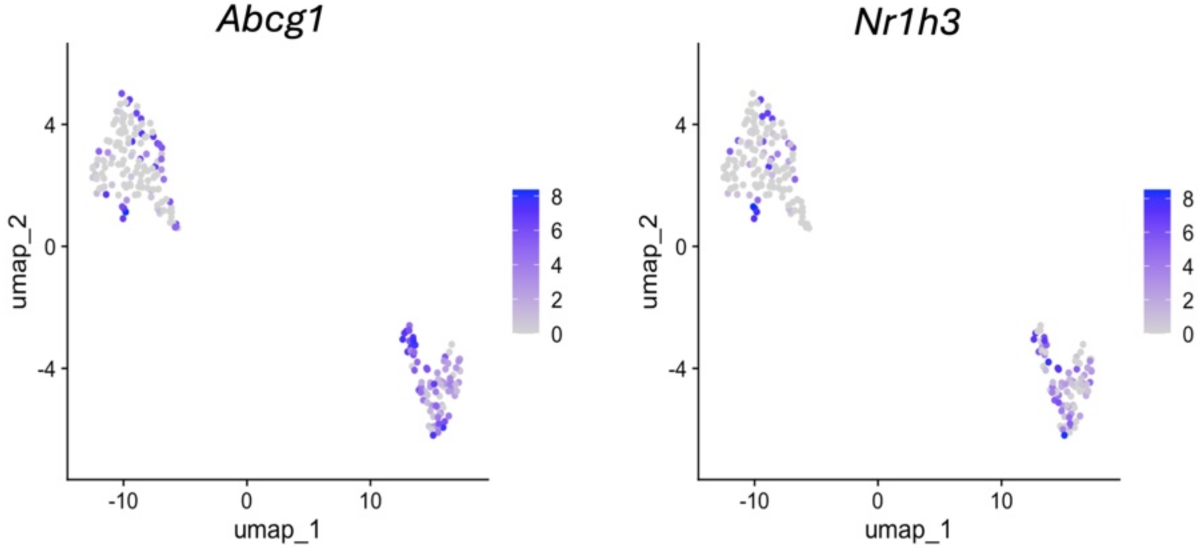


D

Primary Tumor (D425-GFPLuc)

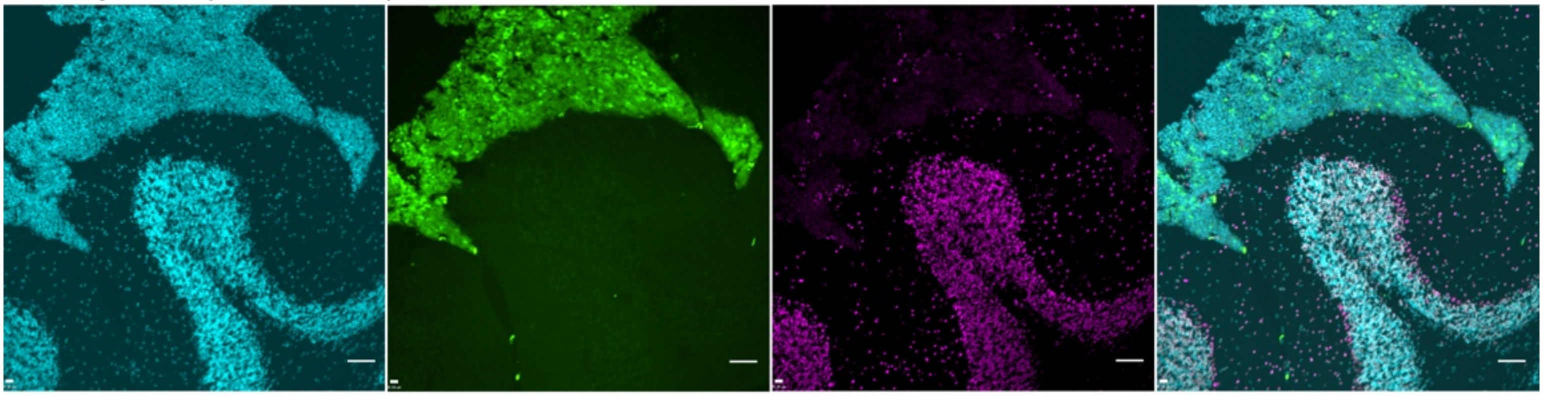

DAPI GFP+ Tumor Cells ABCG1 DAPI GFP ABCG1
