## Supplemental Figure 9 for "Medulloblastoma Forms Symbiotic Metabolic Partnerships with Macrophages to Establish Leptomeningeal Metastases"

**Figure S9**

**A**

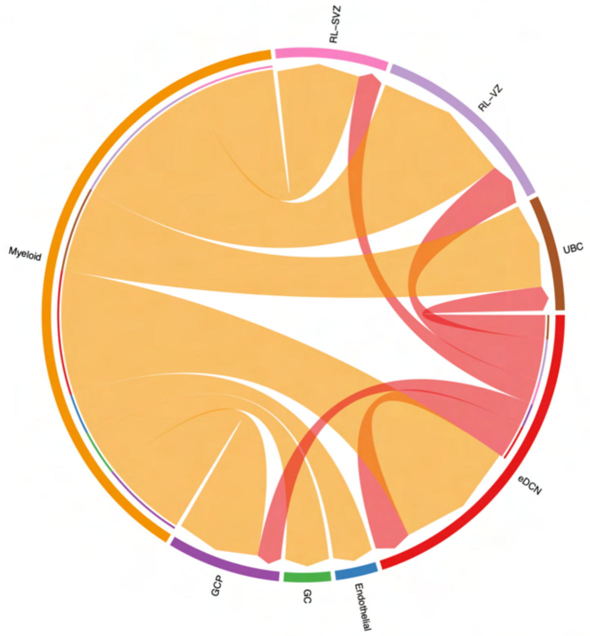


**B**

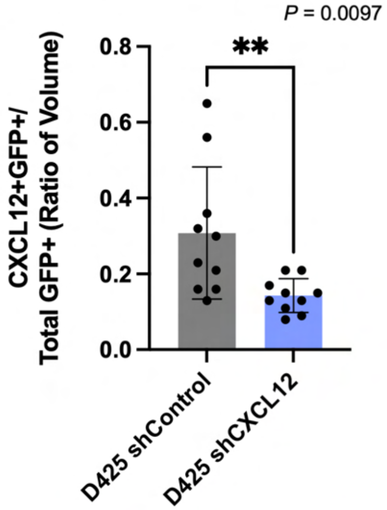


**C**

**shControl D425 Leptomeningeal Tumor**

**Image 1**

**Image 2**

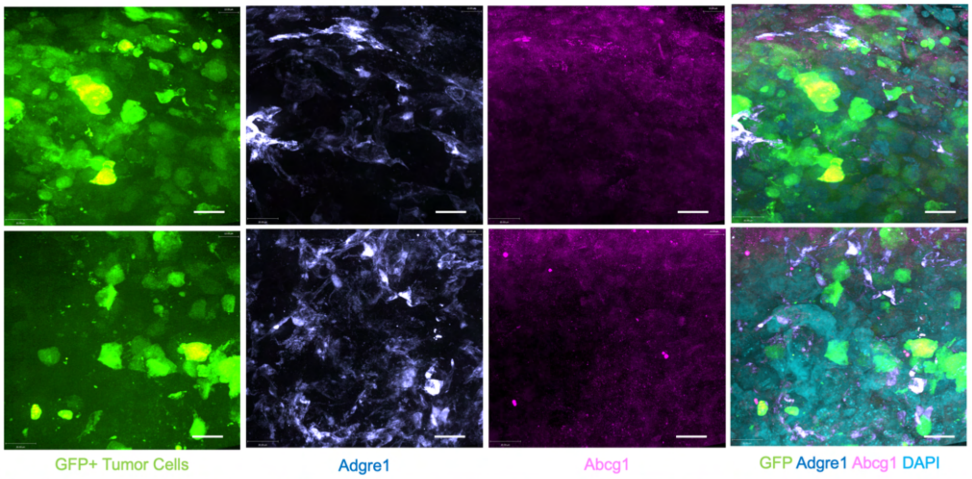


**D**

**shCXCL12 D425 Leptomeningeal Tumor**

**Image 1**

**Image 2**

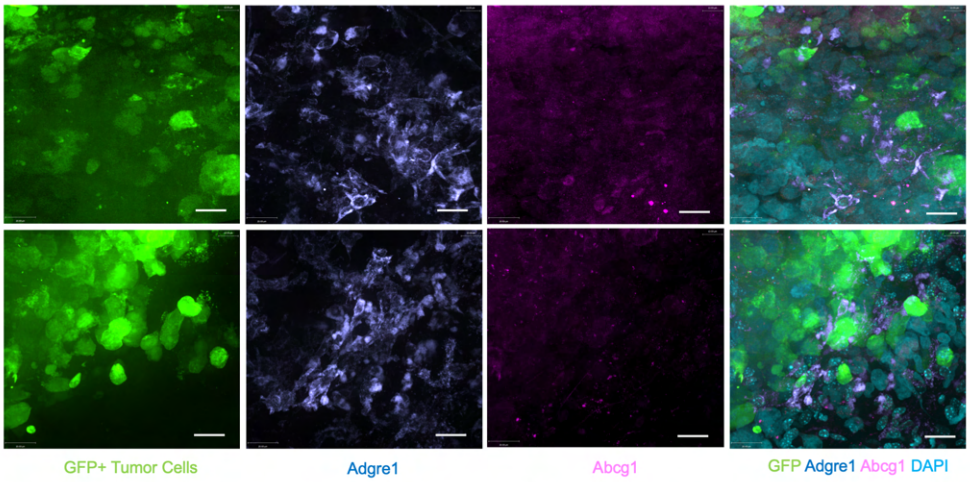


**E**

**Percent Nutrient Compositions of Control and High Fat Diet**

| | Control Diet | High Fat Diet |
|---|---|---|
| Protein % | 18.6 | 19.0 |
| Fat % | 6.2 | 9.0 |
| Crude Fiber % | 3.5 | 2.6 |
| NDF %[1] | 14.7 | 12.1 |
| Metabolizable energy | 3.1 kcal/g<br>13.0 kJ/g | 3.3 kcal/g<br>13.8 kJ/g |
